# Longevity hinders evolutionary rescue through slower growth but not necessarily slower adaptation

**DOI:** 10.1101/2024.04.11.588938

**Authors:** Scott W. Nordstrom, Brett A. Melbourne

**Affiliations:** Department of Ecology and Evolutionary Biology, University of Colorado, Boulder, Boulder, CO, USA, 80309; Population Research Center, Portland State University, Portland, OR, USA, 97214

**Keywords:** Life history, viability selection, frailty effect, longevity

## Abstract

“Evolutionary rescue” is the process by which a population experiencing severe environmental change avoids extinction through adaptation. Theory and empirical work typically focus on short life histories with non-overlapping generations, leaving longevity’s effects on rescue relatively understudied. Recent models demonstrate that longevity can inhibit rescue through slower phenotypic evolution but have assumptions that may not generalize across life histories. We built a model integrating evolutionary rescue with concepts from life-history theory, particularly the fast-slow pace-of-life continuum. Longevity is modeled by the balance of survival and reproduction with selection acting on survival, allowing for multiple selection episodes throughout the lifespan. We used this model to simulate three life-history strategies along the fast-slow continuum responding to sudden environmental change. Under nearly all simulated conditions, higher longevities (slower pace of life) resulted in more time at low density and increased extinctions. With perfect trait heritability, rates of adaptation were nearly identical across longevities. But at lower heritabilities, longevity allowed for repeated selection and decoupling of mean genotypes and phenotypes, producing a transient phase of rapid phenotypic change. Our results demonstrate that prior findings that longevity slows adaptation do not hold in all cases and are relevant to long-lived conservation targets.

## Introduction

Evolutionary rescue, the phenomenon of populations avoiding extinction following environmental change by adapting to novel conditions, has emerged as an important concept in recent decades due to climate change and widespread habitat alteration and degradation. This includes both theoretical (Carlson et al. (2014), Kopp & Matuszewski (2014), Klausmeier et al. (2020)) and empirical work (Bell & Gonzalez (2009), Agashe et al. (2011), Hufbauer et al. (2015), Clark-Wolf et al. (2024)), with considerable progress in generating and testing predictions about which factors influence population persistence following severe environmental change. At present, many rescue models assume or impose non-overlapping generations of short, fixed length. While these assumptions increase analytical tractability and allow for validation with model systems where life history can be controlled in the laboratory (e.g., algae [Bell & Gonzalez (2009), Bell (2012)], flour beetles [Agashe et al. (2011), Hufbauer et al. (2015)], and *Daphnia* [Loria et al. (2022)]), they do not reflect the life history of many organisms. This is particularly true for large-bodied organisms with long, complex life cycles and multiple reproductive episodes over the lifetime that are likely to be of conservation concern (Purvis et al. (2000), Dirzo et al. (2014)).

It has been hypothesized that longevity hinders evolutionary rescue, primarily by slowing rates of phenotypic adaptation. One such posited mechanism is reduced generational turnover due to longer generation times, slowing absolute rates of phenotypic change (e.g., Chapin et al. (1993), Vander Wal et al. (2013), Carlson et al. (2014)). This hypothesis has been supported by models of boreal trees (Kuparinen et al. (2010)) and alpine plants (Cotto et al. (2017)). However, both of these models assumed selection acted only once in a lifespan (on the viability of new recruits). This assumption may limit the applicability of these results to populations where individuals are subject to multiple selection events, e.g., where annual survival is influenced by body size or conditions (e.g., Milner et al. (1999), Ozgul et al. (2010), Clark-Wolf et al. (2024)). A related hypothesized mechanism for longevity’s influence on rate of adaptation and probability of rescue is that longevity may provide for life cycle “buffering” where certain life stages are shielded from selection pressures (e.g., Shoemaker & Lennon (2018), Schmid et al. (2022)). This hypothesis was supported by theoretical findings of Schmid et al. (2022), but this model likewise assumed selection acted once throughout the life cycle and thus may not generalize to long-lived organisms that experience multiple selection events over their lifespan. Additionally, one may argue that life cycle buffering is not an inherent feature of longevity, as many organisms that are considered “annual” or short-lived in their somatic form have dormant life stages that are buffered from selection pressures, such as annual plants with seed banks (Templeton & Levin (1979)), copepods (Hairston & de Stasio (1988)), and even microbes (Shoemaker & Lennon (2018)). Theory has investigated how selection that reduces survival rather than births can reduce generation times as populations become maladapted, hastening rates of adaptation (Draghi et al. (2024)); however, this analysis did not consider variation in underlying generation times or life histories. Thus, while longevity can impede rescue by slowing phenotypic adaptation in some circumstances, a comprehensive understanding of the effects of longevity on rescue involves considering additional evolutionary and demographic patterns that are associated with longer-lived life cycles (Gadgil & Bossert (1970), Jones et al. (2014)).

Consideration of rescue and longevity that focuses primarily on adaptive dynamics between generations ignores dynamics within generations. For example, survival- or growth-related traits such as body size (Milner et al. (1999), Clark-Wolf et al. (2024)) are potentially subject to multiple rounds of selection as individuals age; longer generation times thus may increase the lifetime selection pressure an individual may face. Coulson & Tuljapurkar (2008) demonstrated, in an extension of the Price equation (Price (1970)) how trait distributions in age-structured populations change not only with differential number of offspring produced by parental phenotypes (“fertility selection”) but also with differential survival among phenotypes (“viability selection”), and showed that inter-annual changes to birth weight in a population of red deer were driven more by viability selection than fertility selection. Repeated selection on traits can also produce a decoupling of mean population phenotypes and genotypes (Arnold & Wade (1984), Cotto & Chevin (2020)). Phenotypic-genotypic decoupling, when arising due to clonality rather than repeated selection, can hasten phenotypic change and slow genotypic change, ultimately contributing positively to evolutionary rescue (Orive et al. (2017)). However, these evolutionary dynamics that may be associated with longevity are typically not explored in the context of evolutionary rescue; in particular, their interactions with demography and population dynamics are necessary for considering the role of evolution in avoiding extinction following environmental change.

Whereas considerations of longevity in evolutionary rescue have primarily focused on rates of adaptation, life-history theory has several additional and important insights into the demographic and ecological consequences of longevity. One important, well-supported finding is that longevity produces a trade-off between stability and population- or lineage-level rates of increase. For example, *r*-vs-*K* selection hypothesizes that certain species are “*r*-selected,” typically having shorter lifespans and investing more reproduction that allows populations to achieve rapid growth at low density (Pianka (1970)). In contrast, “*K*-selected” species invest more in survival, longevity, and competitive ability, stabilizing population growth rates in harsh environments but slowing intrinsic population growth rates (Pianka (1970)). This trade-off is similarly apparent in the “fast-slow” pace-of-life continuum, where species closer to the “fast” end of the continuum are shorter lived and have higher low-density population growth rates compared to species on the “slow” end (Oli (2004), Salguero-Gómez et al. (2016)). Thus, while high survival that produces longevity can be advantageous for stabilizing population size in harsh or fluctuating environments or when populations are at low density, a population comprising these longer-lived individuals will take longer to return to original size following a dramatic reduction in size due to its constrained growth rate (Pimm et al. (1988)). Despite the incredible diversity of life histories and demographic patterns across the tree of life (Jones et al. (2014)), the fast-slow continuum explains a considerable amount of variation in life-history traits and population growth rates in animals (Gaillard et al. (2005)) and plants (Salguero-Gómez et al. (2016)) such that there are relatively few species that are long lived with very rapid population growth or short lived but with constrained maximum growth rates. For this reason, consideration of the effects of longevity on evolutionary rescue should incorporate demographic characteristics associated with longer-lived, “slow” paces of life, including lower per-bout fecundity, higher survival, and constrained population growth rates. This is particularly true considering many of these traits are present in larger-bodied, longer-lived species of particular conservation concern (Purvis et al. (2000), Denney et al. (2002), Cardillo (2003), Dirzo et al. (2014)).

We thus move towards a synthesis of evolutionary rescue and life-history theory to answer questions about effects of longevity on rescue. We build a simple demographic and evolutionary model that allows for variable longevity among life-history groups. We impose the assumption that high longevity (slower pace of life) corresponds to lower intrinsic growth rates because this assumption is well supported in empirical life-history literature. Our model features two vital rates, fecundity and survival, with selection acting on survival through a quantitative trait in each time step, thus allowing for repeated bouts of selection throughout the lifespan. Using this model and accompanying simulations, we address how pace of life influences population size, rates of phenotypic and genotypic change, and age structure over time. Because the decoupling of phenotypes and genotypes that can occur with repeated selection relies on imperfect heritability, we also consider the interactions of longevity with heritability and degree of phenotypic variance to influence rescue outcomes.

## Methods

We present a discrete-time stochastic simulation model, providing accompanying analytical expectations for the simulation model where possible. All simulations were performed in R version 4.4.1 (R Core Team (2020)). Simulations were individual-based, allowing us to explicitly model extinctions, which occur when a population has zero surviving individuals. Upon birth or initialization individuals are assigned a sex, male or female, with equal probability. We include sex in our simulations because it is present in many animal species and because stochasticity in sex ratios in small populations contributes to extinction risk (Melbourne & Hastings (2008)). Each time step features mortality followed by reproduction, modeled as follows. Individuals survive with probability *s*_*i*_ (where *i* indexes individuals) as determined by a quantitative trait as described in the paragraph below. Each individual is subject to selection in each time step for which it is alive, allowing for multiple selection episodes over the lifespan. This reflects selection for survival-related traits such as body size or condition (Milner et al. (1999), Ozgul et al. (2010), Clark-Wolf et al. (2024)). After selection, each surviving female is paired with a randomly-chosen surviving male and produces a Poisson-distributed number of offspring with mean 2*r*. Our analytical results assume that mating is panmictic and that populations have equal sex ratios such that mean fecundity is *r*. Both for simplicity and to distill the effects of repeated bouts of selection on survival, we do not assume selection on fecundity, leaving this for future work. Thus, if 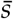 is the mean survival probability in the population at the beginning of the time step, the expected population growth rate is 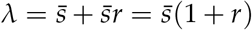.

An individual’s trait (phenotype) takes a continuous value *ζ*_*i*_, which measures the difference between the individual’s trait value and the environmental optimum, i.e., fitness is maximized by *ζ*_*i*_ = 0. Traits are subject to Gaussian selection with a selection surface of width *ω* such that an individual with trait *ζ*_*i*_ has survival proportional to exp 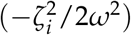 (Lande (1976)). For simplicity, we assume that selection operates with the same strength across the lifespan, i.e., *ω* is independent of age (see Cotto & Chevin (2020) for a similar model where selection strength may vary with age). This allows us to scale phenotypes and their respective components by the width of the selection surface; define the scaled phenotype as *z*_*i*_ = *ζ*_*i*_/*ω*. If *σ*^2^ denotes the variance of unscaled phenotypes, then the variance of scaled phenotypes is *γ*^2^ = *σ*^2^/*ω*^2^. Increasing *γ*^2^ corresponds to increasing phenotypic variance and/or stronger selection. We will focus our analysis on scaled phenotypes (*z*) rather than un-scaled phenotypes (*ζ*) for ease of notation. The phenotype of individual *i* can be decomposed into the sum of a genetic contribution (henceforth, “breeding value”), *b*_*i*_, and a non-inherited contribution, (henceforth, “environmental component”), *e*_*i*_ (Falconer & Mackay (1996)). We denote the variance in breeding values within a population as 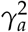 and the variance of environmental components as 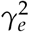. Analytical results assume random mating in the population, such that 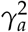 is also the additive genetic variance for the trait in the population (Lynch & Walsh (1998)). For an individual with phenotype *z*_*i*_, probability of survival in one time step is 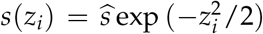, where 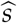 is the per time step maximum rate of survival, i.e., the survival probability of an individual perfectly adapted to the environment. In a constant environment, the intrinsic fitness (expected lifetime number of offspring) of an individual with survival *s*_*i*_ is *W*_*i*_ = *s*_*i*_*r*/(1 − *s*_*i*_), since the number of time steps individual *i* lives for is geometrically distributed with rate *s*_*i*_ and expected reproductive output is *r* in each time step.

### Life-history strategies

We denote the maximum absolute intrinsic fitness, i.e., the expected number of offspring per individual perfectly adapted to its environment, as 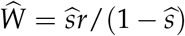. Henceforth, we will hold 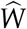 constant among life-history strategies, modifying 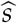 and *r* to define different strategies (e.g., large 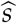 and small *r* corresponds to a high-longevity, slow pace-of-life strategy). Thus, for a given life history with fixed survival 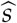, the corresponding fecundity is 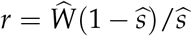.

For a given 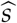 and *r*, the maximum expected population growth rate per time step is 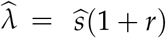. Rewriting *r* in terms of 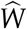 produces 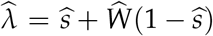, which has a monotonically decreasing relationship with 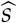. As 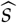 approaches zero (the case of semelparity with non-overlapping generations), 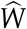 approache s 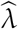 (a lthough when 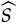 is exactly zero, 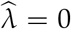). As 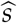 approaches one, 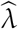 also approaches one. This produces the widely-observed trade-off where populations of longer-lived organisms have lower intrinsic growth rates (Pianka (1970), Pimm et al. (1988), Oli (2004), Salguero-Gómez et al. (2016)).

### Equilibrium dynamics

In this section we analytically characterize the steady-state age distribution and phenotypic variance structure in the population. We assume that the population has mean phenotype 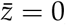, i.e., is adapted to its environment. Our analytical results also assume that the additive genetic variance in the population remains constant over time; this is enforced in simulations as described in subsection “*Phenotypic components and their variances*”.

### Age distribution and population growth

Upon birth, a cohort of newly-born individuals has phenotypes that are normally distributed with variance 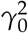. After *k* rounds of Gaussian selection under a constant environment, the phenotypic distribution will remain normally distributed with variance 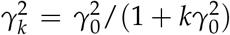 (see Appendix S1). Because the trait is normally distributed, we can use a result of Lande (1976) for the integral of a Gaussian fitness landscape over a trait distribution to obtain 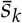, the mean probability that an individual of age *k* will survive to age *k* + 1 :

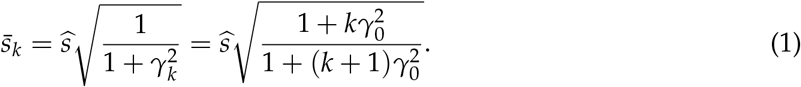

This equation demonstrates that survival within a cohort increases with cohort age, converging on 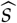 at a rate that increases with 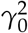. This is an example of the “frailty effect,” wherein survival within a cohort increases over time due to the removal of less-fit individuals (Vaupel et al. (1979)). We demonstrate in Appendix S2 that defining the frailty of an individual with phenotype *z*_*i*_ as 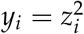 allows several results of Vaupel et al. (1979) to be applied to our model.

Assume that at equilibrium, the population has intrinsic growth rate *λ*^*^ such that 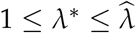. Using the Euler-Lotka equation, the stable age distribution (*p*_*k*_, the probability of a randomly sampled individual being age *k*) can be defined as

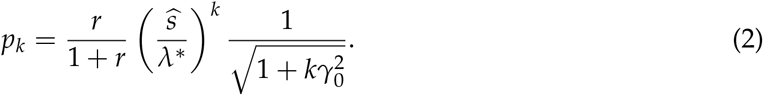

See Appendix S3 for details. An alternative parameterization can be shown by noting that *p*_0_ = *r*/(1 + *r*), in which case 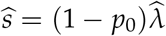 and the age distribution can be expressed as

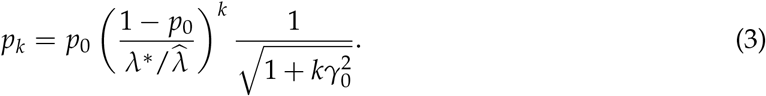

Setting 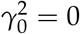, in which case there is zero phenotypic variation and all individuals have survival 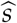 (and thus 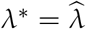), produces a geometric age distribution. Increasing 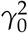 will increase the mass of older cohorts in the distribution due to the frailty effect (Fig. S1). Larger *p*_0_, associated with smaller 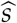, implies a relatively large newborn cohort and therefore a faster pace of life. Likewise, *p*_0_ can be interpreted as proportional to the rate of generational turnover, and 1 − *p*_0_ can similarly be considered a rate of generational overlap (Yamamichi et al. (2019)).

### Phenotypic components and their variances

In simulations, each offspring’s breeding value is generated by taking the mean of its parents’ breeding values and adding two independent normally-distributed random variates: one with variance 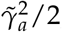, where 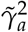 is the phenotypic variance in the population after selection, representing segregation variance (Lynch & Walsh (1998)), the other with variance 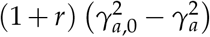, representing mutation. We chose this value for mutational variance for phenomenological rather than mechanistic reasons, as it maintains constant phenotypic and genotypic variances at equilibrium (see Appendix S4). Then, each individual’s phenotype is determined by adding a normally-distributed random variate with mean zero and variance 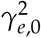 (the environmental component of the phenotype) to its breeding value. The phenotype, breeding value, and environmental components are fixed throughout an individual’s lifespan. When breeding values within the population are normally distributed (as we assume), breeding values within newborn cohorts are normally distributed with mean 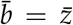 and variance 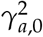 and the environmental components are normally distributed with mean zero and variance 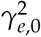. In the newborn age class, breeding values and non-genetic components are uncorrelated such that 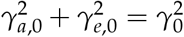. We define the heritability of the trait as 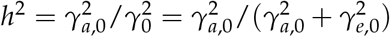.

Individuals in age class *k*, having experienced *k* rounds of selection, will have phenotypic variance 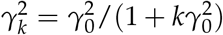 as noted above. However, each round of selection produces negative correlations between breeding values and environmental components within the cohort. For example, the joint distribution of *b* and *e* among survivors of one round of selection is

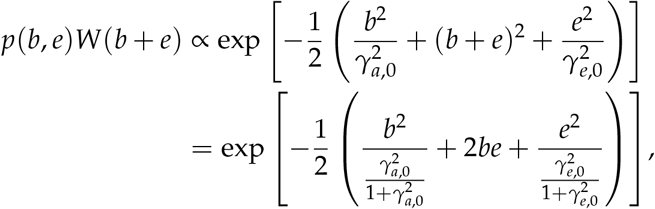

which specifies a bivariate normal distribution with variances 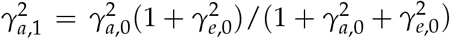 and 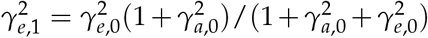 and correlation 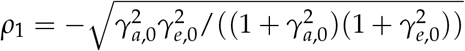. The correlation, first noted by Arnold & Wade (1984), is induced by the nonlinearity in the Gaussian selection function with respect to the phenotype, *z*, which is the sum of components *b* and *e*. Correlation will be induced for any selection surface that is a nonlinear function of the phenotype. The correlation is negative because the optimal phenotype can be reached by multiple combinations of *b* and *e*, with an increase in one necessitating a decrease in the other to maintain the phenotype. In Appendix S5, we show that for age class *k*,

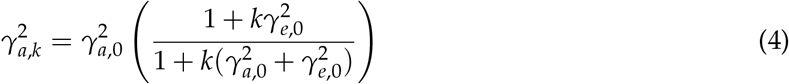

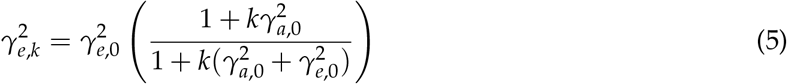

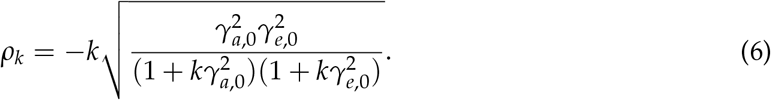

Although the phenotypic variance approaches zero as cohorts age, variances in breeding values and environmental components asymptotically approach non-zero quantities. Surprisingly, this suggests that older age classes may contain considerable genetic variation even as they might contain very little phenotypic variation. As the phenotypic variance approaches zero, *b* and *e* approach perfect negative correlation (*ρ*_*k*_ approaches −1 as *k* grows large). Increasing longevity will increase the proportion of the population in these older age classes, increasing the degree of correlation among phenotypic components in the population.

### Response to environmental change

We now consider an environmental change such that the mean population phenotype 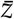 is distance *θ >* 0 from the novel environmental optimum. Say that the environmental shift occurs in time step 0 such that *t* ≥ 0 is the number of time steps since the environmental change. As in other classic models of rescue (Gomulkiewicz & Holt (1995), Kopp & Matuszewski (2014)), we model a constant environment after the shift.

### Demographic effects of pace-of-life trade-off

Fig. 1a plots *λ* as a function of mean population phenotype, 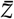, estimated numerically for three different life-history treatments with the same phenotypic variance (i.e., the same *γ*^2^ across life-history groups). See Appendix S6 for an expression for *λ* and information about numerical estimation and approximation. Higher longevity (higher 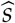) and the associated lower 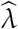 imply that *λ >* 1 for a narrower range of population phenotypes. This suggests that high-longevity populations will require rescue (adaptation to recover *λ >* 1) for a wider range of phenotypic shifts than low-longevity populations. Furthermore, for a given environmental shift that pushes *λ* below one, higher longevity will require more adaptation (greater change in 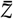 over time) to recover positive growth rates. For illustration purposes, we demonstrate potential demographic consequences of this trade-off in Fig. 1b. Here, we use values of *λ* for the three different life-history groups from Fig. 1a in the analytical expression for population size over time of Gomulkiewicz & Holt (1995) (see Appendix S7 for more information). Estimates in this figure assume that rates of phenotypic change are identical among treatments; we demonstrate in the next section that this assumption is unlikely to be met. However, this figure demonstrates the consequences of constrained population growth rates associated with longevity as well as the result evident in Fig. 1a that higher longevity can mean a longer time to recover positive growth rates. Thus, in the absence of variation in speed of evolution among life-history groups, the trade-off between survival and intrinsic growth rates implies longer declines following an environmental shift.

**Figure 1:**
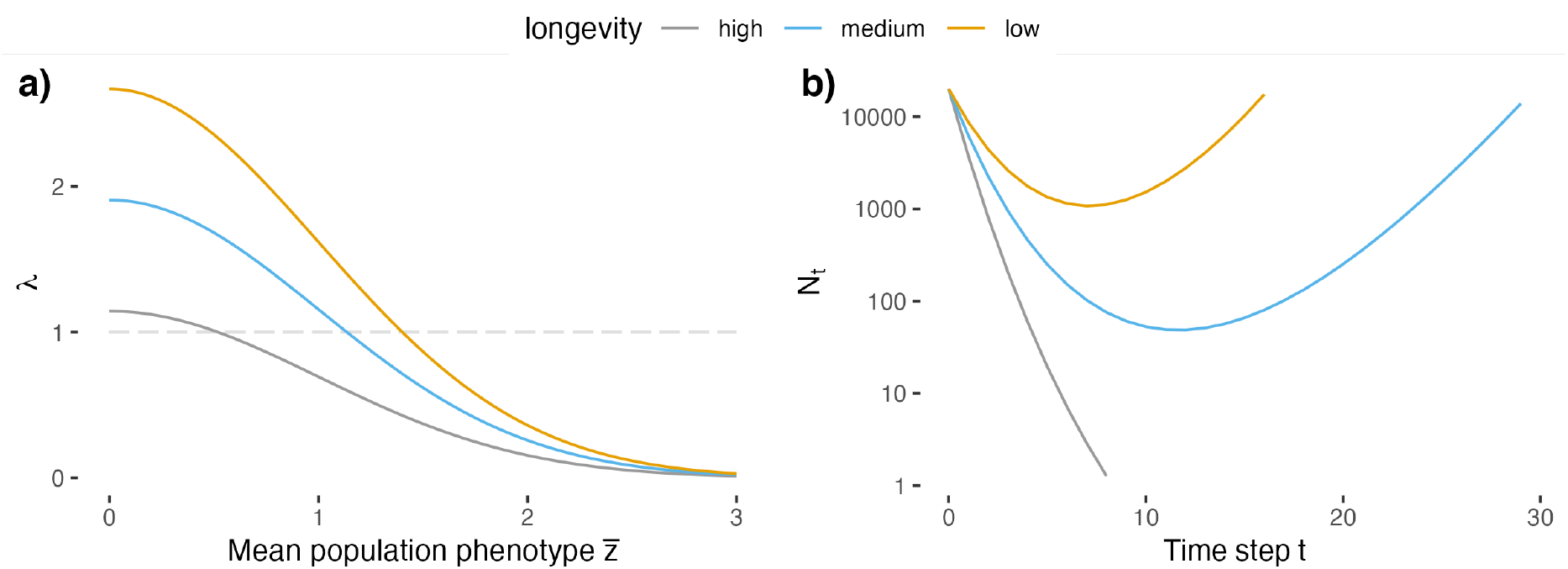
(a) *λ* as a function of mean phenotypic distance from the environmental optimum, 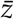, for different life-history strategies. High, medium, and low longevity curves correspond to 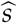 values of 0.9, 0.5, and 0.1, respectively. Horizontal gray dashed line is *λ* = 1. (b) Analytical solutions for population size over time for the same three life-history strategies according to the model of Gomulkiewicz & Holt (1995), with *θ* = 2, *N*_0_ = 20000, *γ*^2^ = 0.1, and *h*^2^ = 0.5 for each life history (see Appendix S7 for more detail).

### Adaptation within cohorts

Consider the cohort of individuals of age *k* at the time of the environmental shift. Denote the mean phenotype, breeding value, and environmental phenotypic components in age class *k* at time step *t* as 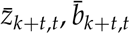 and 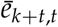, respectively. For all *k* at the time of the environmental shift, 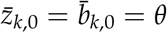 and 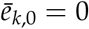 in expectation.

Lande (1976) found expressions for rates of phenotypic change under directional Gaussian selection; it can be shown using these results that the mean phenotype of the cohort age *k* at the time of environmental change after one round of selection is 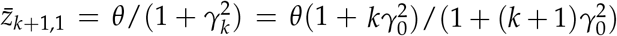. Applying this recurrence relationship iteratively, after *t* rounds of selection the mean cohort phenotype will be

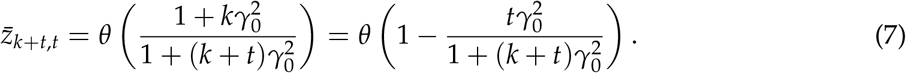

In expectation the mean phenotype for each cohort approaches zero (the optimum), but this rate is slower for older cohorts due to their reduced phenotypic variance at the time of the environmental change. This result also shows that cohort phenotypic change occurs independently of heritability, i.e., cohort trait evolution does not proceed more slowly for traits with lower heritabilities as is true in other models of rescue that lack age or stage structure (e.g., Gomulkiewicz & Holt (1995)). This is because of the distinct contributions of fertility and viability selection (Coulson & Tuljapurkar (2008)) to phenotypic change. Under fertility selection, an individual’s contribution to the population’s phenotypic distribution in the next time step is through offspring phenotypes; in this case, selection is only capable of acting on the heritable portion of the phenotype (i.e., the breeding value, Falconer & Mackay (1996)). Under viability selection, an individual contributes its own phenotype to the population’s phenotypic distribution in the next time step; in this case, the entire phenotype is exposed to selection.

In Appendix S5, we show that the cohort of age *k* at the time of the environmental shift will have the following mean breeding value and environmental components after *t* rounds of selection:

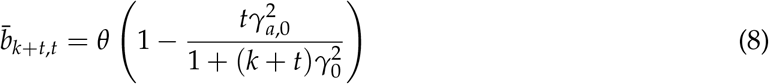

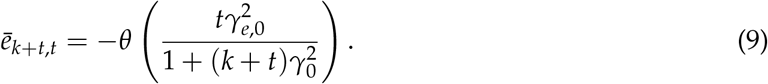

Variances and correlation between the components are the same as defined in Equations 4-6. Unlike Equation 7, Equation 8 demonstrates that the rate of genotypic change depends on the heritability of the trait (i.e., on the magnitude of 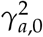 relative to 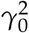.) Mean cohort breeding values will never be closer in expectation to the environmental optimum than the mean cohort phenotype, i.e., 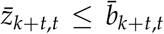. Equality is achieved for *t > k* only when 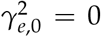, i.e., the trait is perfectly heritable. For 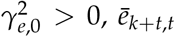 falls below 0, becoming more negative as *t* increases. For fixed 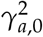, increasing 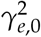 will increase the rate of change in the mean cohort phenotype (by increasing 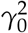; see Equation 7) but Equation 8 demonstrates that it will also slow down the rate of genotypic change (producing a more positive 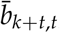, i.e., a mean breeding value further from the novel optimum). This is the decoupling of phenotypes and genotypes noted in other models of adaptive dynamics (e.g., Arnold & Wade (1984), Orive et al. (2017), Cotto & Chevin (2020)).

We can conclude that longevity can have the following effects on cohort evolutionary dynamics in our model. Longevity increases the proportion of the population in older cohorts. Older cohorts, with their reduced phenotypic variance, will on average have slower rates of phenotypic change following an environmental shift compared to younger cohorts. Another consequence of this reduced phenotypic variance in older cohorts is that following a severe environmental change, fewer individuals in older cohorts are likely to have phenotypes close to the novel optimum; this suggests mortality may be particularly high in these older cohorts. Thus, life histories with higher longevity can suffer reduced rates of adaptation compared to life histories with lower longevity. However, longevity increasing the mean population age will also increase the number of rounds of selection individuals in the population face. Under imperfect heritability, this will increase the degree of phenotypic-genotypic decoupling within the population upon environmental change. The decoupling results from viability selection, such that phenotypic change can proceed more quickly than under fertility selection because under viability selection an individual’s entire phenotype can respond to selection, rather than only the heritable portion. Thus, in our model, longevity may be associated with increased phentoypic change (and decreased genotypic change) due to an increased degree of phenotypic-genotypic decoupling in the population. This apparent tension can be further examined using simulations.

The analysis thus far has focused on the dynamics of cohorts alive at the time of the environmental shift (for whom the expected phenotype at the time of the environmental shift is *θ*), but these results also hold for cohorts born after the environmental shift if *θ* is replaced by 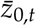 for *t >* 0. However, because we model random mating that does not depend on parental age, mean phenotypes of each new-born cohort depend on the distribution of breeding values within the population as a whole. We are unable to make analytical progress in deriving tractable expressions for the mean breeding value in the population at large following the environmental change, even if we make the assumption that the population remains at stable age distribution following the environmental change. In any case, this assumption is very unlikely to be met, as the reduced phenotypic variance in older cohorts means that fewer individuals in older cohorts will be close to the novel environmental optimum, in which case mortality will be higher in older cohorts. Deviation from the stable age distribution influences not only mean breeding values in successive cohorts, but also variance in breeding values. This means that the predictions made in this section will hold at the level of individual cohorts but might not hold at the population level. We thus conclude analytical discussion of the model and turn to simulation methods to assess effects of longevity on growth and adaptation at the population level.

### Simulation experiment

We analyzed the effects of longevity on evolutionary rescue in our model using a simulated experiment as follows. Our simulation experiment was a three-way factorial design with variables longevity 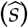, population-wide phenotypic diversity (*γ*^2^), and trait heritability (*h*^*2*^). We set 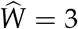 for all life histories. For a fixed 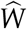, longevity was determined by the balance of the variables *r* and 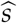. We had three life-history treatments: high longevity 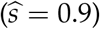, medium longevity 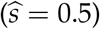, and low longevity 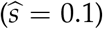, with *r* set appropriately for each. Because the phenotypic variance in the population is important for determining rates of phenotypic change (Lande (1976), Houle (1992)), we modeled treatments for high (*γ*^2^ = 0.4), medium (*γ*^2^ = 0.25), and low (*γ*^2^ = 0.1) phenotypic diversity at the population level. Finally, we included heritability of the trait (i.e., 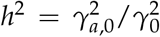), as the analysis above demonstrates that non-heritable phenotypic variance 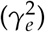 decouples mean genotypes and phenotypes within a cohort as individuals go through multiple rounds of selection. We used heritability values of *h*^2^ = 0.25, *h*^2^ = 0.5, and *h*^2^ = 1. The first two values are realistic for life-history traits (Geber & Griffen (2003), Wood et al. (2016)) and the final value, for which 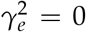, serves as a “control” for examining the decoupling of genotypes and phenotypes. Table 1 gives 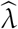 and *λ*^*^ for each life-history strategy. See Appendix S8 and Figs. S2-16 for simulation validation of analytical expressions at these parameter values. We set variance components within a life-history strategy by using the *γ*^2^, *r*, and 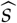 values to solve numerically for 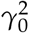 and 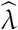, then set 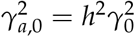 and 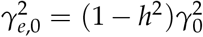; see Appendix S9 for detail on numerical estimation and the relationship between *γ* and 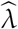.

**Table 1:** Maximum survival 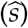 and phenotypic variance (*γ*^2^) treatments in simulation experiments, and their corresponding fecundity per mating bout (*r*), maximum population growth rate 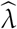, and equilibrium population growth rate (*λ*^*^) for each treatment. Heritability treatments are not included in this table as they do not directly influence equilibrium population growth rates.

| $\hat{s}$ | (longevity) | $\gamma^2$ | $r$ | $\hat{\lambda}$ | $\lambda^*$ |
| --- | --- | --- | --- | --- | --- |
| 0.1 | (low) | 0.10 | 27 | 2.8 | 2.67 |
| 0.1 | (low) | 0.25 | 27 | 2.8 | 2.50 |
| 0.1 | (low) | 0.40 | 27 | 2.8 | 2.37 |
| 0.5 | (medium) | 0.10 | 3 | 2.0 | 1.91 |
| 0.5 | (medium) | 0.25 | 3 | 2.0 | 1.79 |
| 0.5 | (medium) | 0.40 | 3 | 2.0 | 1.69 |
| 0.9 | (high) | 0.10 | 0.33 | 1.2 | 1.14 |
| 0.9 | (high) | 0.25 | 0.33 | 1.2 | 1.08 |
| 0.9 | (high) | 0.40 | 0.33 | 1.2 | 1.03 |

All populations were initialized at size *N*_0_ = 20000 by first drawing *N*_0_ random variates from the stable age distribution (see equation 2) then drawing each individual’s phenotypic components from a multivariate normal distribution according to the individual’s age. All populations were initialized with initial phenotypic distance from the novel environmental optimum *θ* = 2. To reduce unnecessary computational load, we set a carrying capacity of *K* = 20000; whenever a time step ended with a population size greater than *K*, individuals were removed at random until the population was at size *K*. Because individuals were removed at random, the carrying capacity did not affect the age structure or the phenotypic component variances and thus did not bias any mean demographic or trait characteristics in the population.

We ran two distinct simulation experiments. First, we ran a main batch of 1000 simulations per parameter combination (see Table 1) to determine mean population size and extinction rates over time. These simulations lasted 100 time steps or until the population went extinct. For each time step we estimated mean population size, including extinct populations as size zero, and the proportion of populations that had gone extinct in or prior to the time step. Second, to assess evolutionary dynamics and changes in age structure over time, we ran a separate batch of simulations. Unlike in the case of estimating mean population size, where extinct populations can be included in a mean as zeros, extinct populations introduce bias into unconditional estimates of age structure and phenotypic components. As such, this second batch of simulations was designed to reduce extinctions while still demonstrating effects of longevity and heritability on evolutionary dynamics. Thus, our second batch was run only with *γ*^2^ = 0.4, the highest level of population-wide phenotypic variance (shown in preliminary model testing to have the fewest extinctions) and with only 200 replicates per parameter combination (to limit the occurrence of trials from the early tail of the distribution of extinction times) with each trial lasting 50 time steps. For these simulations, we recorded the mean and variance of each phenotypic component (*z, b*, and *e*) and the number of individuals in each age class. We estimated expected size of each age class and expected mean and variance of each component for each time step within a parameter combination up until the first time step where a population went extinct.

### Simulations without life-history trade-off

To examine the effects of the life-history trade-off between longevity and intrinsic growth rate, we ran a third batch of simulations where life histories were set to have equal 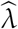 rather than equal 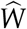. Under this experimental design, there was no trade-off between survival and reproduction, and longer-lived life-history strategies had higher maximum intrinsic fitness than their shorter-lived counterparts. All other aspects of the simulated experimental design were identical to our main simulation experiment. See Appendix S10 for details.

## Results

### Population size, growth, and extinction

In nearly all time steps mean population size was largest for low-longevity strategies and smallest for high-longevity strategies (Fig. 2). Low-longevity life histories were the fastest to return to their original size; low equilibrium growth rates (*λ*^*^) for the high-longevity treatments slowed the rate of population growth as populations of longer-lived organisms took considerably more time to return to their original size. Population sizes and growth rates increased with heritability and typically increased with increasing phenotypic variance; the one exception was high-longevity strategies, for which mean population size was highest at intermediate phenotypic variance rather than high phenotypic variance. This is explained by low phenotypic variance increasing extinction rates for all treatments (as demonstrated below), while high phenotypic variance decreased *λ*^*^ through increased variance load (Lande & Shannon (1996), Ashander et al. (2016)). For the high-longevity strategies, low 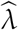 constrained *λ*^*^ to be very close to 1 such that population growth was constrained even though high phenotypic variance allowed for more rapid phenotypic change.

**Figure 2:**
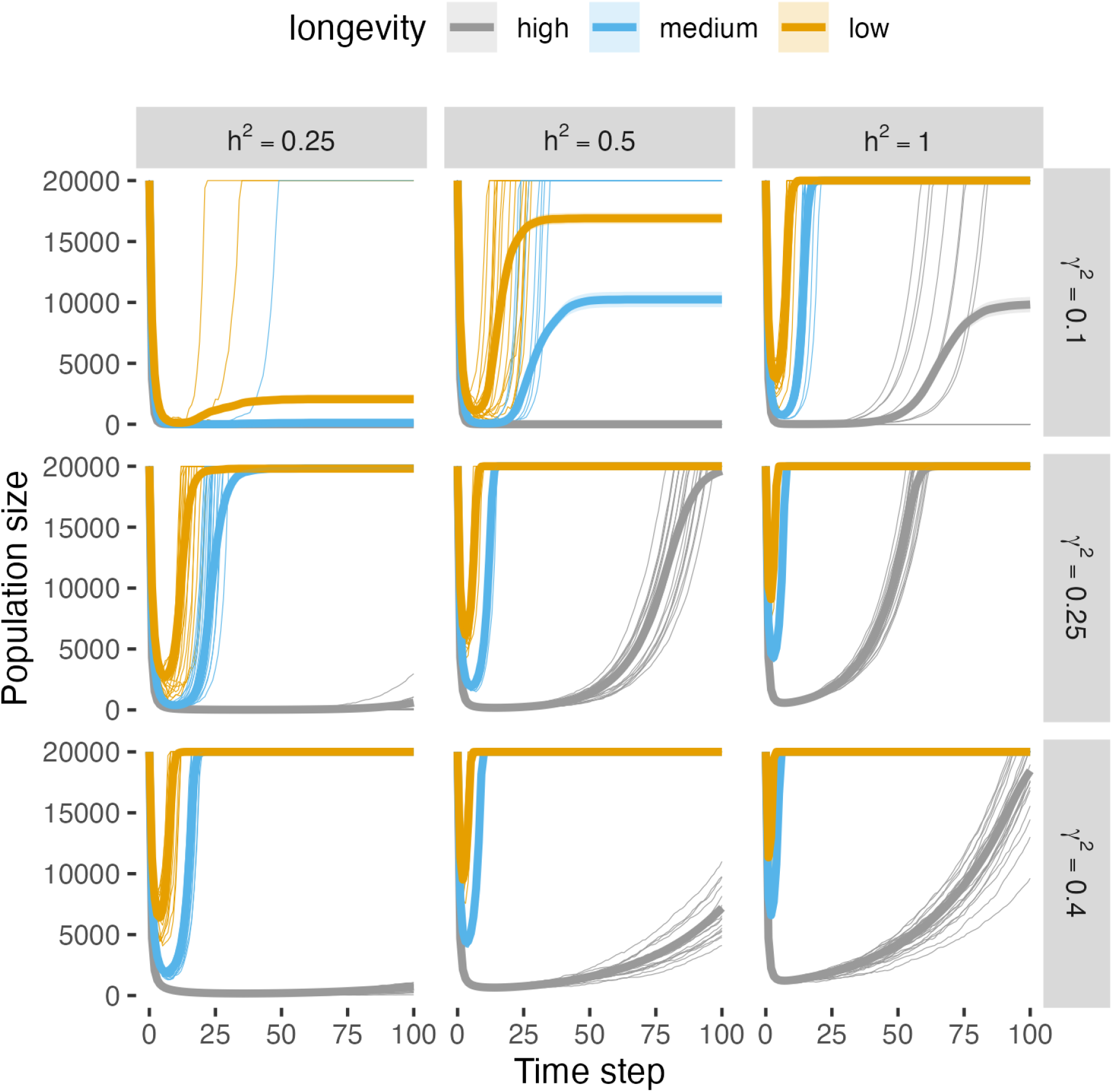
Mean population size over time. Thick lines are the mean population size of 1000 trials per treatment, including extinct populations as size zero. Shaded regions are twice the standard error on each side of the mean (errors are mostly too small to be visible on this plot). Twenty randomly-selected realizations of population size over time per treatment are plotted with thin lines.

We observed extinctions in five of nine combinations of *γ*^2^ and *h*^2^ (Fig. 3, Table S2). As in classic models of rescue (e.g., Gomulkiewicz & Holt (1995)), extinction was more likely at low levels of phenotypic variance or heritability. At all levels of *γ*^2^ or *h*^2^ where extinctions occurred, lower longevity was associated with more extinction; these extinctions mostly occurred well before the end of the simulations. For combinations of *h*^2^ and *γ*^2^ where all longevities experienced extinctions, the earliest extinctions occurred in low-longevity life histories (Table S2); this is likely because the higher variance in growth rates associated with the faster pace of life produced a larger tail of particularly “unlucky” populations with especially poor growth. In all cases, the periods during which low- and medium-longevity strategies experienced extinctions were brief, while high-longevity strategies were at risk of extinction for much longer durations (Fig. 3). This comports with the result evident in Figure 2 where low- and medium-longevity populations were able to rapidly return to original size while high-longevity populations remained at low densities due to constrained growth rates.

**Figure 3:**
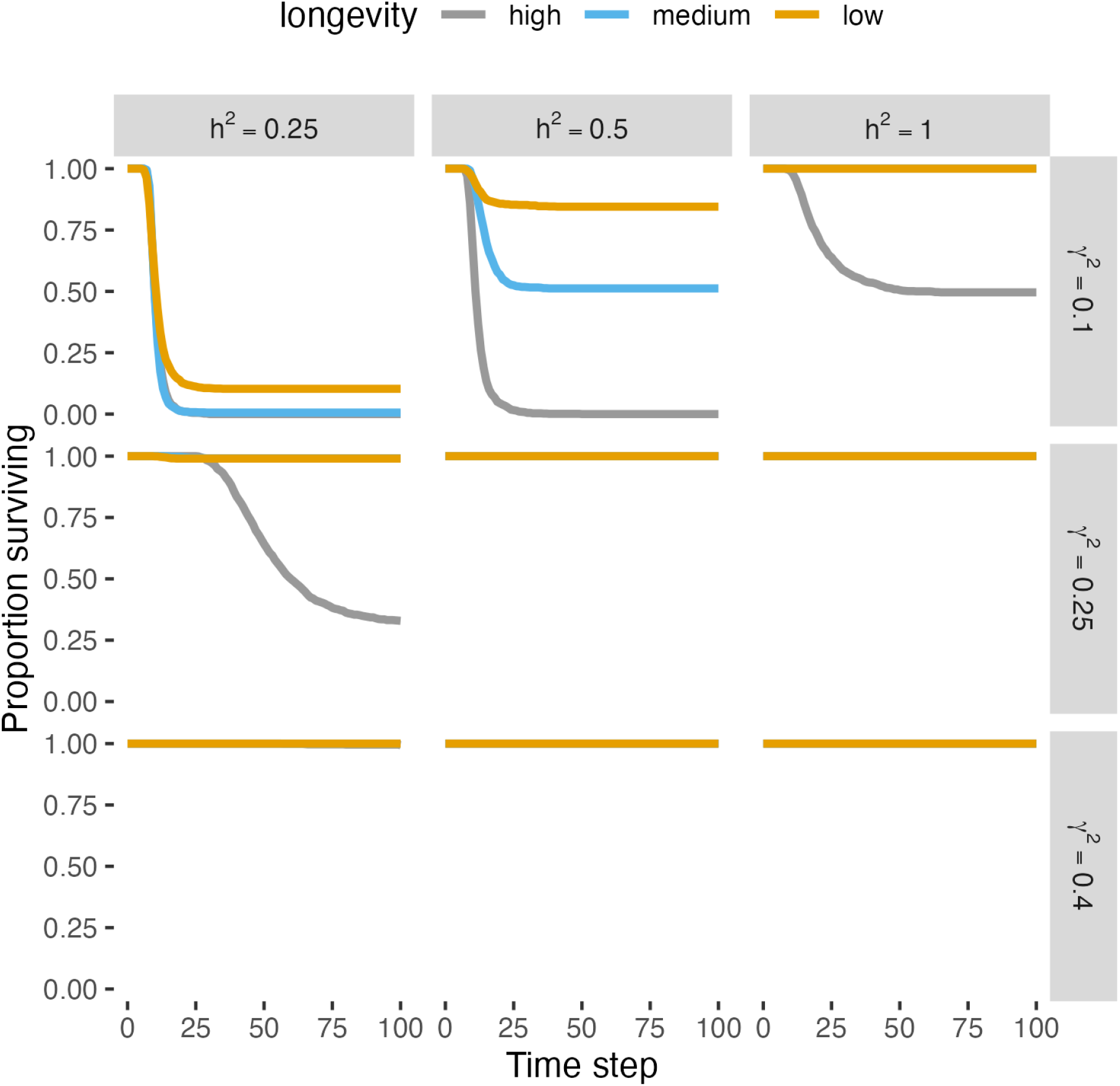
Proportion of surviving populations (out of 1000 per treatment) over 100 time steps.

Our auxiliary batch of simulations, where the life-history trade-off between survival and maximum growth rate was removed and instead all life histories had equivalent 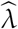, showed the opposite results to our main simulations. In particular, extinctions were more frequent and population sizes were smaller for the low-survival, low-longevity life histories (Figs. S17-18). Conversely, extinctions were rarer for longer-lived life histories.

### Rates of phenotypic and genotypic change

For imperfect heritabilities (i.e., *h*^2^ *<* 1) and high and medium longevity, we observed the predicted decoupling of mean genotype and mean phenotype at the population-level (Fig. 4), although it only occurred transiently. During this transient period, mean phenotypes rapidly approached the environmental optimum. This period can best be marked by two distinct phases in the dynamics of *ē*, the environmental component of the phenotype: a rapid (5-10 time step) period of increase in magnitude of *ē* (negative vertical direction in Fig. 4), followed by a longer period of relaxation as *ē* returned to zero. The durations of both phases increased with increasing longevity and decreasing heritability. During the first phase, there was a rapid rate of phenotypic change as populations approached, but did not necessarily reach, the environmental optimum. In high-longevity populations with larger changes in *ē*, the approach of *ē* to zero was associated with a slowing rate of phenotypic change, eventually falling behind rates of phenotypic change in lower-longevity populations. Thus, although there was an initial short phase of rapid phenotypic change, on longer timescales populations with higher longevity had a slower approach to the novel phenotypic optimum. During both of these phases, rates of genotypic change lagged behind rates of phenotypic change. This decoupling of phenotype and genotype disappeared under perfect heritability, under which there was minimal difference in rates of phenotypic change among longevity treatments. No extinctions were observed in any treatments in the smaller batch of simulations where adaptation was measured, precluding any survivorship bias. Differences in rates of phenotypic and genotypic change between medium- and low-longevity treatments were slight.

**Figure 4:**
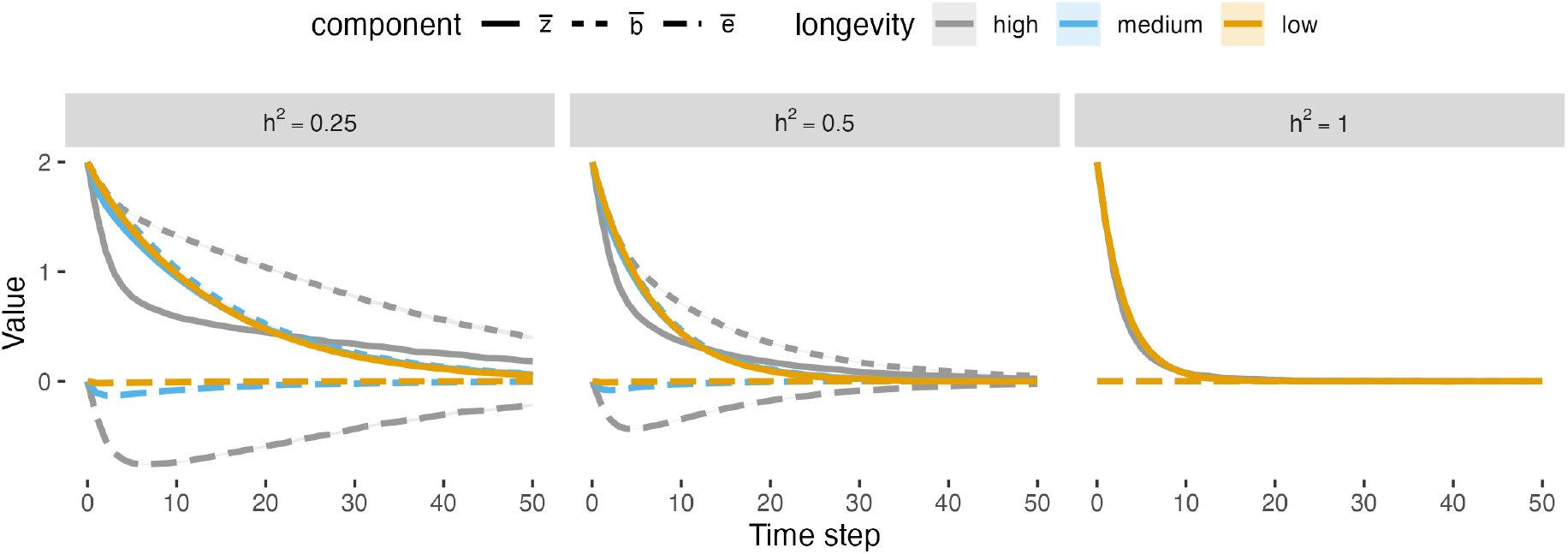
Mean phenotypic components (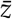, 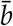, *ē*) for each longevity treatment over time for each of three levels of heritability. Initial (steady-state) population variance is *γ*^2^ = 0.4 in all panels. For all time steps, the phenotypic optimum is zero.

### Age- and age-variance structure

Environmental change perturbed high-longevity populations from the stable age distribution by disproportionately removing older individuals from the population, reducing the mean population age (Fig. 5). These populations returned to their pre-change steady state within 50 time steps; return to this steady state was faster with higher heritability. At low heritability, after the initial reduction in size for older age classes, the size of the older age classes “overshot” the stable age distribution such that they temporarily comprised a larger proportion of the age distribution compared to the steady state distribution. Age structure in medium- and low-longevity populations was minimally changed following the environmental change (Fig. S19).

**Figure 5:**
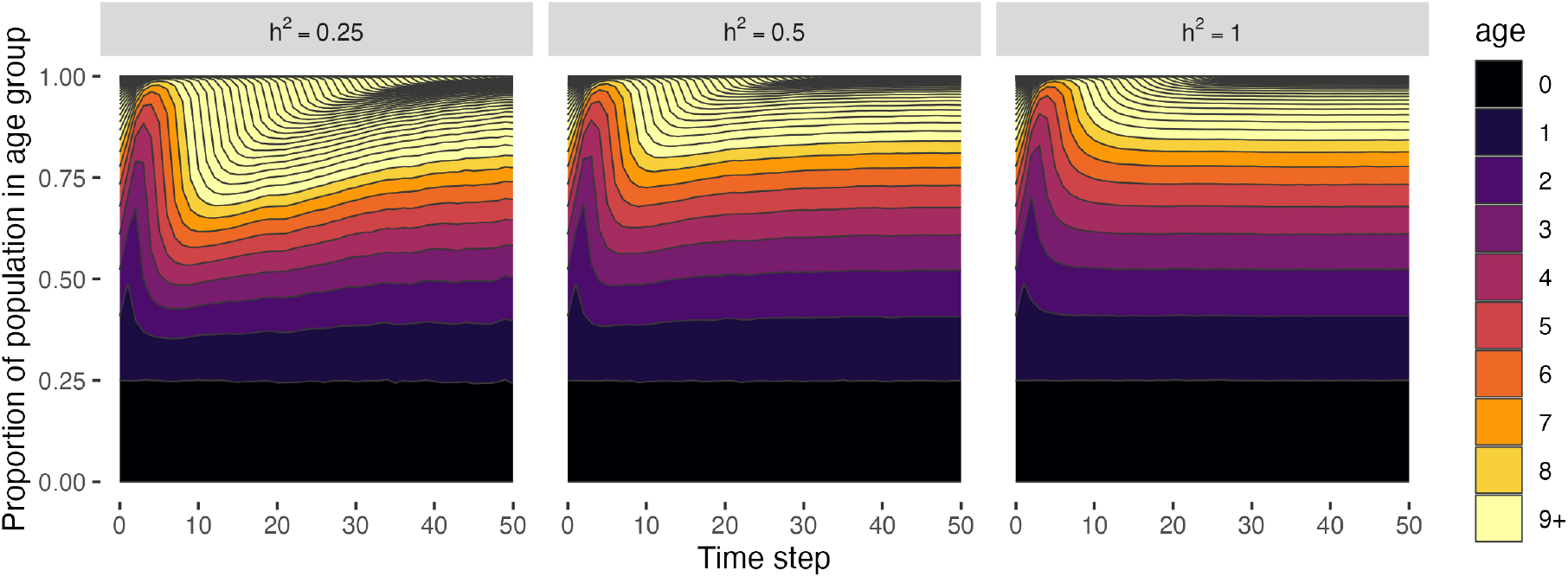
Mean age distribution over time for high-longevity populations at different heritability levels. Classes for ages nine and older are combined into one color code for visual clarity. Initial (steady-state) population variance is *γ*^2^ = 0.4 in all panels.

This disruption to the age structure in high-longevity populations coincided with an increase in phenotypic variance in the population at large (*γ*^2^), followed by a gradual return to equilibrium levels (Fig. 6). Specifically, phenotypic variance increased because the environmental change shifted the population’s age structure towards younger age classes (Fig. 5) with higher phenotypic variance. Medium-longevity strategies also experienced an increase in phenotypic variance, but the increase was much smaller in magnitude and shorter in duration. At perfect heritability, phenotypic variance (and therefore additive genetic variance, 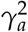) increased but then very quickly returned to equilibrium values, demonstrating that environmental change and disrupted age structure affects additive genetic variance in the population at large regardless of heritability. With lower heritability, the increase in phenotypic variance became slightly larger in magnitude and took longer to subside to equilibrium values; this coincided with a large increase in environmental phenotypic variance 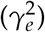, and the increase in additive genetic variance became considerably smaller. These results suggest that there was a transient disruption to population age structure, producing a transient increase in phenotypic variance available for selection to act on, but that with lower heritability most of this available phenotypic variance was in environmental components of the phenotype and thus response to selection was lost when producing new offspring.

**Figure 6:**
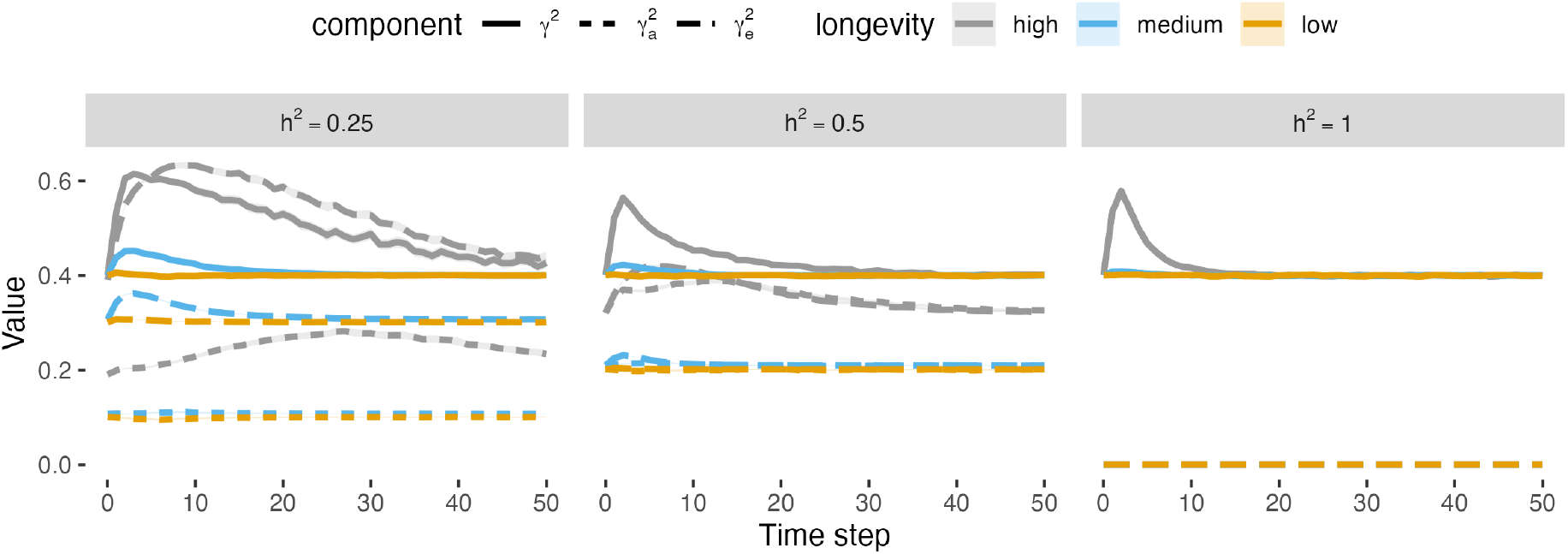
Phenotypic component variances over time.

## Discussion

We demonstrate that longevity, defined here as increased survival that is paired with reduced fecundity per reproductive bout, does impede evolutionary rescue following a single environmental shift as is often posited. However, in our model and simulation results, it was not due to the hypothesized mechanism of slower rates of phenotypic change. The primary difference of our model from others (highlighted below) is the presence of multiple selection episodes over the course of the lifetime, and we analyze our model incorporating the well-documented trade-off between longevity and annual or per-time step population growth (Oli (2004), Gaillard et al. (2005), Salguero-Gómez et al. (2016)). Our model found phenotypic change occurred at near-identical paces for long-and short-lived life histories with perfect trait heritability, and we observed a transient period of rapid phenotypic change for long-lived populations with imperfectly heritable traits. We also saw longer-lived populations took longer to recover positive growth rates and spent more time at lower density, where they were susceptible to extinction, whereas their shorter-lived counterparts were able to quickly return to their original size. Thus, we found conditions under which longer-lived populations were capable of rapid phenotypic change, but were ultimately at higher extinction risk due to slower growth rates. That is, longevity is likely to hinder evolutionary rescue by constraining maximum rates of increase, and while longevity can slow phenotypic change in some long-lived life-history strategies (Kuparinen et al. (2010), Cotto et al. (2017), Schmid et al. (2022)) it does not slow it in all cases.

We find that longevity does not necessarily slow phenotypic rates of change as is often predicted (Chapin et al. (1993), Vander Wal et al. (2013), Shoemaker & Lennon (2018)). Vander Wal et al. (2013) hypothesized that rescue occurs slowly in mammals, reasoning that the generations until rescue found in studies with model organisms in laboratory settings (e.g., Bell & Gonzalez (2009)), multiplied by longer generation times of mammals produced longer times until rescue. Our analysis highlights that this reasoning ignores potential change that can happen within generations. In fact, in our simulations, phenotypic change occurred more rapidly in longer-lived populations when estimating on a per generation (rather than per time step) basis (Appendix S13, Fig. S20). Prior models found phenotypic change was slower for longer-lived life histories (Kuparinen et al. (2010), Cotto et al. (2017), Schmid et al. (2022)), but these models had selection acting only once in a lifespan impeding the selection’s ability to remove maladapted, high-survival individuals from the gene pool. Our model instead allowed for selection in every time step in the lifespan and produced different evolutionary dynamics (Fig. 4), discussed below. Selection occurring once in the lifetime as modeled elsewhere and occurring in each time step as in our model are two extremes on a spectrum; intermediate cases likely exist in nature, as do many other varied cases such as the strength of selection varying by selection episode (Cotto & Chevin (2020)). We do not argue that the evolutionary dynamics we find apply to all long-lived organisms, but instead note that prior results finding that longevity slows adaptation will not hold for all long-lived organisms, and highlight that integrated selection effects over the lifespan produce within-generation evolutionary dynamics previously undocumented in the rescue literature.

In our simulations we found that repeated selection allowed for mean population phenotypes and genotypes to decouple at imperfect heritabilities, allowing for a transient phase of rapid phenotypic change that was offset by slowed genotypic change. This has been observed elsewhere in evolutionary theory (Orive et al. (2017), Cotto & Chevin (2020)) although its interactions with longevity have not been considered. This decoupling is important to conservation biology, as many life-history and demographic traits have relatively low heritability (Geber & Griffen (2003), Wood et al. (2016)). In age-structured populations, an individual potentially provides multiple contributions to changes in the population’s trait distribution over time, including contributions through offspring (subject to “fertility selection”) and a contribution through its own continued survival (subject to “viability selection”, Coulson & Tuljapurkar (2008)). Under fertility selection, only the heritable portion of the phenotype (i.e., the breeding value, Falconer & Mackay (1996)) is propagated forward, while under viability selection the entire phenotype is propagated. Put another way, under directional selection for, say, a large *z* = *b* + *e*, fertility selection selects for individuals with large *b* while viability selection simultaneously selects for large *b* and large *e*. Our analytical work (Equations 7 and 8) and simulations (Fig. 4) demonstrate that higher longevity, lower reproduction, and an older population will experience more rounds of viability selection and thus a greater degree of decoupling, increasing the rate of phenotypic change relative to a population where only fertility selection is occurring. This comes at a cost of decreased rates of underlying genotypic change; directional selection for individuals with either high *b* or high *e* will select for some individuals with low *b* but high *e*, who will then propagate relatively unfavorable breeding values into the gene pool. Our results are strikingly similar to those of Orive et al. (2017), who also found phenotypic-genotypic decoupling occurring in a model with both clonal and sexual reproduction, with a greater degree of decoupling occurring with more clonal reproduction. This similarity can be explained by clonality and survival similarly propagating the whole of the phenotype in response to selection. Our results also are similar qualitatively and in effect to transient buffering due to phenotypic plasticity (Lande (2009), Chevin & Lande (2010)), wherein there is a rapid phase of accelerated phenotypic change due to plastic effects (providing key buffering during population decline, Chevin & Lande (2010)) followed by a longer period as the phenotypic change is “assimilated” into change in the underlying genes.

Longevity and associated demographic features and trade-offs are well studied in life-history theory. This includes a trade-off between lifespan (and, by proxy, survival) and intrinsic growth rate (Cole (1954), Pianka (1970)). Results of our simulation were attributable in part to our simulation design, where expected lifetime fitness was held constant and survival and reproduction varied accordingly, producing a trade-off where high longevity produced low growth rates. Indeed, a separate batch of simulations run with all treatments having equal per time step growth rates (and having varying lifetime fitness) found favorable results for longevity (Appendix S10, Figs. S17-S18). Trade-offs between intrinsic growth rate and longevity (and its correlates, such as body size (Purvis et al. (2000)), body growth rate (Denney et al. (2002)), or litter size (Cardillo (2003))) are widely observed within and across taxa (Oli (2004), Gaillard et al. (2005), Salguero-Gómez et al. (2016)). Thus, while a trade-off was an assumption in our analysis, it is an important assumption for understanding and predicting responses of longer-lived organisms to sudden environmental change. We found that, although longevity can transitorily increase rates of phenotypic adaptation, likely buffering populations by increasing fitness, this effect is outweighed by constrained population growth that increases the risk of extinctions as populations remain at low density for longer (Pimm et al. (1988)), where they are vulnerable to stochasticity inherent to small population densities. Although not explicitly incorporated into our models, increased time spent at low density passes populations through a genetic bottleneck (Whitlock (2000)), possibly slowing rates of phenotypic change (Frankham et al. (1999)) and making populations vulnerable to entering an extinction vortex (Gilpin & Soulé (1986), Nordstrom et al. (2023)).

One surprising result of our analysis is that longevity and associated reduced population growth rates implied a smaller range of environmental shifts or perturbations (values of *θ*) that a population can withstand without incurring decline. This is best illustrated by Fig. 1a: the higher-longevity population has a narrower range of mean phenotypes 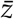 for which *λ* ≥ 1. The same result can be shown by considering the expected lifetime fitness; we show in Appendix S14 that increasing longevity steepens the lifetime selection pressure on *z*, likewise narrowing the phenotypic range in which individuals can replace themselves (Fig. S21). This is counter to observations that high survival provides buffering against increased environmental variability in natural populations (Morris et al. (2008), Dalgleish et al. (2010)). Our model applies the same per-episode degree of selection, but buffering may perhaps be obtained by scaling the degree of selection to produce similar lifetime selection pressures, or to produce a more gradual decline from 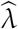 as 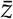 increases away from zero for the high-longevity curves in Fig. 1a. This would require at least one additional model parameter.

Our analysis also provided surprising connections to other results in demographic theory. Repeated selection on survival-related traits (as in our model) provides one evolutionary mechanism for the “frailty effect” of Vaupel et al. (1979), wherein mean survival or fitness within a cohort increases as it ages. Increasing survival with age is a common (although not universal, Jones et al. (2014)) life-history pattern; this mechanism through repeated selection can occur without plasticity or change in underlying traits. Frailty arising from phenotypic selection is particularly relevant to the demography of perennial organisms subject to selection on body size, birth weight, or proxies thereof (Milner et al. (1999), Coulson & Tuljapurkar (2008), Ozgul et al. (2010)). The frailty effect can produce “cohort selection” (Kendall et al. (2011)), which increases population growth rates relative to populations with homogeneity in survival. Our analysis suggests that with a sudden environmental shift, the frailty effect and its associated reduction in phenotypic variance among older cohorts can negatively affect population dynamics through disproportionately high mortality among these older, suddenly maladapted cohorts (Fig. 5). If the frailty effect and cohort selection did buffer long-lived populations in our simulations, it had small effects compared to other factors that increased extinction risk under high longevity.

From these results we can predict the following. For a single, drastic, and sustained change in the environment, longer-lived populations will be the most likely to go extinct, although these extinctions will occur on longer time horizons than extinctions in shorter-lived populations. The first species to go extinct may be short-lived; this is due to increased stochasticity in population growth rates for fast life histories. The higher variance in population growth trajectories could create a heavy “tail” of especially unlucky or poor populations that face extinction risk even if the “typical” population is unlikely to go extinct. This is consistent with recent work in plants (Compagnoni et al. (2021)) and mammals (Jackson et al. (2022)) showing stronger responses to climatic anomalies in shorter-lived species. However, as time since the environmental change increases, our model predicts that extinctions among surviving short-lived species should become rare as populations sufficiently adapt and return to a large size, while extinctions will continue for longer-lived species. These longer-lived species will adapt, but their constrained population growth rates will keep them at smaller densities for longer, leaving them vulnerable to extinction. These predictions are similar to those of Pimm et al. (1988), who suggested that large body size (which typically correlates with high survival and longevity) can buffer populations at small density and forestall extinction on short timescales, but will increase extinction risk on longer timescales by constraining growth and keeping populations at smaller densities where they are more susceptible to stochasticity or catastrophe. The net effect is that long-lived life histories are less likely to be successfully rescued than short-lived life histories. We can also form predictions about rates of adaptation: if the trait in question is susceptible to selection multiple times over the lifespan, on short timescales phenotypic change will occur more rapidly for longer-lived life histories than shorter-lived ones. The underlying decoupling of mean population phenotypes and genotypes can be uncovered by comparing the phenotypes of older parents with their offspring. For example, Coulson & Tuljapurkar (2008)’s study of red deer found the birth weights of offspring were smaller than those of their parents, which would be consistent with repeated selection on birth weight on parents resulting in parental cohorts with phenotypes that were larger than those associated with their breeding values.

Of course, fully integrating the theories of life history and evolutionary rescue will require a more complicated model and analysis than we present here. Our model assumes single, age-invariant parameters for survival and growth, but life histories often have varying investment in growth, survival, and reproduction over the course of the life cycle (Gadgil & Bossert (1970), Jones et al. (2014)). We intentionally modeled populations with simple demography, isolating effects of longevity while removing other life-history features (such as life stages buffered from selection) that are not necessary to produce longevity, even if they sometimes appear in the life histories of long-lived organisms. Our model and results can serve as a null model for comparison if comparing influences of other features that may coincide with longevity, such as senescence. We suspect it is also possible to extend the model to include varying reproduction and survival over the course of the lifetime without extreme complexity by, e.g., modeling survival using the discrete Weibull distribution, adding only one additional model parameter (Nakagawa & Osaki (1975)). We also assumed a simple form of density dependence, acting on survival independently of age or trait value. We chose this because selection on fecundity would have produced systematic deviations from the stable age distribution. However, rescue outcomes can vary depending on which vital rates or life stages are subject to selection, especially when considered in tandem with vital rates subject to density dependence (Klausmeier et al. (2020), Draghi et al. (2024)). Finally, future analysis must also generalize to other regimes of environmental change. As with other classic models of evolutionary rescue (Gomulkiewicz & Holt (1995), Chevin & Lande (2010), Kopp & Matuszewski (2014)), we modeled a single environmental shift but constant conditions thereafter. Further work should consider dynamics under gradually shifting and fluctuating environments (Bürger & Lynch (1995)). Outcomes again may depend on balances between demographic and evolutionary effects of longevity. Longevity and high survival are advantageous in variable environments (Orzack & Tuljapurkar (1989), Morris et al. (2008)), so increasing the magnitude of fluctuations relative to the magnitude of directional change might produce increasingly favorable results to longer-lived strategies. However, the effects of longevity on adaptation under gradual change or fluctuations might instead favor intermediate levels of generational overlap (Yamamichi et al. (2019)), or might depend critically on the scales of environmental variation or autocorrelation relative to the lifespan of the organism (Cotto & Chevin (2020), Peniston et al. (2021)) and will depend on the heritability of the trait (Lyberger et al. (2021)).

## Data archiving statement

We have included an anonymized version of code to reproduce our simulated experiment and analysis, as well as code to validate analytical expressions and raw data from simulation output, with our manuscript upload. A non-anonymized version of our repository was previously uploaded publicly to GitHub but is currently set to private. We pledge to upload all code and output data to Dryad upon acceptance of the manuscript.

## Acknowledgements

The authors would like to thank Dan Doak, Nancy Emery, and Aaron Westmoreland for many helpful discussions and suggestions that shaped the project and ensuing manuscript. The authors would also like to thank Nicolas Loeuille and two anonymous peer reviewers for suggestions that greatly improved the manuscript. SWN was supported by NSF grant award 1930222 to BAM and a dissertation completion grant from the University of Colorado Boulder’s Department of Ecology and Evolutionary Biology.

