## Supplementary material for "Longevity hinders evolutionary rescue through slower growth but not necessarily slower adaptation": R Code: AmNat_MS_template.pdf

### Template and Guidelines for Using LaTeX in *The American Naturalist*

Owen E. Cook[1,*]

Generic H. Collaborator[2,†]

Additional Q. Expert[3]

1. University of Chicago, Chicago, Illinois 60637;

2. University of Toronto, Toronto, Ontario M5S 1A5, Canada;

3. Middle Eastern Technical University, Çankaya, Ankara 06800, Turkey.

† Deceased.

*Manuscript elements*: Figure 1, figure 2, table 1, appendix A (for print; including figure A1, figure A2, and table A1), supplemental PDF. Figure 2 is to print in color.

*Keywords*: Examples, model, template, guidelines.

*Manuscript type*: Article.

Prepared using the suggested LaTeX template for *Am. Nat.*

#### Abstract

Lorem ipsum dolor sit amet, consectetur adipiscing elit. Sed non risus. Suspendisse lectus tortor, dignissim sit amet, adipiscing nec, ultricies sed, dolor. Cras elementum ultrices diam. Praesent quis dolor in dolor molestie cursus et ac nisi. Vestibulum ante purus, semper eget est vitae, vehicula ornare nisl. Morbi efficitur euismod enim, nec feugiat tellus cursus eget.

### Introduction

The quick red fox jumps over the lazy brown dog. Furthermore, the quick brown fox jumps over the lazy red dog. In addition, the quick Rüppell's fox (*Vulpes rueppellii*) jumps over the lazy golden retriever.

### Methods

Lorem ipsum ut velit mauris, egestas sed, gravida nec, ornare ut, mi. Aenean ut orci vel massa suscipit pulvinar. Nulla sollicitudin. Fusce varius, ligula non tempus aliquam, nunc turpis ullamcorper nibh, in tempus sapien eros vitae ligula (**?**).

#### *Second-order heading*

Lorem ipsum nulla facilisi, pace **?**. Etiam semper, orci sit amet facilisis interdum, tellus nunc consequat erat, quis viverra nisi diam ut metus. Pellentesque cursus, sapien malesuada euismod iaculis, mauris purus interdum diam, vel vestibulum justo enim vitae tellus. Nunc interdum lorem sit amet diam volutpat tristique. Quisque pulvinar ac metus commodo lacinia (**??**).

##### *Third-order heading*

Usually two or three levels of heading will be all you need. Journal style even permits a fourth level in case you need it.

*Fourth-order heading.* The quick red fox jumps over the lazy brown dog in this paragraph as well. Donec mauris nibh, volutpat vehicula viverra at, iaculis congue sem. Praesent eget erat rhoncus erat sollicitudin volutpat.

$$\frac{1}{N_k - 1} \sum_{t=1}^{N_k} (M_{tjk} - \bar{M}_{jk})^2 \qquad (1)$$

Praesent quis dolor in dolor molestie cursus et ac nisi. Vestibulum ante purus, semper eget est vitae, vehicula ornare nisl. Morbi efficitur euismod enim, nec feugiat tellus cursus eget. Donec mauris nibh, volutpat vehicula viverra at, iaculis congue sem.

##### *Another second-order heading*

As **?** argued, phasellus porttitor eros et ante condimentum, eget facilisis orci condimentum. Nulla facilisi. Proin placerat elit blandit, euismod dolor nec, dapibus diam. Mauris posuere malesuada lacus, at elementum lacus auctor eu (fig **??**A).

#### Results

Lorem ipsum dolor sit amet. Aenean pulvinar malesuada commodo (see **?**; table **??**). Sed aliquet mauris odio, in tristique dui egestas a. Etiam eu malesuada quam. Suspendisse tincidunt eu erat sit amet vulputate. Duis at arcu et nisl dictum mattis. Maecenas vel cursus ante. Cras eleifend elit nec velit sollicitudin fermentum in ac mauris. Pellentesque rutrum magna vel elit maximus hendrerit.

##### *The height of the jump*

Aenean eu pellentesque quam (fig. **??**). Nam pellentesque augue eu finibus lacinia. Nullam nec justo vitae odio imperdiet rhoncus vitae vitae quam. Pellentesque porttitor metus et lectus ornare, ac cursus urna efficitur (fig **??**B).

##### *The laziness of the dog*

Example paragraph with embedded references (video **??**, fig. **??**). If you have deposited data to Dryad, it is advisable to cite them somewhere in the main text (usually in the Methods or Results sections). A sentence like the following will do: All data are available in the Dryad Digital Repository (**?**).

### Discussion

Nam pulvinar lorem at lorem ultrices, vel accumsan massa feugiat (**?**). Proin tristique velit eget lacus iaculis, in pellentesque nulla varius. Phasellus sodales est odio, eu pulvinar magna pellentesque eu. Sed ut lobortis eros. Aliquam eget metus turpis. Sed et convallis lectus, id tincidunt enim. In porta nibh ut lacus feugiat, non consequat orci rhoncus. Morbi blandit at augue nec tempor. Sed fringilla ipsum ut justo viverra, ut euismod nisi gravida.

Curabitur non posuere augue, id suscipit orci. Nunc luctus accumsan aliquam. Cras egestas turpis vitae nisl vulputate interdum. Donec pellentesque libero egestas tortor pharetra laoreet. Phasellus facilisis auctor ligula, eu sollicitudin mi sagittis non.

### Conclusion

Duis pharetra enim at libero cursus, eu commodo mi vestibulum. Nullam eget velit nec lectus viverra sodales. Suspendisse egestas, eros at dictum tincidunt, mi orci laoreet libero, eget rutrum sapien arcu blandit odio.

### Acknowledgments

OEC would like to thank Madlen Wilmes, Gyuri Barabás, Flo Débarre, Vlastimil Křivan, and Greg Dwyer for their comments and suggestions on this template.

### Statement of Authorship

OEC conceived the experiments, collected the data, and wrote the original draft. GHC provided specimens and analyzed the model. AQE oversaw data analysis and developed the code. All authors reviewed and edited the writing at all stages of revision.

### Data and Code Availability

On initial submission, you may use this section to provide a URL for editors and reviewers that is 'private for peer review'. After acceptance, this section must be updated with correct, working DOIs for data deposits (such as on the Dryad Digital Repository, **?**) and code deposits (such as in Zenodo).

### Appendix A: Additional Methods and Parameters

#### *Fox–dog encounters through the ages*

The quick red fox jumps over the lazy brown dog. The quick red fox has always jumped over the lazy brown dog. The quick red fox began jumping over the lazy brown dog in the 19th century and has never ceased from so jumping, as we shall see in figure **??**. But there can be surprises (figure **??**).

If the order and location of figures is not otherwise clear, feel free to include explanatory dummy text like this:

[Figure A1 goes here.]

[Figure A2 goes here.]

#### *Further insights*

Tables in the appendices can appear in the appendix text (see table **??** for an example), unlike appendix figure legends which should be grouped at the end of the document together with the other figure legends.

Lorem ipsum dolor sit amet, as we have seen in figures **??** and **??**.

Table A1: Various rivers, cities, and animals

| River | City | Animal |
|---|---|---|
| Chicago | Chicago | Raccoon |
| Des Plaines | Joliet | Coyote |
| Illinois | Peoria | Cardinal |
| Kankakee | Bourbonnais | White-tailed deer |
| Mississippi | Galena | Bald eagle |

Note: See table **??** below for further table formatting hints.

#### References Cited Only in the Online Enhancements

Tytler, W. 1759. The Inquiry, Historical and Critical, into the Evidence against Mary Queen of Scots, and an Examination of the Histories of Dr. Robertson and David Hume with respect to that Evidence. W. Creech, Edinburgh.

### Tables

Table 1: Founders of *The American Naturalist*

| Early editor | Years with the journal |
|---|---|
| Alpheus S. Packard Jr. | 1867–1886 |
| Frederick W. Putnam | 1867–1874 |
| Edward S. Morse | 1867–1871 |
| Alpheus Hyatt | 1867–1871 |
| Edward Drinker Cope[a] | 1878–1897 |
| J. S. Kingsley | 1887–1896 |

Note: Table titles should be short. Further details should go in a ‘notes’ area after the tabular environment, like this.

[a] Published the first description of *Dimetrodon*.

### Figure legends

Figure 1: Figure legends can be longer than the titles of tables. However, they should not be excessively long—in most cases, they should be no more than 100 words each.

Figure 2: In this way, figure legends can be listed at the end of the document, with references that work, even though the graphic itself should be included for final files after acceptance. Instead, upload the relevant figure files separately to Editorial Manager; Editorial Manager should insert them at the end of the PDF automatically.

Figure A1: *A*, the quick red fox proceeding to jump 20 m straight into the air over not one, but several lazy dogs. *B*, the quick red fox landing gracefully despite the skepticism of naysayers.

Figure A2: The quicker the red fox jumps, the likelier it is to land near an okapi. For further details, see **?**.

Video S1: Video legends can follow the same principles as figure legends. Counters should be set and reset so that videos and figures are enumerated separately.
