## Supplementary material for "Longevity hinders evolutionary rescue through slower growth but not necessarily slower adaptation": R Code: AmNat_supp.tex

\documentclass[11pt]{article}
\usepackage[sc]{mathpazo} %Like Palatino with extensive math support
\usepackage{fullpage}
\usepackage[authoryear,sectionbib,sort]{natbib}
\linespread{1.7}
\usepackage[utf8]{inputenc}
\usepackage{lineno}
\usepackage{titlesec}
\usepackage{amsmath}
\usepackage{graphicx}
\titleformat{\section}[block]{\Large\bfseries\filcenter}{\thesection}{1em}{}
\titleformat{\subsection}[block]{\Large\itshape\filcenter}{\thesubsection}{1em}{}
\titleformat{\subsubsection}[block]{\large\itshape}{\thesubsubsection}{1em}{}
\titleformat{\paragraph}[runin]{\itshape}{\theparagraph}{1em}{}[. ]
\usepackage{fancyhdr}
\pagestyle{fancy}

%%%%%%%%%%%%%%%%%%%%%
% Header
%%%%%%%%%%%%%%%%%%%%%
%
% Customize the line below with the last name of your first author and
% the short title of your MS. You can comment authorship information out
% while your MS is undergoing double-blind review.
%
\rhead{Supplement to ``Longevity hinders evolutionary rescue \dots,'' \textit{Am.~Nat.}}
\setlength{\headsep}{0.3in}  
\lhead{} 

%%%%%%%%%%%%%%%%%%%%%
% Line numbering
%%%%%%%%%%%%%%%%%%%%%
%
% Please use line numbering with your initial submission and
% subsequent revisions. After acceptance, please turn line numbering
% off by adding percent signs to the lines %\usepackage{lineno} and
% to %\linenumbers{} and %\modulolinenumbers[3] below.
%
% To avoid line numbering being thrown off around math environments,
% the math environments have to be wrapped using
% \begin{linenomath*} and \end{linenomath*}

\title{Online Supplement: \\ 
Longevity hinders evolutionary rescue through slower growth but not necessarily slower adaptation}

%%%%%%%%%%%%%%%%%%%%%
% Authorship
%%%%%%%%%%%%%%%%%%%%%
% Please remove authorship information while your paper is under review,
% unless you wish to waive your anonymity under double-blind review. You
% will need to add this information back in to your final files after
% acceptance.
%
% Once accepted for publication, author-supplied PDFs should have a 
% title page that includes (at least) the authors' names, the title of 
% the MS, and the name of the journal. It should also have a header and
% page numbers.

% \author{Owen E. Cook$^{1,\ast}$ \\ 
% Generic H. Collaborator$^{2,\dag}$ \\ 
% Additional Q. Expert$^{3}$}

\author{Anonymous}

\date{}

\begin{document}

\maketitle

% \noindent{} 1. University of Chicago, Chicago, Illinois 60637;

% \noindent{} 2. University of Toronto, Toronto, Ontario M5S 1A5, Canada;

% \noindent{} 3. Middle Eastern Technical University, Çankaya, Ankara 06800, Turkey.


%\linenumbers{}
%\modulolinenumbers[3]


% In many cases, The American Naturalist allows authors to typeset their
% own supplementary material in an author-supplied PDF. This template
% applies to such cases. 
% 
% For appendices that will be typeset by the AmNat editorial staff, 
% please see the main LaTeX template, available from
% https://www.journals.uchicago.edu/journals/an/instruct
% Such appendices typically include descriptions of methods and tables
% defining parameters.
%
% In general, you have wide discretion for how you want to format an
% author-supplied PDF. They should in any case have a title page, 
% page numbers, and a header identifying the MS's (short) title.
%
% Counters for the online supplement should normally begin with an S
% (thus normally figure S1, figure S2, table S1, equation S1, etc.).

\renewcommand{\theequation}{S\arabic{equation}}
% redefine the command that creates the equation number.
\renewcommand{\thetable}{S\arabic{table}}
\renewcommand{\thesection}{S\arabic{section}}
\renewcommand{\thefigure}{S\arabic{figure}}
\setcounter{equation}{0}  % reset counter 
\setcounter{figure}{0}
\setcounter{table}{0}

% In online supplementary PDFs, sections can be numbered or not
% (at your discretion). If they are numbered, sections should usually
% begin with an S.

\newpage{}

\section{Phenotypic distribution after repeated selection}
\label{sec:02_pheno_normality}

Assume that the phenotypic distribution is centered at the environmental optimum (i.e., $\bar{z} = 0$) and that upon birth the variance of this distribution is $\gamma^2_0.$ Following selection, the distribution will be

\begin{align*}
    W(z) p(z) &  \propto \exp{\left( - \frac{z^2}{2} \right)} \exp{ \left( - \frac{z^2}{2 \gamma^2_0} \right)} \\
    & = \exp{ \left( - \frac{1}{2} \left( z^2 +  \frac{z^2}{\gamma^2_0} \right) \right)} \\
    & = \exp{ \left( - \frac{1}{2} \left( \frac{z^2_0}{\frac{\gamma^2_0}{1 + \gamma^2_0}}\right) \right)}.
\end{align*}

\noindent This demonstrates that the phenotypic distribution remains normal with mean $0$ and variance $\gamma^2_1 = \gamma^2_0 / (1 + \gamma^2_0).$ As expected, phenotypic variance shrinks with selection, with a greater proportional loss of variance as $\gamma^2_0$ increases, i.e., as selection strength increases relative to variance. 

It can similarly be shown that $\gamma^2_{k+1} = \gamma^2_k / (1 + \gamma^2_k).$ This recursive relationship is solved by $\gamma^2_k = \gamma^2_0 / (1 + k \gamma^2_0).$ The base case of $k = 0$ is demonstrated above. The inductive step to prove the relationship is as follows:

\begin{align*}
    \gamma^2_{k+1} & = \frac{\frac{\gamma^2_0}{1 + k \gamma^2_0}}{1 + \frac{\gamma^2_0}{1 + k \gamma^2_0}} \\ 
    & = \frac{\gamma^2_0}{1 + k \gamma^2_0 + \gamma^2_0} \\
    & = \frac{\gamma^2_0}{1 + (k+1) \gamma^2_0},
\end{align*}

\noindent proving the claim.

We make the simplifying assumption that the phenotypic distribution is centered on the environmental optimum. However, we demonstrate later in Appendix~\ref{sec:02_bivariate} that relaxing this assumption does not change the variance of the distribution, but instead only affects the mean.

\clearpage

%%%%%%%%%%%%%%%%%%%%%%%%%%%%%%%%%%%%%%%%%%%%%%%%%

\section{The frailty effect in adult survival}
\label{sec:02_frailty}

Consider that at birth, the phenotypic distribution ($z$) within a cohort is assumed to be normally distributed with variance $\gamma^2_0.$ For individual $i,$ $(z_i / \gamma_0)^2$ is $\chi^2_1$ distributed within the population. The $\chi^2_1$ distribution is equivalent to a gamma distribution with shape parameter $1/2$ and scale parameter $2$ (rate parameter $1/2.$) Letting $y_i = z_i^2 = \gamma^2_0 (z_i / \gamma_0)^2,$ $y_i$ will also be gamma distributed but with scale parameter $2 \gamma^2_0$ (rate $1/(2\gamma^2_0).)$ For a cohort born with a mean phenotype that is not matched to the environmental optimum, i.e.,  $\bar{z}_0 = \theta \neq 0,$ the frailties can be defined as $y_i = (z_i - \theta)^2$ to produce the same results so long as the environment remains constant over time.

While the analysis of Vaupel et al. (1979) allows for a ``force of mortality'' that can depend on time, age, and/or frailty, in our model individual mortality rates are independent of age by model construction and independent of time because we assume a constant environment. This latter point also allows satisfaction of another important characteristic of frailty: that an individual's frailty is constant over the course of its life, which occurs in this case because an individual's distance from the phenotypic optimum does not change with time if the environment is constant. Thus, we can let the force of mortality of individual $i$ be defined by its frailty, $y_i = z_i^2.$  As expected, frailty increases with increasing distance from the phenotypic optimum. Comparing two individuals, denoted individuals $i$ and $j,$ the relative odds of mortality of $i$ compared to $j$ is $y_i / y_j = (z_i / z_j)^2.$ Vaupel et al. (1979) demonstrated that a gamma-distributed population of frailties will remain gamma distributed as the cohort ages. Similarly, in our model, the frailties will remain gamma distributed, with similarly-shrinking mean and variance in frailties with time. This can also be seen by noting that successive rounds of Gaussian selection on a cohort will maintain a normally-distributed population of surviving phenotypes, with decreasing phenotypic variance as cohorts age, matching the result of Vaupel et al. (1979) that variance in frailties also decreases with cohort age.
% NOTE: references here are hard coded

\clearpage 

%%%%%%%%%%%%%%%%%%%%%%%%%%%%%%%%%%%%%%%%%%%%%%%%%

\section{Stable age distribution}
\label{sec:02_stable_age_dist}

Assume that the equilibrium population growth rate is $\lambda^*.$ Letting $l_k$ be the probability of survival to age $k$ and noting that the mean reproductive output for all age classes is $r,$ population dynamics can be described by the Euler-Lotka equation as

\begin{align*}
    \displaystyle\sum_{k=1}^\infty r l_k (\lambda^*)^{-k} = 1.
\end{align*}

\noindent For $k > 0$, cumulative survival until age $k$ is the product of survivals from ages $0$ through age $k-1.$

\begin{align*}
    l_k & = \displaystyle\prod_{j=0}^{k-1} \bar{s}_j \\
    & = \displaystyle\prod_{j=0}^{k-1} \widehat{s} \sqrt{\frac{1}{1+\gamma^2_j}} \\
    & = \widehat{s}^k \sqrt{\frac{1}{1+\gamma^2_0}} \sqrt{\frac{1}{1+\gamma^2_1}} \dots \sqrt{\frac{1}{1+\gamma^2_{k-1}}} \\
    & = \widehat{s}^k \sqrt{\frac{1}{1+\gamma^2_0}} \sqrt{\frac{1 + \gamma^2_0}{1+2\gamma^2_0}} \dots \sqrt{\frac{1+(k-1)\gamma^2_0}{1+k\gamma^2_0}} \\
    & = \widehat{s}^k \sqrt{\frac{1}{1+k\gamma^2_0}}.
\end{align*}

\noindent Substituting this in to the Euler-Lotka equation gives

\begin{align*}
    \displaystyle\sum_{k=1}^\infty r \widehat{s}^k \sqrt{\frac{1}{1+k\gamma^2_0}} (\lambda^*)^{-k} = 1.
\end{align*}

For all $k > 0,$ the $k$th term inside the above summation is proportional to the size of age class $k$ in the stable age distribution. Denote $p_k$ as the mass of age class $k$ in the stable age distribution. We require $p_0$ such that the sum of $p_k$ terms is $1.$ Note that for $k = 0,$ $l_k = 1$ and $(\lambda^*)^{-k} = 1.$ Thus the term $r l_0 (\lambda^*)^{-1} = r.$

\begin{align*}
    \displaystyle\sum_{k=0}^\infty r s^k \sqrt{\frac{1}{1+k\gamma^2_0}} (\lambda^*)^{-k} & = r + \displaystyle\sum_{k=1}^\infty r s^k \sqrt{\frac{1}{1+k\gamma^2_0}} (\lambda^*)^{-k} \\
    & = r + 1.
\end{align*}

\noindent Dividing both sides by $r+1$ and taking the $r$ out of the summation produces the following:

\begin{align*}
    \frac{r}{1+r} \displaystyle\sum_{k=0}^\infty \left( \frac{\widehat{s}}{\lambda^*} \right)^k \sqrt{\frac{1}{1+k\gamma^2_0}} = 1.
\end{align*}

\noindent The first term in the summation (i.e., $k = 0$) is equal to $1.$  We can thus conclude that $p_0 = r / (1+r),$ in which case, $p_k$ is equal to $p_0$ times the $k$th term in the summation, i.e., 

\begin{align*}
    p_k = \frac{r}{1+r} \left( \frac{\widehat{s}}{\lambda^*} \right)^k \sqrt{\frac{1}{1+k\gamma^2_0}}.
\end{align*}

\begin{figure}[!h]
    \centering
    \includegraphics[width=0.95\textwidth]{figs/figS1_age_distn.png}
    \caption{Stable age distribution as a function of phenotypic variance for three different $\widehat{s}$ values plotted on the natural scale (top row) and log scale (bottom row). For all panels, $\widehat{W} = 3.$}
    \label{fig:02_agedist}
\end{figure}

\clearpage


\section{Mutation rate}
\label{sec:02_mutation}

Using the convention in the main text, let $\tilde{\gamma}^2_a$ be the additive genetic variance in the population after selection; this is equivalent to the variance in adults (i.e., all individuals that are not in the new cohort) at mutation-selection balance. 

\begin{align*}
    \gamma^2_a = p_0 \gamma^2_{a,0} + (1 - p_0) \Tilde{\gamma}^2_a.
\end{align*}

\noindent Solving for $\Tilde{\gamma}^2_a$ gives the following:

\begin{align*}
    \Tilde{\gamma}^2_a = \frac{\gamma^2_a - p_0 \gamma^2_{a,0}}{1 - p_0}.
\end{align*}

\noindent We assume the additive genetic variance in offspring is the additive genetic variance in adults (i.e., parents) plus mutational variance, such that $\gamma^2_{a,0} = \Tilde{\gamma}^2_a + \gamma^2_m.$ Solving for $\gamma^2_m$ and substituting the expression for $\Tilde{\gamma}^2_a$ above gives

\begin{align*}
    \gamma^2_m & = \gamma^2_{a,0} - \frac{\gamma^2_a - p_0 \gamma^2_{a,0}}{1 - p_0} \\
    & = \frac{\gamma^2_{a,0} - \gamma^2_a}{1 - p_0}.
\end{align*}

\clearpage

%%%%%%%%%%%%%%%%%%%%%%%%%%%%%%%%%%%%%%%%%%%%%%%%%

\section{General form of bivariate distribution of $b_k$ and $e_k$}
\label{sec:02_bivariate}

Let $b_k$ and $e_k$ be, respectively, the distribution of breeding values and environmental components of the phenotype in a cohort at age $k.$ Assume that the joint distribution of $b_k$ and $e_k$ is bivariate normal with respective variances $\gamma^2_{a,k}$ and $\gamma^2_{e,k}$ and correlation $\rho_k$ as defined by equations 4-6 in the main text. The joint distribution of $b_{k+1}$ and $e_{k+1}$ after selection is
% NOTE: reference to equations here are hard-coded

\begin{align*}
& p(b_k,e_k) W(b_k + e_k) \\
    \propto & \exp{ \left[ - \frac{1}{2} \left( \frac{(b - \bar{b}_k)^2}{(1 - \rho_k^2) \gamma^2_{a,k}} - \frac{2 \rho_k (be - b\bar{e}_k - e \bar{e}_k + \bar{b}_k \bar{e}_k)}{(1-\rho_k^2) \gamma_{a,k} \gamma_{e,k}} + \frac{(e - \bar{e}_k)^2}{(1 - \rho_k^2) \gamma^2_{e,k}} + (b+e)^2 \right)  \right]} \\
    = & \exp{ \left[ - \frac{1}{2} \left( \frac{b^2 - 2b\bar{b}_k}{(1 - \rho_k^2) \gamma^2_{a,k}} + b^2 - \frac{2 \rho_k (be - e \bar{b}_k - b \bar{e}_k)}{(1-\rho_k^2) \gamma_{a,k} \gamma_{e,k}} + 2be + \frac{e^2 - 2e \bar{e}_k}{(1 - \rho_k^2) \gamma^2_{e,k}} + e^2 \right)  \right]} \\
    & \exp{ \left[ -\frac{1}{2} \left( \frac{\bar{b}_k^2}{(1 - \rho^2_k) \gamma^2_{a,k}} - \frac{2 \rho_k \bar{b}_k \bar{e}_k}{(1 - \rho^2_k)\gamma_{a,k} \gamma_{e,k}} + \frac{\bar{e}_k^2}{(1 - \rho^2_k)\gamma^2_{e,k}} \right) \right]} \\
    \propto & \exp{ \left[ - \frac{1}{2} \left( \frac{b^2 (1 + (1-\rho^2_k)\gamma^2_{a,k}) - 2b\bar{b}_k}{(1 - \rho_k^2) \gamma^2_{a,k}} + \frac{e^2 + (1 + (1 - \rho_k^2) \gamma^2_{e,k}) - 2 e \bar{e}_k}{(1 - \rho_k^2) \gamma^2_{e,k}} \right)  \right]} \\
    & \exp{ \left[ -\frac{1}{2} \left( \frac{2 be (\rho_k - (1 -\rho_k^2) \gamma_{a,k} \gamma_{e,k}) - \rho_k e \bar{b}_k - \rho_k b \bar{e}_k)}{(1-\rho_k^2) \gamma_{a,k} \gamma_{e,k}} \right) \right] } \\
    = & \exp{ \left[ - \frac{1}{2} \left( \frac{b^2 - 2b \frac{\bar{b}_k}{1 + (1-\rho^2_k)\gamma^2_{a,k}}} {\frac{(1 - \rho_k^2) \gamma^2_{a,k}}{1 + (1-\rho^2_k)\gamma^2_{a,k}}} + \frac{e^2 - 2 e \frac{\bar{e}_k}{1 + (1 - \rho_k^2) \gamma^2_{e,k}}}{\frac{(1 - \rho_k^2) \gamma^2_{e,k}}{1 + (1 - \rho_k^2) \gamma^2_{e,k}}} \right)  \right]} \\
    & \exp{ \left[ -\frac{1}{2} \left( \frac{2 \left( be - e \frac{ \bar{b}_k}{\rho_k - (1 -\rho_k^2) \gamma_{a,k} \gamma_{e,k}} - b \frac{ \bar{e}_k}{\rho_k - (1 -\rho_k^2) \gamma_{a,k} \gamma_{e,k}} \right)}{\frac{(1-\rho_k^2) \gamma_{a,k} \gamma_{e,k}}{\rho_k - (1 -\rho_k^2) \gamma_{a,k} \gamma_{e,k}}} \right) \right] }.
\end{align*}

\noindent We can use the denominators of the three respective terms above to construct the following relationships:

\begin{align*}
    (1 - \rho^2_{k+1}) \gamma^2_{a,k+1} & = \frac{(1 - \rho^2_k) \gamma^2_{a,k}}{1 + (1 - \rho^2_k) \gamma^2_{a,k}} \\
    (1 - \rho^2_{k+1}) \gamma^2_{e,k+1} & = \frac{(1 - \rho^2_k) \gamma^2_{e,k}}{1 + (1 - \rho^2_k) \gamma^2_{e,k}}.
\end{align*}

\noindent The square root of the product of these two terms is

\begin{align*}
    (1 - \rho^2_{k+1}) \gamma_{a,k+1} \gamma_{e,k+1} & = \frac{(1 - \rho^2_k) \gamma_{a,k} \gamma_{e,k}}{\sqrt{(1 + (1 - \rho_k^2)\gamma^2_{a,k})(1 + (1 - \rho^2_k) \gamma^2_{e,k})}}.
\end{align*}

\noindent We can use these to solve for $\rho_{k+1}$ as follows:

\begin{align*}
    \rho_{k+1} &= \frac{\rho_k - (1 - \rho^2_k)\gamma_{a,k} \gamma_{e,k}}{(1 - \rho^2_k)\gamma_{a,k} \gamma_{e,k}} \left( (1 - \rho^2_{k+1}) \gamma_{a,k+1} \gamma_{e,k+1} \right) \\
    \rho_{k+1} &= \frac{\rho_k - (1 - \rho^2_k)\gamma_{a,k} \gamma_{e,k}}{(1 - \rho^2_k)\gamma_{a,k} \gamma_{e,k}} \frac{(1 - \rho^2_k) \gamma_{a,k} \gamma_{e,k}}{\sqrt{(1 + (1 - \rho_k^2)\gamma^2_{a,k})(1 + (1 - \rho^2_k) \gamma^2_{e,k})}} \\
    & = \frac{\rho_k - (1 - \rho^2_k) \gamma_{a,k} \gamma_{e,k}}{\sqrt{(1 + (1 - \rho_k^2)\gamma^2_{a,k})(1 + (1 - \rho^2_k) \gamma^2_{e,k})}}.
\end{align*}

\noindent The bivariate normal distribution is proportional to the following term:

\begin{align*}
    \exp{ \left[ -\frac{1}{2} \left( \frac{(b - \bar{b}_{k+1})^2}{(1 - \rho^2_{k+1}) \gamma^2_{a,k+1}} - \frac{2 (be - b\bar{e}_{k+1} - e \bar{b}_{k+1} + \bar{b}_{k+1}\bar{e}_{k+1})}{ \frac{(1-\rho^2_{k+1}) \gamma_{a,k+1} \gamma_{e,k+1}}{\rho_{k+1}}} + \frac{(e - \bar{e}_{k+1})^2}{(1-\rho^2_{k+1})^2 \gamma^2_{e,k+1}} \right) \right] }.
\end{align*}

\noindent The two remaining unknowns are $\bar{b}_{k+1}$ and $\bar{e}_{k+1}.$ We can solve for them by constructing a system of equations as follows:

\begin{align}
    \label{eq:02_supp_mvnorm2a}
    - \frac{2 b \bar{b}_{k+1}}{(1 - \rho^2_{k+1}) \gamma^2_{a,k}} + \frac{2b \bar{e}_{k+1}}{\frac{(1-\rho^2_{k+1}) \gamma_{a,k+1} \gamma_{e,k+1}}{\rho_{k+1}}} & = - \frac{2b\frac{\bar{b}_k}{1 + (1 - \rho^2_k) \gamma^2_{a,k}}}{(1 - \rho^2_{k+1}) \gamma^2_{a,k+1}} + \frac{2b \frac{\bar{e}_k}{\rho_k - (1 - \rho^2_k) \gamma_{a,k} \gamma_{e,k}}}{\frac{(1 - \rho^2_{k+1})\gamma_{a,k+1} \gamma_{e,k}}{\rho_{k+1}}} \\
    \label{eq:02_supp_mvnorm2b}
    \frac{2 e \bar{b}_{k+1}}{\frac{(1 - \rho_{k+1}) \gamma_{a,k+1} \gamma_{e,k+1}}{\rho_{k+1}}} - \frac{2 e \bar{e}_{k+1}}{(1 - \rho_{k+1}^2) \gamma^2_{e,k+1}} & = -\frac{2e\frac{\bar{e}_k}{1+(1-\rho^2_k)\gamma^2_{e,k}}}{(1 - \rho^2_{k+1})\gamma^2_{e,k+1}} + \frac{2e \frac{\bar{b}_k \rho_k}{\rho_k (1 - \rho_k^2) \gamma_{a,k} \gamma_{e,k}}}{\frac{(1 - \rho_{k+1}^2) \gamma_{a,k+1} \gamma_{e,k+1}}{\rho_{k+1}}}.
\end{align}

\noindent With algebra and cancellation of like-terms, the above equations become

\begin{align*}
    \bar{b}_{k+1} - \rho_{k+1} \frac{\gamma_{a,k+1}}{\gamma_{e,k+1}} \bar{e}_{k+1} & = \frac{\bar{b}_k}{1 + (1 - \rho^2_k) \gamma^2_{a,k}} - \rho_{k+1} \frac{\gamma_{a,k+1}}{\gamma_{e,k+1}} \frac{\bar{e}_k \rho_k}{\rho_k - (1 - \rho^2_k) \gamma_{a,k} \gamma_{e,k}}  \\
    \bar{e}_{k+1} - \rho_{k+1} \frac{\gamma_{e,k+1}}{\gamma_{a,k+1}} \bar{b}_{k+1} & = \frac{\bar{e}_k}{1 + (1 - \rho_k^2) \gamma^2_{e,k}} - \rho_{k+1} \frac{\gamma_{e,k+1}}{\gamma_{a,k+1}} \frac{\bar{e}_k \rho_k}{\rho_k - (1 - \rho_k^2) \gamma_{a,k} \gamma_{e,k}} .
\end{align*}

\noindent We can solve for $\bar{e}_{k+1}$ in terms of $\bar{b}_{k+1}$ in the latter of these equations, producing

\begin{align*}
    \bar{e}_{k+1} &= \frac{\bar{e}_{k+1}}{1 + (1 - \rho^2_k)\gamma^2_{e,k}} + \rho_{k+1} \frac{\gamma_{e,k+1}}{\gamma_{a,k+1}} \left( \bar{b}_{k+1} - \frac{\bar{b}_k \rho_k}{\rho_k - (1 - \rho^2_k) \gamma_{a,k} \gamma_{e,k}} \right).
\end{align*}

\noindent Substituting this into the first of the equations gives

\begin{align*}
    \bar{b}_{k+1} = & \frac{\bar{b}_k}{1 + (1 - \rho^2_k)\gamma^2_{a,k}} - \rho_{k+1} \frac{\gamma_{a,k+1}}{\gamma_{e,k+1}} \frac{\bar{e}_k \rho_k}{\rho_k - (1 - \rho^2_k) \gamma_{a,k} \gamma_{e,k}} + \\
    & \rho_{k+1} \frac{\gamma_{a,k+1}}{\gamma_{e,k+1}} \left( \frac{\bar{e}_k}{1 + (1 - \rho^2_k) \gamma^2_{e,k}} + \rho_{k+1} \frac{\gamma_{e,k+1}}{\gamma_{a,k+1}} \left( \bar{b}_{k+1} - \frac{\bar{b}_k \rho_k}{\rho_k - (1 - \rho^2_k) \gamma_{a,k} \gamma_{e,k}} \right) \right) \\
    = & \frac{\bar{b}_k}{1 + (1 - \rho^2_k) \gamma^2_{a,k}} - \rho_{k+1} \frac{\gamma_{a,k+1}}{\gamma_{e,k+1}} \bar{e}_k \left( \frac{\rho_k}{\rho_k - (1 - \rho^2_k) \gamma_{a,k} \gamma_{e,k}} - \frac{1}{1 + (1 - \rho^2_k)\gamma^2_{e,k}} \right) \\
    & + \rho^2_{k+1} \left(\bar{b}_{k+1} -  \frac{\bar{b}_k \rho_k}{\rho_k - (1 - \rho^2_k) \gamma_{a,k} \gamma_{e,k}} \right). \\
    \bar{b}_{k+1} (1 - \rho^2_{k+1}) = & \bar{b}_k \left( \frac{1}{1 + (1 - \rho^2_k)} - \rho^2_{k+1} \frac{\rho_k^2}{\rho_k - (1 - \rho^2_k) \gamma_{a,k} \gamma_{e,k}} \right) \\
    & - \rho_{k+1} \frac{\gamma_{a,k+1}}{\gamma_{e,k+1}} \bar{e}_k \left( \frac{\rho_k}{\rho_k - (1 - \rho^2_k) \gamma_{a,k} \gamma_{e,k}} - \frac{1}{1 + (1 - \rho^2_k)\gamma^2_{e,k}} \right)
\end{align*}

\noindent The term with $\bar{b}_k$ can be rewritten as:

\begin{align*}
    & \bar{b}_k \left( \frac{1}{1 + (1 - \rho^2_k) \gamma^2_{a,k}} - \rho^2_{k+1} \frac{\rho_k}{\rho_k - (1 - \rho^2_k) \gamma_{a,k} \gamma_{e,k}} \right) \\
    = & \bar{b}_k \left( \frac{1}{1 + (1 - \rho^2_k) \gamma^2_{a,k}} - \frac{(\rho^2 - (1 - \rho^2_k) \gamma_{a,k} \gamma_{e,k})^2}{(1 + (1 - \rho^2_k)\gamma^2_{a,k})(1 + (1 - \rho^2_k)\gamma^2_{e,k})} \frac{\rho_k}{\rho_k - (1 - \rho^2_k) \gamma_{a,k} \gamma_{e,k}} \right) \\ 
    = & \bar{b}_k \left( \frac{1}{1 + (1 - \rho^2_k) \gamma^2_{a,k}} - \frac{\rho_k^2 - (1 - \rho^2_k) \rho_k \gamma_{a,k} \gamma_{e,k}}{(1 + (1 - \rho^2_k) \gamma^2_{a,k})(1 + (1 - \rho^2_k) \gamma^2_{e,k})} \right) \\
    = & \bar{b}_k \left( \frac{1 + (1 - \rho^2_k) \gamma^2_{e,k} - \rho_k^2 + (1 - \rho^2_k) \rho_k \gamma_{a,k} \gamma_{e,k}}{(1 + (1 - \rho^2_k) \gamma^2_{a,k})(1 + (1 - \rho^2_k) \gamma^2_{e,k})} \right) \\
    = & \bar{b}_k \left( \frac{(1 - \rho^2_k) (1 + \gamma^2_{e,k} + \rho_k \gamma_{a,k} \gamma_{e,k})}{(1 + (1 - \rho^2_k) \gamma^2_{a,k})(1 + (1 - \rho^2_k) \gamma^2_{e,k})} \right).
\end{align*}

Using the definitions of $(1 - \rho^2_{k+1}) \gamma^2_{a,k+1}$ and $(1 - \rho^2_{k+1}) \gamma^2_{e,k+1}$ above, the terms $\gamma^2_{a,k+1}$ and $\gamma^2_{e,k+1}$ are

\begin{align*}
    & \gamma^2_{a,k+1} = \frac{\gamma^2_{a,k} (1 + (1 - \rho^2_k) \gamma^2_{e,k})}{1 + \gamma^2_{a,k} + \gamma^2_{e,k} + 2 \rho_k \gamma_{a,k} \gamma_{e,k}}, & \gamma^2_{e,k+1} = \frac{\gamma^2_{e,k} (1 + (1 - \rho^2_k) \gamma^2_{a,k})}{1 + \gamma^2_{a,k} + \gamma^2_{e,k} + 2 \rho_k \gamma_{a,k} \gamma_{e,k}}. & 
\end{align*}

\noindent With these, the term $\rho_{k+1} (\gamma_{a,k+1} / \gamma_{e,k+1})$ is

\begin{align*}
    \frac{\gamma_{a,k+1}}{\gamma_{e,k+1}} \rho_{k+1} & = \frac{\gamma_{a,k}}{\gamma_{e,k}} \sqrt{ \frac{1 + (1 - \rho^2_k) \gamma^2_{e,k}}{1 + (1 - \rho^2_k) \gamma^2_{a,k}}} \left( \frac{\rho_k - (1 - \rho_k^2)\gamma_{a,k} \gamma_{e,k}}{\sqrt{(1 + (1 - \rho^2_k) \gamma^2_{a,k})(1 + (1 - \rho^2_k) \gamma^2_{e,k})}} \right) \\
    & = \frac{\gamma_{a,k}}{\gamma_{e,k}} \left( \frac{\rho_k - (1 - \rho^2_k)\gamma_{a,k} \gamma_{e,k}}{1 + (1 - \rho^2_k) \gamma^2_{a,k}} \right).
\end{align*}

\noindent Substituting this into the term for $\bar{e}_k$ in the expression for $\bar{e}_{k+1}$ above gives

\begin{align*}
    & \bar{e}_k \rho_{k+1} \frac{\gamma_{a,k}}{\gamma_{e,k}} \left( \frac{\rho_k}{\rho_k - (1 - \rho^2_k) \gamma_{a,k} \gamma_{e,k}} - \frac{1}{1 + (1 - \rho^2_k) \gamma^2_{e,k}} \right) & = \\
    & \bar{e}_k \frac{\gamma_{a,k}}{\gamma_{e,k}} \left( \frac{\rho_k}{(1 + (1 - \rho^2_k)\gamma^2_{a,k}} - \frac{\rho_k - (1 - \rho^2_k)\gamma_{a,k}\gamma_{e,k}}{(1 + (1 - \rho^2_k)\gamma_{a,k}^2)(1 + (1 - \rho^2) \gamma^2_{e,k})} \right) & = \\
    & \bar{e}_k \frac{\gamma_{a,k}}{\gamma_{e,k}} \left( \frac{\rho_k + \rho_k(1-\rho_k^2)\gamma^2_{e,k} - \rho_k + (1 - \rho_k^2)\gamma_{a,k}\gamma_{e,k}}{(1 + (1-\rho^2_k)\gamma^2_{a,k})(1+(1-\rho^2_k)\gamma^2_{e,k})} \right) & = \\
    & \bar{e}_k \left( \frac{(1 - \rho^2_k)(\rho_k \gamma_{a,k}\gamma_{e,k} + \gamma^2_{a,k})}{(1 + (1-\rho^2_k)\gamma^2_{a,k})(1 + (1 - \rho^2_k)\gamma^2_{e,k})} \right).
\end{align*}

Combining these two terms gives us

\begin{align*}
    \bar{b}_{k+1} (1 - \rho^2_{k+1}) = (1 - \rho^2_k) \frac{\bar{b}_k  (1 + \gamma^2_{e,k} + \rho_k \gamma_{a,k} \gamma_{e,k}) - \bar{e}_k (\rho_k \gamma_{a,k} \gamma_{e,k} + \gamma^2_{a,k})}{(1 + (1 - \rho^2_k)\gamma^2_{a,k})(1 + (1-\rho^2_k)\gamma^2_{e,k})}.
\end{align*}

\noindent Using the fact that

\begin{align*}
    1 - \rho^2_{k+1} &= \frac{(1 - \rho^2_k) (1 + \gamma^2_{a,k} + \gamma^2_{e,k} + 2 \rho_k \gamma_{a,k} \gamma_{e,k})}{(1 + (1 - \rho^2_k) \gamma^2_{a,k}) (1 + (1 - \rho^2_k) \gamma^2_{e,k})},
\end{align*}

\noindent we can say

\begin{align*}
    \bar{b}_{k+1} &= \frac{\bar{b}_k  (1 + \gamma^2_{e,k} + \rho_k \gamma_{a,k} \gamma_{e,k}) - \bar{e}_k (\rho_k \gamma_{a,k} \gamma_{e,k} + \gamma^2_{a,k})}{1 + \gamma^2_{a,k} + \gamma^2_{e,k} + 2 \rho_k \gamma_{a,k} \gamma_{e,k}}.
\end{align*}

Using Equations \ref{eq:02_supp_mvnorm2a} and \ref{eq:02_supp_mvnorm2b} to solve for $\bar{b}_{k+1}$ is involved. Instead, consider that $\bar{z}_{k+1} = \bar{b}_{k+1} + \bar{e}_{k+1}.$ For a given $\bar{z}_k,$ $\bar{z}_{k+1} = \bar{z}_k / (1 + \gamma^2_k) = (\bar{b}_k + \bar{e}_k) / (1 + \gamma^2_{a,k} + \gamma^2_{e,k} + 2 \rho_k \gamma_{a,k} \gamma_{e,k}).$ Solving for $\bar{e}_{k+1}$ gives

\begin{align*}
    \bar{e}_{k+1} & = \bar{z}_{k+1} - \bar{b}_{k+1} \\
    & = \frac{\bar{b}_k + \bar{e}_k - \bar{b}(1 + \gamma^2_{e,k} + \rho_k \gamma_{a,k} \gamma_{e,k}) + \bar{e}_k (\rho_k \gamma_{a,k} \gamma_{e,k} + \gamma^2_{a,k})}{1 + \gamma^2_{a,k} + \gamma^2_{e,k} + 2 \rho_k \gamma_{a,k} \gamma_{e,k}} \\
    & = \frac{\bar{e}_k (1 + \gamma^2_{a,k} + \rho_k \gamma_{a,k} \gamma_{e,k}) - \bar{b}_k ( \gamma^2_{e,k} + \rho_k \gamma_{a,k} \gamma_{e,k})}{1 + \gamma^2_{a,k} + \gamma^2_{e,k} + 2 \rho_k \gamma_{a,k} \gamma_{e,k}}
\end{align*}

Variances and covariance are independent of $\theta$. Because of this, we can write these in terms of the variance of newborn cohorts, $\gamma^2_{a,0}$ and $\gamma^2_{e,0}$ as follows:

\begin{align*}
    \bar{b}_{k+1} &= \frac{\bar{b}_k \left( 1 + \frac{\gamma^2_{e,0}}{1 + k(\gamma^2_{a,0} + \gamma^2_{e,0})} \right) - \bar{e}_{k} \frac{\gamma^2_{a,0}}{1 + k(\gamma^2_{a,0} + \gamma^2_{e,0})}}{1 + \frac{\gamma^2_{a,0} + \gamma^2_{e,0}}{1 + k(\gamma^2_{a,0} + \gamma^2_{e,0})}} \\
    &= \frac{\bar{b}_k \left(1 + k(\gamma^2_{a,0} + \gamma^2_{e,0}) + \gamma^2_{e,0}\right) - \bar{e}_k \gamma^2_{a,0}}{1 + (k+1)(\gamma^2_{a,0} + \gamma^2_{e,0})} \\
    &= \bar{b}_k - \frac{(\bar{b}_k + \bar{e}_k) \gamma^2_{e,0}}{1 + (k+1)(\gamma^2_{a,0} + \gamma^2_{e,0})} \\
    \bar{e}_{k+1} &= \frac{\bar{e}_k \left(1 + \frac{\gamma^2_{a,0}}{1 + k(\gamma^2_{a,0} + \gamma^2_{e,0})} \right) - \bar{b}_k \frac{\gamma^2_{e,0}}{1 + k(\gamma^2_{a,0} + \gamma^2_{e,0})}}{1 + \frac{\gamma^2_{a,0} + \gamma^2_{e,0}}{1 + k(\gamma^2_{a,0} + \gamma^2_{e,0})}} \\
    &= \frac{\bar{e}_k \left( 1 + k(\gamma^2_{a,0} + \gamma^2_{e,0}) + \gamma^2_{a,0}\right) - \bar{b}_k \gamma^2_{e,0}}{1 + (k+1)(\gamma^2_{a,0} + \gamma^2_{e,0})} \\
    &= \bar{e}_k - \frac{(\bar{b}_k + \bar{e}_k)\gamma^2_{e,0}}{1 + (k+1)(\gamma^2_{a,0} + \gamma^2_{e,0})}.
\end{align*}

This recursion relationship is solved by the following terms:

\begin{align*}
    \bar{b}_{k+t,t} &= \bar{b}_k - \frac{t (\bar{b}_k + \bar{e}_k) \gamma^2_{e,0}}{1 + (k+t)(\gamma^2_{a,0} + \gamma^2_{e,0})} \\
    \bar{e}_{k+t,t} &= \bar{e}_k - \frac{t (\bar{b}_k + \bar{e}_k) \gamma^2_{a,0}}{1 + (k+t)(\gamma^2_{a,0} + \gamma^2_{e,0})}.
\end{align*}

\noindent This can be shown by induction. Let $\bar{z}_k = \bar{b}_k + \bar{e}_k.$ Assume the relationship is true for all $t \geq 0.$ Then, using the recursion relationship,

\begin{align*}
    \bar{b}_{k+t+1,t+1} = & \frac{\left(\bar{b}_k - \frac{t\bar{z}_k \gamma^2_{e,0}}{1 + (k+t)(\gamma^2_{a,0} + \gamma^2_{e,0})}\right) \left( 1 + (k+t+1)(\gamma^2_{a,0} + \gamma^2_{e,0}) \right)}{{1 + (k+t+1)(\gamma^2_{a,0} + \gamma^2_{e,0})}} - \\
    & \frac{\gamma^2_{e,0} \left( \bar{b}_k - \frac{t\bar{z}_k \gamma^2_{a,0}}{1 + (k+t)(\gamma^2_{a,0} + \gamma^2_{e,0})}\right) - \gamma^2_{e,0} \left( \bar{e}_k - \frac{t \bar{z}_k \gamma^2_{e,0}}{1 + (k+t)(\gamma^2_{a,0} + \gamma^2_{e,0})} \right)}{1 + (k+t+1)(\gamma^2_{a,0} + \gamma^2_{e,0})} \\
    = & \bar{b}_k - \frac{t\bar{z}_k \gamma^2_{e,0}}{1 + (k+t)(\gamma^2_{a,0} + \gamma^2_{e,0})} - \\ 
    & \frac{\gamma^2_{e,0}}{1 + (k+t+1)(\gamma^2_{a,0} + \gamma^2_{e,0})} \left( \bar{b}_k + \bar{e}_k - \frac{t\bar{z}_k (\gamma^2_{a,0} + \gamma^2_{e,0})}{1 + (k+t)(\gamma^2_{a,0} + \gamma^2_{e,0})} \right) \\
    = & \bar{b}_k - \frac{t \bar{z}_k \gamma^2_{e,0}}{1 + (k+t)(\gamma^2_{a,0} + \gamma^2_{e,0})} - \\
    & \frac{\bar{z}_k \gamma^2_{e,0}}{1 + (k+t+1)(\gamma^2_{a,0} + \gamma^2_{e,0})} \left(1 - \frac{t(\gamma^2_{a,0} + \gamma^2_{e,0})}{1 + (k+t)(\gamma^2_{a,0} + \gamma^2_{e,0})} \right) \\
    = & \bar{b}_k - \frac{t \bar{z}_k \gamma^2_{e,0}}{1 + (k+t)(\gamma^2_{a,0} + \gamma^2_{e,0})} - \\
    & \frac{\bar{z}_k \gamma^2_{e,0}}{1 + (k+t+1)(\gamma^2_{a,0} + \gamma^2_{e,0})} \frac{1 + k(\gamma^2_{a,0} + \gamma^2_{e,0})}{1 + (k+t)(\gamma^2_{a,0} + \gamma^2_{e,0})} \\
    = & \bar{b}_k - \frac{\bar{z}_k \gamma^2_{e,0}}{1 + (k+t)(\gamma^2_{a,0} + \gamma^2_{e,0})} \left(t + \frac{1 + k(\gamma^2_{a,0} + \gamma^2_{e,0})}{1 + (k+t+1)(\gamma^2_{a,0} + \gamma^2_{e,0})} \right) \\
    = & \bar{b}_k - \frac{\bar{z}_k \gamma^2_{e,0}}{1 + (k+t)(\gamma^2_{a,0} + \gamma^2_{e,0})} \left( \frac{(t + 1) + (k(t+1)+t(t+1))(\gamma^2_{a,0} + \gamma^2_{e,0})}{1 + (k+t+1) (\gamma^2_{a,0} + \gamma^2_{e,0})} \right) \\
    = & \bar{b}_k - \frac{(\bar{b}_k + \bar{e}_k) \gamma^2_{e,0}}{1 + (k+t+1)(\gamma^2_{a,0} + \gamma^2_{e,0})}
\end{align*}

\begin{align*}    
    \bar{e}_{k+t+1,t+1} = & \frac{\left( \bar{e}_k - \frac{t \bar{z}_k \gamma^2_{a,0}}{1 + (k+t)(\gamma^2_{a,0} + \gamma^2_{e,0})} \right)(1 + (k+t+1)(\gamma^2_{a,0} + \gamma^2_{e,0}))}{1 + (k+t+1)(\gamma^2_{a,0} + \gamma^2_{e,0}))} - \\
    & \frac{\gamma^2_{a,0} \left(\bar{b}_k - \frac{t\bar{z}_k \gamma^2_{e,0}}{1 + (k+t)(\gamma^2_{a,0} + \gamma^2_{e,0})}\right) - \gamma^2_{a,0} \left(\bar{e}_k - \frac{t \bar{z}_k \gamma^2_{a,0}}{1 + (k+t)(\gamma^2_{a,0} + \gamma^2_{e,0})}\right)}{1 + (k+t+1)(\gamma^2_{a,0} + \gamma^2_{e,0})} \\
    = & \bar{e}_k - \frac{t \bar{z}_k \gamma^2_{a,0}}{1 + (k+t)(\gamma^2_{a,0} + \gamma^2_{e,0})} - \\
    & \frac{\gamma^2_{a,0}}{1 + (k+t+1)(\gamma^2_{a,0} + \gamma^2_{e,0})} \left( \bar{b}_k + \bar{e}_k - \frac{t \bar{z}_k (\gamma^2_{a,0} + \gamma^2_{e,0})}{1 + (k+t)(\gamma^2_{a,0} + \gamma^2_{e,0})} \right) \\
    = & \bar{e}_k - \frac{t \bar{z}_k \gamma^2_{a,0}}{1 + (k+t)(\gamma^2_{a,0} + \gamma^2_{e,0})} - \\
    & \frac{\bar{z}_k \gamma^2_{e,0}}{1 + (k+t+1)(\gamma^2_{a,0} + \gamma^2_{e,0})} \left( 1 - \frac{t(\gamma^2_{a,0} + \gamma^2_{e,0})}{1 + (k+t)(\gamma^2_{a,0} + \gamma^2_{e,0})} \right) \\
    = & \bar{e}_k - \frac{t \bar{z}_k \gamma^2_{a,0}}{1 + (k+t)(\gamma^2_{a,0} + \gamma^2_{e,0})} - \\
    & \frac{\bar{z}_k \gamma^2_{e,0}}{1 + (k+t+1)(\gamma^2_{a,0} + \gamma^2_{e,0})} \frac{1 + k(\gamma^2_{a,0} + \gamma^2_{e,0})}{1 + (k+t)(\gamma^2_{a,0} + \gamma^2_{e,0})} \\
    = & \bar{e}_k - \frac{\bar{z}_k \gamma^2_{e,0}}{1 + (k+t)(\gamma^2_{a,0} + \gamma^2_{a,0})} \left( t + \frac{1 + k(\gamma^2_{a,0} + \gamma^2_{e,0})}{1 + (k+t+1)(\gamma^2_{a,0} + \gamma^2_{e,0})} \right) \\
    = & e_k - \frac{(t+1)\bar{z}_k \gamma^2_{a,0}}{1 + (k+t)(\gamma^2_{a,0} + \gamma^2_{e,0})} \left( \frac{(t+1) + \left( k(t+1) + t(t+1) \right)(\gamma^2_{a,0} + \gamma^2_{e,0})}{1 + (k+t+1)(\gamma^2_{a,0} + \gamma^2_{e,0})} \right) \\
    = & \bar{e}_k - \frac{(t+1)(\bar{b}_k + \bar{e}_k)\gamma^2_{a,0}}{1 + (k+t+1)(\gamma^2_{a,0} + \gamma^2_{e,0})}.
\end{align*}

For the specific case in our model where $\bar{b}_{k,0} = \theta,$ $\bar{e}_k = 0$ for all $k,$ the recursion relationships for $\bar{b}_{k+t,t}$ and $\bar{e}_{k+t,t}$ can be solved by

\begin{align*}
    \bar{b}_{k+t,t} &= \theta \left( 1 - \frac{t \gamma^2_{a,0}}{1 + (k+t)(\gamma^2_{a,0} + \gamma^2_{e,0})} \right) \\
    \bar{e}_{k+t,t} &= - \frac{\theta t \gamma^2_{e,0}}{1 + (k+t)(\gamma^2_{a,0} + \gamma^2_{e,0})},
\end{align*}

\noindent for all $t \geq 0.$

\clearpage

%%%%%%%%%%%%%%%%%%%%%%%%%%%%%%%%%%%%%%%%%%%%%%%%%

\section{Numerical estimation of $\lambda$}

For $z \neq 0$ (i.e., a population not perfectly adapted to the environment), the mean survival in the population is the cohort-weighted mean of cohort survivals. It follows from Lande (1976) (p. 322, integrating the mean of a Gaussian fitness function over a normally distributed population of phenotypes) and Equation 1 in main text that the mean survival in the population is

\begin{align}
    \label{eq:02_supp_sbar}
    \bar{s} = \displaystyle\sum_{k=0}^\infty p_k \left( \widehat{s} \sqrt{ \frac{1 + k \gamma^2_0}{1 + (k+1)\gamma^2_0}} \right) \left( \exp{ \left( - \frac{\bar{z}^2_k}{2} \right)} \right),
\end{align}

\noindent where $\bar{z}_k$ is the mean population phenotype in cohort $k.$ The right hand side of Equation~\ref{eq:02_supp_sbar} can be substituted into the expression $\lambda = \bar{s} (1 + r).$ Pulling out the common $\widehat{s}$ term from the summation and using $\widehat{\lambda} = \widehat{s} (1 + r)$ produces

\begin{align*}
    \lambda = \widehat{\lambda} \displaystyle\sum_{k=0}^\infty p_k \left( \sqrt{\frac{1 + k \gamma^2_0}{1 + (k+1)\gamma^2_0}} \right) \left( \exp{ \left( - \frac{\bar{z}_k^2}{2} \right)} \right).
\end{align*}

We assume in generating Fig. 1 (main text) that the population remains at equilibrium, with $p_k$ set to the stable stage distribution (see Appendix~\ref{sec:02_stable_age_dist}) and that $\bar{z}_k = \bar{z}$ for all cohorts. We approximated $\bar{s}$ and $\lambda$ by taking the sum over $k$ from zero to $100,000.$ Details on numerical estimation of $\gamma^2_0$ and $\lambda^*$ in the stable stage distribution are provided in Appendix~\ref{sec:02_supp_newton}.

For small $\gamma^2,$ mean survival can be approximated by

\begin{align*}
    \bar{s} \approx \widehat{s} \left( \frac{1}{\sqrt{1 + \gamma^2}} \right) \left( \exp{ \left( - \frac{\bar{z}^2}{2} \right)}  \right),
\end{align*}

\noindent  and this value can be used to approximate $\lambda$ accordingly. The first parenthesized term is the ``variance load'' (Lande and Shannon (1996), Ashander et al. (2016))) where mean population fitness is decreased as populations deviate further from the environmental optimum, and the second parenthesized term is the maladaptation load. However, this approximation fails as $\gamma^2$ grows large, producing error on the order of $0.1$ for $\gamma^2 = 0.4,$ the highest value used in our simulation experiment.

\clearpage

%%%%%%%%%%%%%%%%%%%%%%%%%%%%%%%%%%%%%%%%%%%%%%%%%

\section{Comparison with Gomulkiewicz and Holt's (1995) model}
\label{sec:02_g_and_h}

The model of Gomulkiewicz and Holt (1995) assumes a population with life history lasting at most one time step adapting at a constant rate to a constant environment with a constant degree of phenotypic variance. In their model, the log of the population size at time $t$ (denoted $n_t$) is
% NOTE: reference here is hard coded

\begin{align}
\label{eq:02_gh_orig}
    n_t = n_0 + t \left( \log{\widehat{W}} + \frac{1}{2} \log{ \left( \frac{\omega^2}{\sigma^2 + \omega^2} \right)} \right) - \frac{z_0^2}{2(\omega^2 + \sigma^2)} \left( \frac{1 - k^{2t}}{1 - k^2} \right),
\end{align}

\noindent where $n_0$ is the log of the initial population size, $k = (\omega^2 + (1-h^2)\sigma^2)/(\omega^2 + \sigma^2)$ is the rate of phenotypic adaptation, and all other variables are defined the same as in our main text. The term $\log{\widehat{W}}$ is the maximum rate of population increase per time step, from which the variance and maladaptation loads are subtracted to reduce population growth rates; because the model assumes generations last one time step, $\widehat{\lambda} = \widehat{W}.$ 

Substituting in our more general form of maximum population growth rate ($\widehat{\lambda} = \widehat{W}(1 - \widehat{s}) + \widehat{s}$), and adopting our convention of rescaling phenotypes (see main text) gives

\begin{align}
    \label{eq:02_gh_longevity}
    n_t = n_0 + t \left( \log{\left( \widehat{W}(1 - \widehat{s})  + \widehat{s} \right)} - \frac{1}{2} \log{(1+\gamma^2)} \right) - \frac{(z_0 / \omega)^2}{2(1 + \gamma^2)} \left( \frac{1 - k^{2t}}{1 - k^t} \right).
\end{align}

\noindent The exponentiated form of the right hand side appears in Figure 1b of our main text as $N_t$. Setting $\widehat{s} = 0$ reproduces Equation~\ref{eq:02_gh_orig}. The expressions in~\ref{eq:02_gh_orig} and~\ref{eq:02_gh_longevity} make the simplifying assumption that $k$ is held constant over time. 
% NOTE: reference to figure here is hard coded

\clearpage

%%%%%%%%%%%%%%%%%%%%%%%%%%%%%%%%%%%%%%%%%%%%%%%%%

\section{Model validation}
\label{sec:02_supp_validation}

We validated analytical expressions described in the main text and supplements by comparing expressions to simulation output. Our simulation model is described in the Methods section of the main text. We ran simulations both at steady state ($\bar{z}_0 = \theta = 0$) and following an environmental change ($\bar{z}_0 = 1$). We ran 20 simulations per parameter combination for the steady state visualizations and 100 simulations per parameter combination for the post-change visualizations, using the same parameter combinations as used for the simulated experiment as described in the main text, with the following exceptions. All simulations were run at initial size $N_0 = 5000$ (rather than $N_0 = 20000$ as in the main experiments) for computational ease. For simulations following environmental change, and simulations were run only for $h^2 = 0.5$ to reduce the complexity of visualizations while still highlighting non-trivial dynamics of all variables. Simulations in the constant environment were run for ten time steps, to show that expected values are similar across time, while simulations in the changed environment were run for three time steps to show the accuracy of expressions for dynamics following environmental change.

\subsection{Steady state validation}

For population-level variables (Figures~\ref{fig:02_supp_age_validate} --~\ref{fig:02_supp_zbar_validate}), we present results for each trial in each time step. For age-level variables (Figures~\ref{fig:02_supp_bvar_age_validate} -- ~\ref{fig:02_supp_zvar_age_validate}), we present time-specific means by aggregating across all trials within a parameter combination for the given age and time step. We consider plots with overlapping lines to be evidence that simulated populations are remaining at steady state. Age-specific variable plots do not include the initial time step ($t = 0$) because this would only demonstrate that populations are initialized as expected. Variances presented are population-level variances, not sample variances, i.e., they are not Bessel-corrected. Because of this, variances of zero may reflect multiple individuals with values identical to the age-class mean, but may also represent cohorts represented by a single individual (which trivially has the cohort mean). Thus, because these simulations are of finite populations, age-classes that are relatively low-density may have an excess of zeros due to having only one individual in them. We only plot group-means estimated from more than three observations in order to focus on precise estimates.

\begin{figure}[!htp]
    \includegraphics[width=0.95\textwidth]{figs/fig_v2_age_full.png}
    \caption{Age distribution at steady state. Each line is the mean age distribution for a time step, averaged across 20 trials per parameter combination. Circles are analytical expectations, as defined by Equation 2 in the main text.}
    \label{fig:02_supp_age_validate}
\end{figure}

\begin{figure}[!htp]
    \includegraphics[width=0.95\textwidth]{figs/fig_v3_age_0-9.png}
    \caption{Age distribution at steady state, shown only for ages zero through nine. Each line is the mean age distribution for a time step, averaged across 20 trials per parameter combination. Circles are analytical expectations, as defined by Equation 2 in the main text.}
    % NOTE: figure ref is hard coded
\end{figure}

\begin{figure}[!htp]
    \includegraphics[width=0.95\textwidth]{figs/fig_v4_bbar_steady.png}
    \caption{Mean population breeding value ($\bar{b}_t$) at steady state for 20 trials per parameter combination.}
\end{figure}

\begin{figure}[!htp]
    \includegraphics[width=0.95\textwidth]{figs/fig_v5_ebar_steady.png}
    \caption{Mean population environmental component to phenotypes ($\bar{e}_t$) at steady state for 20 trials per parameter combination.}
\end{figure}

\begin{figure}[!htp]
    \includegraphics[width=0.95\textwidth]{figs/fig_v6_zbar_steady.png}
    \caption{Mean population phenotype ($\bar{z}_t$) at steady state for 20 trials per parameter combination.}
    \label{fig:02_supp_zbar_validate}
\end{figure}

\begin{figure}[!htp]
    \includegraphics[width=0.95\textwidth]{figs/fig_v7_bvar_age_steady.png}
    \caption{Mean additive genetic variance for each age class ($\gamma^2_{a,k}$) at steady state, shown for ages zero through nine. Each line shows the additive genetic variance in one time step, averaged across 20 trials per parameter combination. Circles give the analytical expression defined by Equation 4 in the main text.}
    \label{fig:02_supp_bvar_age_validate}
\end{figure}

\begin{figure}[!htp]
    \includegraphics[width=0.95\textwidth]{figs/fig_v8_evar_age_steady.png}
    \caption{Mean variance of environmental contribution to phenotype for each age class ($\gamma^2_{e,k}$) at steady state, shown for ages zero through nine. Each line shows the environmental contribution to phenotypic variance in one time step, averaged across 20 trials per parameter combination. Circles give the analytical expression defined by Equation 5 in the main text.}
\end{figure}

\begin{figure}[!htp]
    \includegraphics[width=0.95\textwidth]{figs/fig_v9_zvar_age_steady.png}
    \caption{Mean phenotypic variance for each age class ($\gamma^2_k$) at steady state, shown for ages zero through nine. Each line shows the phenotypic variance in one time step, averaged across 20 trials per parameter combination. Circles give the analytical expression defined in the main text.}
    \label{fig:02_supp_zvar_age_validate}
\end{figure}

\subsection{Following environmental change}

Figures~\ref{fig:02_supp_bbar_change_validate} --~\ref{fig:02_supp_zbar_change_validate} demonstrate cohort mean phenotypic component values over time following environmental change, for cohorts extant at the time of the environmental change. Figures~\ref{fig:02_supp_bvar_change_validate} --~\ref{fig:02_supp_rho_change_validate} show age-specific phenotypic component variances and correlations following environmental change; overlapping lines demonstrate the stability of component variances following environmental changes.

\begin{figure}[!htp]
    \includegraphics[width=0.95\textwidth]{figs/fig_v11_bbar_change.png}
    \caption{Mean breeding value of cohorts for three time steps following environmental change. One line represents one cohort. For visual clarity, only cohorts age zero through four at the time of the environmental shift for which there were more than 10 observations used to estimate the mean are shown. Circles give the analytical expression defined by Equation 8 in the main text.}
    \label{fig:02_supp_bbar_change_validate}
\end{figure}

\begin{figure}[!htp]
    \includegraphics[width=0.95\textwidth]{figs/fig_v12_ebar_change.png}
    \caption{Mean environmental component of phenotype within cohorts for three time steps following environmental change. One line represents one cohort. For visual clarity, only cohorts age zero through four at the time of the environmental shift for which there were more than 10 observations used to estimate the mean are shown. Circles give the analytical expression defined by Equation 9 in the main text.}
\end{figure}

\begin{figure}[!htp]
    \includegraphics[width=0.95\textwidth]{figs/fig_v13_zbar_change.png}
    \caption{Mean phenotype within cohorts for three time steps following environmental change. One line represents one cohort. For visual clarity, only cohorts age zero through four at the time of the environmental shift for which there were more than 10 observations used to estimate the mean are shown. Circles give the analytical expression defined by Equation 7 in main text.}
    \label{fig:02_supp_zbar_change_validate}
\end{figure}

\begin{figure}[!htp]
    \includegraphics[width=0.95\textwidth]{figs/fig_v14_bvar_change.png}
    \caption{Mean additive genetic variance for each age class ($\gamma^2_{a,k}$) following environmental change, shown for ages zero through nine. Each line shows additive genetic variance in one time step, averaged across 100 trials per parameter combination. Circles give the analytical expression defined by Equation 4 in the main text. Cohorts birthed after the environmental change are excluded from this figure, as disruptions to age structure following environmental change cause a perturbation to population-level means, influencing their means.}
    \label{fig:02_supp_bvar_change_validate}
    % note: ref here is hard coded
\end{figure}

\begin{figure}[!htp]
    \includegraphics[width=0.95\textwidth]{figs/fig_v15_evar_change.png}
    \caption{Mean environmental contribution to phenotypic variance for each age class ($\gamma^2_{e,k}$) following environmental change, shown for ages zero through nine. Each line shows variance in one time step, averaged across 100 trials per parameter combination. Circles give the analytical expression defined by Equation 5 in the main text. Cohorts birthed after the environmental change are excluded from this figure, as disruptions to age structure following environmental change perturb population-level means, influencing cohort means.}
    % note: ref here is hard coded
\end{figure}

\begin{figure}[!htp]
    \includegraphics[width=0.95\textwidth]{figs/fig_v16_zvar_change.png}
    \caption{Mean phenotypic variance for each age class ($\gamma^2_{k}$) following environmental change, shown for ages zero through nine. Each line shows phenotypic variance in one time step, averaged across 100 trials per parameter combination. Circles give the analytical expression defined in the main text and section~\ref{sec:02_pheno_normality}. Cohorts birthed after the environmental change are excluded from this figure, disruptions to age structure following environmental change perturb population-level means, influencing cohort means.}
\end{figure}

\begin{figure}[!htp]
    \includegraphics[width=0.95\textwidth]{figs/fig_v17_rhok_change.png}
    \caption{Mean correlation of breeding values and environmental portions of phenotype for each age class ($\rho_k$) following environmental change, shown for ages zero through nine. Each line shows correlation in one time step, averaged across 100 trials per parameter combination. Circles give the analytical expression defined by Equation 6 in the main text. Cohorts birthed after the environmental change are excluded from this figure, as disruptions to age structure following environmental change perturb population-level means, influencing cohort means.}
    \label{fig:02_supp_rho_change_validate}
\end{figure}

\clearpage

%%%%%%%%%%%%%%%%%%%%%%%%%%%%%%%%%%%%%%%%%%%%%%%%%

\section{Phenotypic component variances in simulations}
\label{sec:02_supp_newton}

Phenotypic variance in the whole population is the age-weighted mean of cohort variances, i.e.,

\begin{align*}
    \gamma^2 = \displaystyle\sum_{k=0}^\infty p_k \gamma^2_k = \displaystyle\sum_{k=0}^\infty \left( \frac{r}{1+r} \right) \left( \frac{\widehat{s}}{\lambda^*} \right)^k \frac{1}{\sqrt{1 + k \gamma^2_0}} \frac{\gamma^2_0}{1 + k\gamma^2_0}.
\end{align*}

For given $\widehat{s}, r,$ and $\gamma^2$ (i.e., specifications for a life history treatment), we solved for $\gamma^2_0$ (phenotypic variance in newborn cohorts) and $\lambda^*$ (equilibrium population growth rate) satisfying the following constraints using Newton's Method:

\begin{align*}
    f_1(\lambda^*, \gamma^2_0) &=  \Bigg( \displaystyle\sum_{k=0}^\infty \left( \frac{r}{1+r} \right) \left( \frac{\widehat{s}}{\lambda^*} \right)^k \frac{1}{\sqrt{1 + k \gamma^2_0}} \frac{\gamma^2_0}{1 + k \gamma^2_0}  \Bigg) - \gamma^2 \\
    f_2(\lambda^*, \gamma^2_0) &= \Bigg( \displaystyle\sum_{k=0}^\infty \left( \frac{r}{1+r} \right) \left( \frac{\widehat{s}}{\lambda^*} \right)^k \frac{1}{\sqrt{1 + k \gamma^2_0}} \Bigg) - 1.
\end{align*}

\noindent $f_1$ is from the equation for variance (see above) and $f_2$ is from the stable age distribution. All infinite sums were estimated by taking the terms for $k = 0$ through $k = 100,000.$

\clearpage

%%%%%%%%%%%%%%%%%%%%%%%%%%%%%%%%%%%%%%%%%%%%%%%%%

\section{Simulations with equal $\widehat{\lambda}$}

To evaluate the effects of the assumed life history trade-off where longer-lived (high-survival, $\widehat{s}$) populations have lower maximum intrinsic growth rates ($\widehat{\lambda}$), we ran a separate batch of simulations that relaxed this assumption. Whereas the simulations analyzed in main text were designed such that the maximum expected lifetime fitness ($\widehat{W}$) was equivalent across life histories (producing a trade-off between per time step survival and per-bout reproductive output), this auxiliary batch of simulations was run with all life histories having equivalent per time step maximum growth rates, set to $\widehat{\lambda} = 1.4;$ this value was arbitrarily chosen. With $\widehat{\lambda}$ held equal across all strategies, the per-reproductive bout fecundity is $r = (\widehat{\lambda} / \widehat{s}) - 1$ and the maximum expected lifetime fitness is $\widehat{W} = (\widehat{\lambda} - \widehat{s}) / (1 - \widehat{s}).$ The intrinsic fitness approaches $\infty$ as $\widehat{s}$ approaches zero for any $\widehat{\lambda} > 1.$ 

We ran a three-way factorial experiment with the same values for $\widehat{s}, \gamma^2,$ and $h^2$ as the main experiment (see Table 1 of main text). Table~\ref{tab:02_aux_treatments} contains all treatments. We ran 500 simulations per parameter combination with all parameters (other than those listed in Table!\ref{tab:02_aux_treatments}) identical to those in our main simulations.
% NOTE: reference to table in main text is hard-coded

\begin{table}[!t]
    \begin{center}
    \begin{tabular}{l r l r  r  r  r}
    \hline
    $\widehat{s}$ & (longevity) & $\gamma^2$ & $r$ & $\widehat{\lambda}$ & $\lambda^*$ & $\widehat{W}$\\
    \hline
    $0.1$ & (low) & $0.10$ & $13$ & $1.4$ & $1.33$ & $1.44$ \\
    $0.1$ & (low) & $0.25$ & $13$ & $1.4$ & $1.25$ & $1.44$ \\
    $0.1$ & (low) & $0.40$ & $13$ & $1.4$ & $1.18$ & $1.44$ \\
    $0.5$ & (medium) & $0.10$ & $1.8$ & $1.4$ & $1.33$ & $1.80$ \\
    $0.5$ & (medium) & $0.25$ & $1.8$ & $1.4$ & $1.25$ & $1.80$ \\
    $0.5$ & (medium) & $0.40$ & $1.8$ & $1.4$ & $1.19$ & $1.80$ \\
    $0.9$ & (high) & $0.10$ & $0.56$ & $1.4$ & $1.33$ & $5.00$ \\
    $0.9$ & (high) & $0.25$ & $0.56$ & $1.4$ & $1.25$ & $5.00$ \\
    $0.9$ & (high) & $0.40$ & $0.56$ & $1.4$ & $1.19$ & $5.00$ \\
    \hline
    \end{tabular}
    \caption[Longevity and phenotypic variance treatments in simulation experiments.]{Maximum survival ($\widehat{s}$) and phenotypic variance ($\gamma^2$) treatments in auxiliary simulation experiment with equal $\widehat{\lambda}$ per treatment. Also included for each treatment are fecundity per mating bout ($r$), equilibrium population growth rate ($\lambda^*$), and maximum expected lifetime fitness ($\widehat{W}$) for each treatment. Heritability treatments are not included in this table as they do not directly influence equilibrium population growth rates.}
    \label{tab:02_aux_treatments}
    \end{center}
\end{table}


\subsection{Results of equal-$\widehat{\lambda}$ simulations}

In contrast to our main results (Figs. 1-2), in the simulations where $\widehat{\lambda}$ was constant across all life histories, populations were largest and extinctions were rarest for the longest-lived treatments (Fig.~\ref{fig:02_aux_popsize}, Fig.~\ref{fig:02_aux_extinctions}). For this auxiliary batch of simulations, for all treatments the shortest-lived life history group had smaller densities than longer-lived counter parts, began going extinct earlier, and were more likely to have gone extinct by the end of the simulation trial. Medium-longevity treatments had both mean population size and extinction rates in between those of long- and short-lived, producing a more monotonic relationship between longevity and these demographic measures than in our main simulations. As with main results, extinctions primarily occurred early in the simulations. Dynamics of rates of adaptation and age structure in these simulations were qualitatively identical to those from the main simulations (Figs. 4-6), albeit with more extinctions occurring that truncated the amount of usable data (results not shown).

\begin{figure}[!htp]
    \includegraphics[width=0.95\textwidth]{amnat_for_submission-copy_for_overleaf_upload/figs/figE1_popsize.png}
    \caption{Mean population size over time for auxiliary simulations. Thick lines are the mean population size of 500 trials per treatment, including extinct populations as size zero. Shaded regions are twice the standard error on each side of the mean (errors are mostly too small to be visible on this plot). Twenty randomly-selected realizations of population size over time per treatment are plotted with thin lines.}
    \label{fig:02_aux_popsize}
\end{figure}

\begin{figure}[!htp]
    \includegraphics[width=0.95\textwidth]{amnat_for_submission-copy_for_overleaf_upload/figs/figE2_extinctions.png}
    \caption{Proportion of surviving populations (out of 500 per treatment) over 100 time steps in auxiliary simulations.}
    \label{fig:02_aux_extinctions}
\end{figure}

\clearpage

%%%%%%%%%%%%%%%%%%%%%%%%%%%%%%%%%%%%%%%%%%%%%%%%%


\section{Time of first observed extinction per life history}

\begin{table}[!htp]
    \centering
    \begin{tabular}{l l r  r  r}
      \hline
      $h^2$ & $\gamma^2$ & low longevity & medium longevity & high longevity \\
      \hline
      $0.25$ & $0.1$ & $6$ & $7$ & $7$ \\
      $0.5$ & $0.1$ & $7$ & $9$ & $8$ \\
      $1$ & $0.1$ & $-$ & $-$ & $9$ \\
      $0.25$ & $0.25$ & $10$ & $14$ & $26$ \\
      $0.5$ & $0.25$ & $-$ & $-$ & $-$ \\
      $1$ & $0.25$ & $-$ & $-$ & $-$ \\
      $0.25$ & $0.4$ & $-$ & $-$ & $57$ \\
      $0.5$ & $0.4$ & $-$ & $-$ & $-$ \\
      $1$ & $0.4$ & $-$ & $-$ & $-$ \\
      \hline
    \end{tabular}
    \caption{First observed time step in simulations where extinctions are observed for each life history. Dash ($-$) indicates zero treatments were observed for the life history.}
\end{table}

\clearpage

%%%%%%%%%%%%%%%%%%%%%%%%%%%%%%%%%%%%%%%%%%%%%%%%%

\section{Age structure in medium- and low-longevity treatments}
\label{sec:02_supp_age_fig}

\begin{figure}[!htp]
    \centering
    \includegraphics[width=0.95\textwidth]{figs/figS20_agedist_medlowlong.png}
    \caption[Age distribution over time for medium and low longevity treatments.]{Age distribution over time for medium and low longevity treatments. Individuals age four and older are combined into one color code for visual clarity.}
    \label{fig:02_supp_agedist}
\end{figure}

\clearpage

%%%%%%%%%%%%%%%%%%%%%%%%%%%%%%%%%%%%%%%%%%%%%%%%%

\section{Phenotypic change scaled by generation time}

We estimated generation time using the formula

\begin{align*}
    T = \displaystyle\sum_{k=0}^\infty k \lambda^{-k} l(k) b(k),
\end{align*}

\noindent where $l(k)$ is the cumulative probability of surviving to age $k,$ $b(k)$ is the expected reproductive output at age $k,$ and $\lambda$ is the equilibrium population growth rate. This formula gives the mean age of parents in a population, averaged over offspring in the population at the stable age distribution (Caswell 2001, p. 129, eq. 5.77). Other formulae may also be used for generation time (Caswell 2001, pp. 128-129); we used this one because it is the relevant timescale for considering rates of phenotypic change due to transmission of parental breeding values to offspring. 

As in Section~\ref{sec:02_supp_age_fig}, we use $l(k) = \widehat{s}^k / \sqrt{1 + k\gamma^2_0},$ $b(k) = r$ (expected reproductive output per individual is $r$ for all ages) and $\lambda = \lambda^*.$ As such, in our model the formula for generation time is

\begin{align}
    T = \displaystyle\sum_{k=0}^\infty k \left( \frac{\widehat{s}}{\lambda^*} \right)^k \sqrt{\frac{1}{1 + k \gamma^2_0}} r.
\end{align}

\noindent Using this formula, our generation times for the simulation experiment (for population-level phenotypic variance is $\gamma^2 = 0.4,$ the only trials for which we measured phenotypic variance in each time step) are $T = 1.04$ for the low-longevity treatment, $T = 1.36$ for the medium-longevity treatment, and $T = 5.68$ for the high-longevity treatment. We used these values to re-scale time to be in units of generations to evaluate phenotypic change for each treatment on the scale of the pace of life of each organism. Figure~\ref{fig:02_supp_gen_time} shows that, because of the longer generation time of the high-longevity populations, on a per-generation basis high-longevity populations have faster rates of phenotypic change than low-longevity populations on a per-time step basis (compare with Fig. 4, main text). Furthermore, Figure~\ref{fig:02_supp_gen_time} suggests that the phase of increase in magnitude for the non-inherited phenotypic components ($\bar{e}$) is one generation.
% NOTE main text ref is hard-coded

\begin{figure}[!h]
    \includegraphics[width=0.95\textwidth]{figs/figS21_generation_time_scaling.png}
    \caption{Mean phenotypic components ($\bar{z}$, $\bar{b}$, $\bar{e}$) for each longevity treatment, with time re-scaled to reflect the generation times for each treatment. Only the first ten generations are shown; the high-longevity treatment has fewer than ten generations because the length of simulations (50 time steps) was shorter than ten generations for this treatment.}
    \label{fig:02_supp_gen_time}
\end{figure}

\clearpage

%%%%%%%%%%%%%%%%%%%%%%%%%%%%%%%%%%%%%%%%%%%%%%%%%

\section{Lifetime fitness under a pace-of-life trade-off}

For notational ease, denote the function $s_z = \widehat{s} \exp{(- z^2 / 2)},$ defining the per-time step survival probability of an individual with phenotype $z.$ As mentioned in Methods, the lifetime (absolute) fitness of an individual with phenotype $z$ is 

\begin{align*}
    W(z) = \frac{r s_z}{1 - s_z}. 
\end{align*}

\noindent Under the assumption that maximum lifetime (absolute) fitness is fixed at $\widehat{W}$ and $r$ is accordingly defined as $r = \widehat{W} \left( 1 - \widehat{s} \right) / \widehat{s},$ absolute fitness of an individual can be rewritten as

\begin{align*}
    W(z) & = \widehat{W} \left( \frac{1 - \widehat{s}}{\widehat{s}} \right) \left( \frac{s_z}{1 - s_z} \right) \\
    & = \widehat{W} \left(1 - \widehat{s} \right) \frac{s_z / \widehat{s}}{1 - s_z} \\
    & = \widehat{W} \frac{\left( 1 - \widehat{s} \right) \exp{(-z^2/2)}}{1 - \widehat{s}\exp{(-z^2/2})}.
\end{align*}

\noindent This is a monotonically decreasing function for $\widehat{s}.$ In the limiting case of $\widehat{s} = 0,$ the lifetime fitness is $W(z) = \widehat{W} \exp{(-z^2 / 2)},$ the same as in Gomulkiewicz and Holt's (1995) original model. 

\begin{figure}[!htp]
    \centering
    \includegraphics[width=0.95\linewidth]{figs/figS22_lifetime_fitness.png}
    \caption{Lifetime fitness ($W(z)$) across maximum survival, $\widehat{s}.$ Highlighted lines are survival values used in longevity treatments for simulations described in main text. $\widehat{W} = 3$ for all curves, matching the value used in simulations. Horizontal gray line is $W = 1,$ i.e., replacement.}
    \label{fig:02_supp_lifetime_fitness}
\end{figure}

Fig.~\ref{fig:02_supp_lifetime_fitness} confirms that the fitness function becomes steeper as survival increases. It also demonstrates by another way the result from Figure 1a (main text) that increasing survival decreases the range of phenotypes that promote persistence, in this case absolute fitness greater than 1. 

% \newpage{}

%%%%%%%%%%%%%%%%%%%%%
% Bibliography
%%%%%%%%%%%%%%%%%%%%%
%
% Any references in this section should also be copied into the main 
% template, under the heading
% \section*{References Cited Only in the Online Enhancements}
% except when they are already cited in the main text.

% \begin{thebibliography}{}

% \end{thebibliography}

\end{document}
