## Supplementary material for "Longevity hinders evolutionary rescue through slower growth but not necessarily slower adaptation": R Code: refs.bib

@article{kea20,
    author = {Chris K. Klausmeier and Matthew M. Osmond and Colin T. Kremer and Elena Lichtman},
    title = {Ecological limits to evolutionary rescue},
    year = {2020},
    journal = PhilosTransRSocLondSerB,
    volume = {375},
    pages = {20190453}
}

@article{pea21,
    author = {James H. Peniston and Michael Barfield and Robert D. Holt and Maria E. Orive},
    title = {Environmental fluctuations dampen the effects of clonal reproduction on evolutionary rescue},
    year = {2021},
    journal = JEvolBiol,
    volume = {34},
    pages = {710 -- 722}
}

@article{gg03,
    author = {Monica A. Geber and Lauren R. Griffen},
    title = {Inheritance and natural selection in functional traits},
    year = {2003},
    journal = IntJPlantSci,
    volume = {164},
    pages = {S21 -- S42}
}

@article{yea19,
    author = {Masato Yamamichi and Nelson G. {Hairston Jr}. and Mark Rees and Stephen P. Ellner},
    title = {Rapid evolution with generation overlap: the double-edged effect of dormancy},
    year = {2019},
    journal = {Theoretical Ecology},
    volume = {12},
    pages = {179 -- 195}
}

@article{dea02,
    author = {Nicola A. Denney and Simon Jennings and John D. Reynolds},
    title = {Life-history correlates of maximum population growth rates in marine fishes},
    year = {2002},
    journal = ProcRSocLondSerB,
    volume = {269},
    pages = {2229 -- 2237}
}

@article{mh08,
    author = {Brett A. Melbourne and Alan H. Hastings},
    title = {Extinction risk depends strongly on factors contributing to stochasticity},
    year = {2008},
    journal = Nature,
    volume = {454},
    pages = {100 -- 103}
}

@article{c-wea24,
    author = {T.J. Clark-Wolf and P. Dee Boersma and Floriane Plard and Ginger E. Rebstock and Briana Abrahms},
    title = {Increasing environmental variability inhibits evolutionary rescue in a long-lived vertebrate},
    year = {2024},
    journal = ProcNatlAcadSciUSA,
    volume = {121},
    number = {34},
    pages = {e2406314121}
}

@article{oea17,
    author = {Maria E. Orive and Michael Barfield and Carlos Fernandez and Robert D. Holt},
    title = {Effects of clonal reproduction on evolutionary lag and evolutionary rescue},
    year = {2017},
    journal = AmNat,
    volume = {190},
    number = {4},
    pages = {469 -- 490}
}

@article{aw84,
    author = {Stevan J. Arnold and Michael J. Wade},
    title = {On the measurement of natural and sexual selection: theory},
    year = {1984},
    journal = Evolution,
    volume = {38},
    number = {4},
    pages = {709 -- 719}
}

@article{cc20,
    author = {Olivier Cotto and Luis-Miguel Chevin},
    title = {Fluctuations in lifetime selection in an autocorrelated environment},
    year = {2020},
    journal = TheorPopulBiol,
    volume = {134},
    pages = {119 -- 128}
}

@article{dea24,
    author = {Jeremy A. Draghi and Joel W. McGlothlin and Holly K. Kindsvater},
    title = {Demographic feedbacks during evolutionary rescue can slow or speed adaptive evolution},
    year = {2024},
	journal = ProcRSocLondSerB,
    number = {291},
    pages = {20231553},
}

@book{c01,
    author = {Hal Caswell},
    title = {Matrix Population Models},
    year = {2001},
    publisher = {Sinauer Associates, Sunderland, Massachusetts, USA},
    edition = {Second Edition}
}

@article{nea23,
	author = {Scott W. Nordstrom and Ruth A. Hufbauer and Laure Olazcuaga and Lily F. Durkee and Brett A. Melbourne},
	date-added = {2023-11-29 13:24:56 -0800},
	date-modified = {2023-11-29 13:26:40 -0800},
	journal = ProcRSocLondSerB,
	pages = {20231228},
	title = {How density dependence, genetic erosion, and the extinction vortex impact evolutionary rescue},
	volume = {290},
	year = {2023}}

@incollection{gs86,
	author = {Michael E. Gilpin and Michael E. Soul\'{e}},
	title = {Minimum viable populations: processes of extinction},
	date-added = {2023-06-15 09:19:19 -0600},
	date-modified = {2023-07-13 14:33:29 -0600},
	editor = {Michael E. Soul\'{e}},
	publisher = {Sinauer Associates, Sunderland, Massachusetts, USA},
	booktitle = {Conservation Biology: The Science of Scarcity and Diversity},
	year = {1986}}

@article{fea99,
	author = {Richard Frankham and Kelly Lees and Margaret E. Montgomery and Philip R. England and Edwin H. Lowe and David A. Briscoe},
	date-added = {2023-06-13 21:46:43 -0600},
	date-modified = {2023-06-13 21:49:30 -0600},
	journal = {Animal Conservation},
	pages = {255--260},
	title = {Do population size bottlenecks reduce evolutionary potential?},
	volume = {2},
	year = {1999}}

@article{w00,
	author = {Michael C. Whitlock},
	date-added = {2023-06-15 18:06:28 -0600},
	date-modified = {2023-06-15 18:07:01 -0600},
	journal = {Evolution},
	number = {6},
	pages = {1855--1861},
	title = {Fixation of new alleles and the extinction of small populations: drift load, beneficial alleles, and sexual selection},
	volume = {54},
	year = {2000}}

@article{wea16,
	author = {Jacquelyn L. A. Wood and Matthew C. Yates and Dylan J. Fraser},
	date-added = {2023-06-15 17:17:02 -0600},
	date-modified = {2023-06-23 21:38:39 -0600},
	journal = {Evolutionary Applications},
	number = {5},
	pages = {640--647},
	title = {Are heritability and selection related to population size in nature? {M}eta-analysis and conservation implications},
	volume = {9},
	year = {2016}}

@article{h92,
	author = {David Houle},
	date-added = {2023-06-14 09:42:05 -0600},
	date-modified = {2023-06-14 09:43:09 -0600},
	journal = Genetics,
	pages = {195--204},
	title = {Comparing evolvability and variability of quantitative traits},
	volume = {130},
	year = {1992}}

@article{bh16,
	author = {Michael Barfield and Robert D. Holt},
	date-added = {2023-06-15 09:18:28 -0600},
	date-modified = {2023-06-15 09:19:05 -0600},
	journal = EvolEcolRes,
	pages = {771--786},
	title = {Evolutionary rescue in novel environments: towards improving predictability},
	volume = {17},
	year = {2016}}

@article{km14,
	author = {Michael Kopp and Sebastian Matuszewski},
	date-added = {2023-06-13 21:36:27 -0600},
	date-modified = {2023-06-13 21:38:16 -0600},
	journal = {Evolutionary Applications},
	pages = {169--191},
	title = {Rapid evolution of quantitative traits: theory and theoretical perspectives},
	volume = {7},
	year = {2014}}

@article{odm13,
	author = {Matthew M. Osmond and Claire {de Mazancourt}},
	date-added = {2023-07-13 18:08:23 -0600},
	date-modified = {2023-07-13 18:09:06 -0600},
	journal = ProcRSocLondSerB,
	pages = {20120085},
	title = {How competition affects evolutionary rescue},
	volume = {368},
	year = {2013}}

@article{to80,
	author = {S. D. Tuljapurkar and Steven H. Orzack},
	date-added = {2023-07-11 19:31:37 -0600},
	date-modified = {2023-07-11 19:31:37 -0600},
	journal = TheorPopBio,
	number = {3},
	pages = {314--342},
	title = {Population dynamics in variable environments I. Long-run growth rates and extinction},
	volume = {18},
	year = {1980}}

@article{ns75,
	author = {Toshio Nakagawa and Shunji Osaki},
	date-added = {2023-07-11 18:40:57 -0600},
	date-modified = {2023-07-11 18:40:57 -0600},
	journal = {IEEE Transactions on Reliability},
	number = {5},
	pages = {300-301},
	title = {The discrete {W}eibull distribution},
	volume = {R-24},
	year = {1975}}

@article{price70,
	author = {George R. Price},
	date-added = {2023-07-11 18:38:39 -0600},
	date-modified = {2023-07-11 18:38:39 -0600},
	journal = Nature,
	pages = {520--521},
	title = {Selection and covariance},
	volume = {227},
	year = {1970}}

@article{kea11,
	author = {Bruce E. Kendall and Gordon Fox and Masami Fujiwara and Theresa Nogeire},
	date-added = {2023-07-11 18:31:24 -0600},
	date-modified = {2023-07-11 18:31:24 -0600},
	journal = Ecology,
	number = {10},
	pages = {1985--1993},
	title = {Demographic heterogeneity, cohort selection and population growth},
	volume = {92},
	year = {2011}}

@article{cea21,
	author = {Aldo Compagnoni and Sam Levin and Dylan Z. Childs and Stan Harpole and Maria Paniw and Gesa R\"{o}mer and Jean H. Burns and Judy Che-Castaldo and Nadja R\"{u}ger and Georges Kunstler and Joanne M. Bennett and C. Ruth Archer and Owen R. Jones and Robert Salguero-G\'{o}mez and Tiffany M. Knight},
	date-added = {2023-07-11 18:31:09 -0600},
	date-modified = {2023-07-11 18:31:09 -0600},
	journal = {Nature Communications},
	number = {1},
	title = {Herbaceous perennial plants with short generation time have stronger responses to climate anomalies than those with longer generation time},
	volume = {12},
    pages = {1824},
	year = {2021}}

@article{dea10,
	author = {Harmony J. Dalgleish and David N. Koons and Peter B. Adler},
	date-added = {2023-07-11 18:31:04 -0600},
	date-modified = {2023-07-11 18:31:04 -0600},
	journal = JEcol,
	number = {1},
	pages = {209--217},
	title = {Can life-history traits predict the response of forb populations to changes in climate variability?},
	volume = {98},
	year = {2010}}

@article{vea79,
	author = {Jamers W. Vaupel and Kenneth G. Monton and Eric Stallard},
	date-added = {2023-07-11 18:30:57 -0600},
	date-modified = {2023-07-11 18:30:57 -0600},
	journal = {Demography},
	number = {3},
	pages = {439--454},
	title = {The impact of heterogeneity in individual frailty on the dynamics of mortality},
	volume = {16},
	year = {1979}}

@book{fm96,
	author = {Douglas S. Falconer and Trudy F.C. Mackay},
	date-added = {2023-07-11 18:30:51 -0600},
	date-modified = {2023-07-13 14:33:17 -0600},
	edition = {Fourth},
	publisher = {Longman, London, UK},
	title = {Introduction to Quantitative Genetics},
	year = {1996}}

@article{c03,
	author = {Marcel Cardillo},
	date-added = {2023-07-11 18:30:35 -0600},
	date-modified = {2023-07-11 18:30:35 -0600},
	journal = AnimConserv,
	number = {1},
	pages = {63--69},
	title = {Biological determinants of extinction: why are smaller species less vulnerable?},
	volume = {6},
	year = {2003}}

@article{s-gea16,
	author = {Roberto Salguero-G\'{o}mez and Owen R. Jones and Eelke Jongejans and Simon P. Blomberg and David J. Hodgson and Cyril Mbeau-Ache and Pieter A. Zuidema and Hans de Kroon and Yvonne M. Buckley},
	date-added = {2023-07-11 18:30:30 -0600},
	date-modified = {2023-07-11 18:30:30 -0600},
	journal = ProcNatlAcadSciUSA,
	number = {1},
	pages = {230--235},
	title = {Fast-slow continuum and reproductive strategies structure plant life-history variation worldwide},
	volume = {113},
	year = {2016}}

@article{o04,
	author = {Mandan K. Oli},
	date-added = {2023-07-11 18:30:23 -0600},
	date-modified = {2023-07-11 18:30:23 -0600},
	journal = BasicApplEcol,
	number = {5},
	pages = {449--463},
	title = {The fast-slow continuum and mammalian life-history patterns: an empirical evaluation},
	volume = {5},
	year = {2004}}

@article{tl79,
	author = {Alan R. Templeton and Donald A. Levin},
	date-added = {2023-07-11 18:30:13 -0600},
	date-modified = {2023-07-11 18:30:13 -0600},
	journal = AmNat,
	number = {2},
	pages = {232--249},
	title = {Evolutionary consequences of seed banks},
	volume = {114},
	year = {1979}}

@article{jea14,
	author = {Owen R. Jones and Alexander Scheurlein and Roberto Salguero-G\'{o}mez and Carlo Giovanni Camarda and Ralf Schaible and Brenda B. Casper and Johan P. Dahlgren and Johan Ehrl\'{e}n and Mar\'{i}a B. Garcia and Eric S. Menges and Pedro F. Quintana-Ascencio and Hal Caswell and Annette Baudisch and James W. Vaupel},
	date-added = {2023-07-11 18:30:03 -0600},
	date-modified = {2023-07-11 18:30:03 -0600},
	journal = Nature,
	pages = {169--173},
	title = {Diversity of ageing across the tree of life},
	volume = {505},
	year = {2014}}

@article{ct08,
	author = {Tim Coulson and Shripad Tuljapurkar},
	date-added = {2023-07-11 18:29:56 -0600},
	date-modified = AmNat,
	journal = AmNat,
	number = {5},
	pages = {599--612},
	title = {The dynamics of a quantitative trait in an age-structured population living in a variable environment},
	volume = {172},
	year = {2008}}

@article{sl18,
	author = {William R. Shoemaker and Jay T. Lennon},
	date-added = {2023-07-11 18:29:50 -0600},
	date-modified = {2023-07-11 18:29:50 -0600},
	journal = {Evolutionary Applications},
	number = {1},
	pages = {60--75},
	title = {Evolution with a seed bank: the population genetic consequences of microbial dormancy},
	volume = {11},
	year = {2018}}

@article{cea93,
	author = {F. Stuart Chapin and Kellar Autumn and Francisco Pugnaire},
	date-added = {2023-07-11 18:29:43 -0600},
	date-modified = {2023-07-13 16:57:24 -0600},
	journal = AmNat,
	pages = {78--92},
	title = {Evolution of suites of traits in response to environmental stress},
	volume = {142},
	year = {1993}}

@article{gb70,
	author = {Madhav Gadgil and William H. Bossert},
	date-added = {2023-07-11 18:28:29 -0600},
	date-modified = {2023-07-11 18:28:29 -0600},
	journal = AmNat,
	number = {935},
	pages = {1--24},
	title = {Life historical consequences of natural selection},
	volume = {104},
	year = {1970}}

@article{pianka70,
	author = {Eric R. Pianka},
	date-added = {2023-07-11 18:23:31 -0600},
	date-modified = {2023-07-11 18:23:31 -0600},
	journal = AmNat,
	number = {940},
	pages = {592--597},
	title = {On r- and {K}-selection},
	volume = {104},
	year = {1970}}

@article{pea00,
	author = {Andy Purvis and John L. Gittleman and Guy Cowlishaw and Georgiana M. Mace},
	date-added = {2023-07-11 18:22:38 -0600},
	date-modified = {2023-07-11 18:22:38 -0600},
	journal = ProcRSocLondSerB,
	number = {1456},
	pages = {1947--1952},
	title = {Predicting extinction risk in declining species},
	volume = {267},
	year = {2000}}

@article{lea21,
	author = {Kelsey P. Lyberger and Matthew M. Osmond and Sebastian J. Schreiber},
	date-added = {2023-06-14 09:53:07 -0600},
	date-modified = {2023-06-14 09:56:02 -0600},
	journal = AmNat,
	number = {1},
	pages = {44--52},
	title = {Is evolution in response to extreme events good for population persistence?},
	volume = {198},
	year = {2021}}

@article{lea22,
	author = {Alessandra Loria and Melanie Cristecu and Andrew Gonzalez},
	date-added = {2023-07-11 18:19:57 -0600},
	date-modified = {2023-07-11 18:22:00 -0600},
	journal = JEvolBiol,
	number = {2},
	pages = {265--277},
	title = {Genotype diversity promotes the persistence of \textit{Daphnia} populations exposed to severe copper stress},
	volume = {35},
	year = {2022}}

@article{b12,
	author = {Graham Bell},
	date-added = {2023-07-11 18:18:58 -0600},
	date-modified = {2023-07-11 18:19:31 -0600},
	journal = BiolLett,
	pages = {20120823},
	title = {Evolutionary rescue of a green alga kept in the dark},
	volume = {9},
	year = {2012}}

@article{mea99,
	author = {J. M. Milner and S. D. Albion and A. W. Illius and J. M. Pemberton and T. H. Clutton-Brock},
	date-added = {2023-06-25 18:26:28 -0600},
	date-modified = {2023-06-25 18:26:28 -0600},
	journal = JAnimEcol,
	number = {3},
	pages = {472--488},
	title = {Repeated selection of morphometric traits in the {S}oay sheep on {S}t. {K}ilda},
	volume = {68},
	year = {1999}}

@article{vwea13,
	author = {E. {Vander Wal} and D. Garant and M. Festa-Bianchet and F. Pelletier},
	date-added = {2023-06-25 18:13:54 -0600},
	date-modified = {2023-06-25 18:13:54 -0600},
	journal = PhilosTransRSocLondSerB,
	pages = {20120090},
	title = {Evolutionary rescue in vertebrates: evidence, applications and uncertainty},
	volume = {368},
	year = {2013}}

@article{jea22,
	author = {John Jackson and Christie {Le Coeur} and Owen Jones},
	date-added = {2023-06-25 18:09:04 -0600},
	date-modified = {2023-06-25 18:09:04 -0600},
	journal = {eLife},
	pages = {e74161},
	title = {Life history predicts global population responses to the weather in terrestrial mammals},
	volume = {11},
	year = {2022}}

@article{hds88,
	author = {Nelson G. Hairston and Bart T. {{d}e Stasio}},
	date-added = {2023-06-24 11:44:29 -0600},
	date-modified = {2023-06-24 11:47:17 -0600},
	journal = Nature,
	pages = {239--242},
	title = {Rate of evolution slowed by a dormant propagule pool},
	volume = {366},
	year = {1988}}

@article{mea08,
	author = {William F. Morris and Catherine A. Pfister and Shripad Tuljapurkar and Chirrakal V. Haridas and Carol L. Boggs and Mark S. Boyce and Emilio M. Bruna and Don R. Church and Tim Coulson and Daniel F. Doak and Stacey Forsyth and Jean-Michel Gaillard and Carol C. Horvitz and Susan Kalisz and Bruce E. Kendall and Tiffany M. Knight and Charlotte T. Lee and Eric S. Menges},
	date-added = {2023-06-24 11:37:49 -0600},
	date-modified = {2023-06-24 11:40:40 -0600},
	journal = Ecology,
	number = {1},
	pages = {19--25},
	title = {Longevity can buffer plant and animal populations against changing climate variability},
	volume = {89},
	year = {2008}}

@article{kea10,
	author = {Anna Kuparinen and Outi Savolainen and Frank M. Schurr},
	date-added = {2023-06-23 19:18:34 -0600},
	date-modified = {2023-06-23 19:19:34 -0600},
	journal = ForEcolManage,
	number = {5},
	pages = {1003--1008},
	title = {Increased mortality can promote evolutionary adaptation of forest trees to climate change},
	volume = {259},
	year = {2010}}

@article{ot89,
	author = {Steven H. Orzack and Shripad Tuljapurkar},
	date-added = {2023-06-23 09:25:10 -0600},
	date-modified = {2023-06-23 09:26:16 -0600},
	journal = AmNat,
	number = {6},
	pages = {901--923},
	title = {Population dynamics in a variable environment. {VII}. {T}he evolution of iteroparity},
	volume = {133},
	year = {1989}}

@article{sea22,
	author = {Max Schmid and Maria Paniw and Maarten Postuma and Arpat Ozgul and Frederic Guillaume},
	date-added = {2023-06-22 23:57:05 -0600},
	date-modified = {2023-06-22 23:58:07 -0600},
	journal = AmNat,
	number = {1},
	pages = {E16--E35},
	title = {A trade-off between robustness to environment fluctuations and speed of evolution},
	volume = {200},
	year = {2022}}

@article{oea10,
	author = {Arpat Ozgul and Dylan Z. Childs and Madan K. Oli and Kenneth B. Armitage and Daniel T. Blumstein and Lucretia E. Olson and Shripad Tuljapurkar and Tim Coulson},
	date-added = {2023-06-22 23:48:25 -0600},
	date-modified = {2023-07-25 17:08:42 -0600},
	journal = Nature,
	pages = {482--485},
	title = {Coupled dynamics of body mass and population growth in response to environmental change},
	volume = {466},
	year = {2010}}

@article{pea88,
	author = {Stuart L. Pimm and H. Lee Jones and Jared Diamond},
	date-added = {2023-06-22 23:42:58 -0600},
	date-modified = {2023-06-22 23:48:15 -0600},
	journal = AmNat,
	number = {6},
	pages = {757--785},
	title = {On the risk of extinction},
	volume = {132},
	year = {1988}}

@article{l09,
	author = {R. Lande},
	date-added = {2023-06-22 16:56:21 -0600},
	date-modified = {2023-06-22 16:58:57 -0600},
	journal = JEvolBiol,
	number = {7},
	pages = {1435--1446},
	title = {Adaptation to an extraordinary environment by evolution of phenotypic plasticity and genetic assimilation},
	volume = {22},
	year = {2009}}

@article{gea05,
	author = {Jean-Michel Gaillard and Nigel G. Yoccoz and Jean-Dominique Lebreton and Christophe Bonenfant and S\'{e}bastien Devillard and Anne Loison and Dominique Pontier and Dominique Allaine},
	date-added = {2023-06-22 16:17:12 -0600},
	date-modified = {2023-06-22 16:18:02 -0600},
	journal = AmNat,
	number = {1},
	pages = {119--123},
	title = {Generation time: a reliable metric to measure life-history variation among mammalian populations},
	volume = {166},
	year = {2005}}

@article{cea17,
	author = {Olivier Cotto and Johannes Wessely and Damien Georges and G\"{u}nther Klonner and Max Schmid and Stefan Dullinger and Willfried Thuiller and Fr\'{e}d\'{e}ric Guillaume},
	date-added = {2023-06-22 16:02:59 -0600},
	date-modified = {2023-06-22 16:04:21 -0600},
	journal = {Nature Communications},
	number = {1},
	title = {A dynamic eco-evolutionary model predicts slow response of alpine plants to climate warming},
	volume = {8},
    pages = {15399},
	year = {2017}}

@article{c54,
	author = {Lamont C. Cole},
	date-added = {2023-06-21 16:51:13 -0600},
	date-modified = {2023-06-21 16:51:48 -0600},
	journal = QRevBiol,
	number = {2},
	pages = {103--137},
	title = {The population consequences of life history phenomena},
	volume = {29},
	year = {1954}}

@article{dea14,
	author = {Rodolfo Dirzo and Hillary S. Young and Mauro Galetti and Gerardo Ceballos and Nick J.B. Isaac and Ben Collen},
	date-added = {2023-06-21 16:46:18 -0600},
	date-modified = {2023-06-21 16:49:01 -0600},
	journal = {Science},
	number = {6195},
	pages = {401--406},
	title = {Defaunation in the Anthropocene},
	volume = {345},
	year = {2014}}

@article{ls96,
	author = {Russell Lande and Susan Shannon},
	date-added = {2023-06-15 18:01:18 -0600},
	date-modified = {2023-06-15 18:02:51 -0600},
	journal = Evolution,
	number = {1},
	pages = {434--437},
	title = {The role of genetic variation in adaptation and population persistence in a changing environment},
	volume = {50},
	year = {1996}}

@misc{RCore,
	author = {{R Core Team}},
	date-added = {2023-06-15 17:19:55 -0600},
	date-modified = {2023-07-13 15:07:04 -0600},
	howpublished = {R Foundation for Statistical Computing, Vienna, Austria},
	title = {R: A language environment for statistical computing},
	url = {http://www.R-project.org/},
	year = {2020},
	Bdsk-Url-1 = {http://www.R-project.org/}}

@book{lw98,
	author = {Michael Lynch and Bruce Walsh},
	date-added = {2023-06-15 17:07:49 -0600},
	date-modified = {2023-07-13 14:33:23 -0600},
	publisher = {Sinauer Associates, Sunderland, Massachusetts, USA},
	title = {Genetics and the Analysis of Quantitative Traits},
	year = {1998}}

@article{bl95,
	author = {Reinhard B\"{u}rger and Michael Lynch},
	date-added = {2023-06-14 09:49:34 -0600},
	date-modified = {2023-06-25 18:05:20 -0600},
	journal = Evolution,
	number = {1},
	pages = {151--163},
	title = {Evolution and extinction in a changing environment: a quantitative-genetic analysis},
	volume = {49},
	year = {1995}}

@article{aea16,
	author = {Jaime Ashander and Luis-Miguel Chevin and Marissa L. Baskett},
	date-added = {2023-06-14 09:46:39 -0600},
	date-modified = {2023-06-25 18:06:11 -0600},
	journal = ProcRSocLondSerB,
	pages = {20161690},
	title = {Predicting evolutionary rescue via evolving plasticity in stochastic environments},
	volume = {283},
	year = {2016}}

@article{l76,
	author = {Russell Lande},
	date-added = {2023-06-13 21:44:56 -0600},
	date-modified = {2023-06-15 17:26:17 -0600},
	journal = Evolution,
	pages = {314--334},
	title = {Natural selection and random genetic drift in phenotypic evolution},
	volume = {30},
	year = {1976}}

@article{cl10,
	author = {Luis-Miguel Chevin and Russell Lande},
	date-added = {2023-06-13 21:41:35 -0600},
	date-modified = {2023-06-21 13:47:59 -0600},
	journal = Evolution,
	number = {4},
	pages = {1143--1150},
	title = {When do adaptive plasticity and genetic evolution prevent extinction of a density-regulated population?},
	volume = {64},
	year = {2010}}

@article{hea15,
	author = {Ruth A. Hufbauer and Marianna Sz\"{u}cs and Emily Kayson and Courtney Youngberg and Michael J. Koontz and Christopher Richards and Ty Tuff and Brett A. Melbourne},
	date-added = {2023-06-13 21:20:23 -0600},
	date-modified = {2023-07-25 17:09:23 -0600},
	journal = ProcNatlAcadSciUSA,
	number = {33},
	pages = {10557--19562},
	title = {Three types of rescue can avert extinction in a changing environment},
	volume = {112},
	year = {2015}}

@article{aea11,
	author = {Deepa Agashe and Jay J. Falk and Daniel I. Bolnick},
	date-added = {2023-06-13 21:18:04 -0600},
	date-modified = {2023-06-13 21:19:10 -0600},
	journal = Evolution,
	number = {9},
	pages = {2481--2491},
	title = {Effects of founding genetic variation on adaptation to a novel resource},
	volume = {65},
	year = {2011}}

@article{bg09,
	author = {Graham Bell and Andrew Gonzalez},
	date-added = {2023-06-13 20:42:59 -0600},
	date-modified = {2023-06-13 20:53:58 -0600},
	journal = EcolLett,
	number = {9},
	pages = {942--948},
	title = {Evolutionary rescue can prevent extinction following environmental change},
	volume = {12},
	year = {2009}}

@article{cea14,
	author = {Stephanie M. Carlson and Curry J. Cunningham and Peter A.H. Westley},
	date-added = {2023-06-13 20:41:38 -0600},
	date-modified = {2023-06-13 20:45:16 -0600},
	journal = TrendsEcolEvol,
	number = {9},
	pages = {521--530},
	title = {Evolutionary rescue in a changing world},
	volume = {29},
	year = {2014}}

@article{gh95,
	author = {Richard Gomulkiewicz and Robert D. Holt},
	date-added = {2023-06-13 18:52:10 -0600},
	date-modified = {2023-06-13 19:26:31 -0600},
	journal = Evolution,
	number = {1},
	pages = {201--207},
	read = {1},
	title = {When does evolution by natural selection prevent extinction?},
	volume = {49},
	year = {1995}}
