## Supplementary figures and images for "Longevity hinders evolutionary rescue through slower growth but not necessarily slower adaptation"

### fig1_lambda_n.png

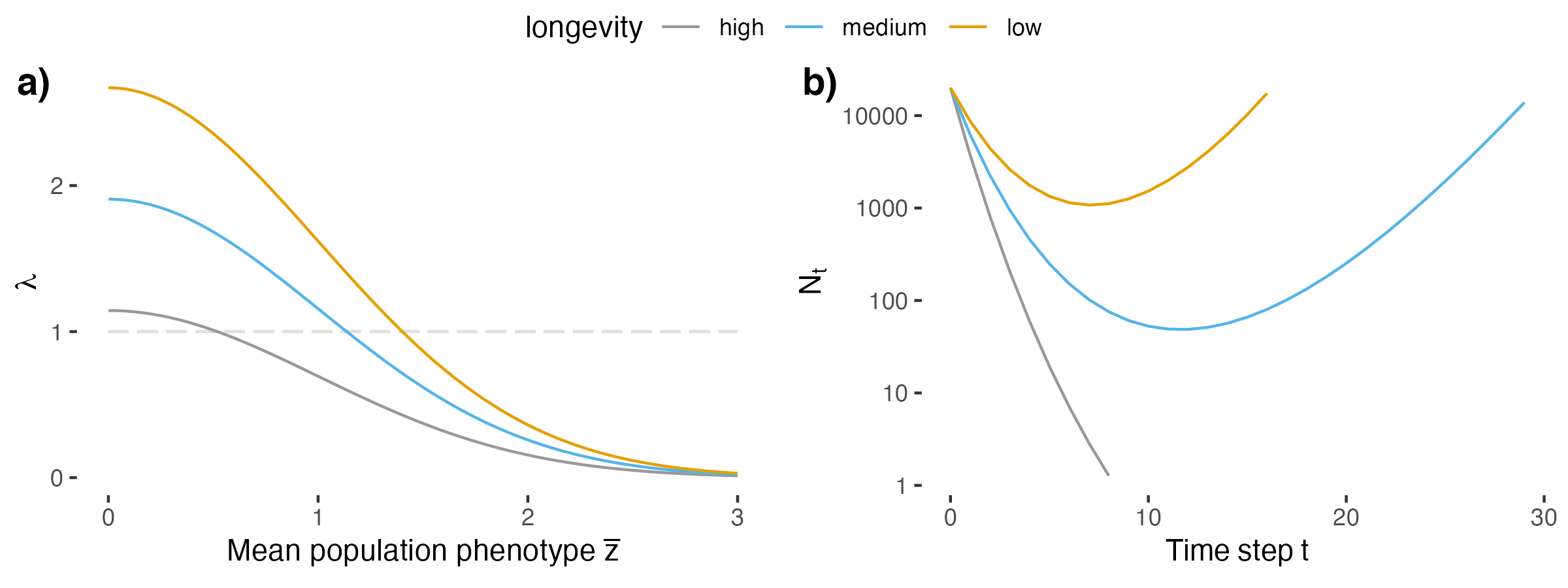

### fig2_popsize.png

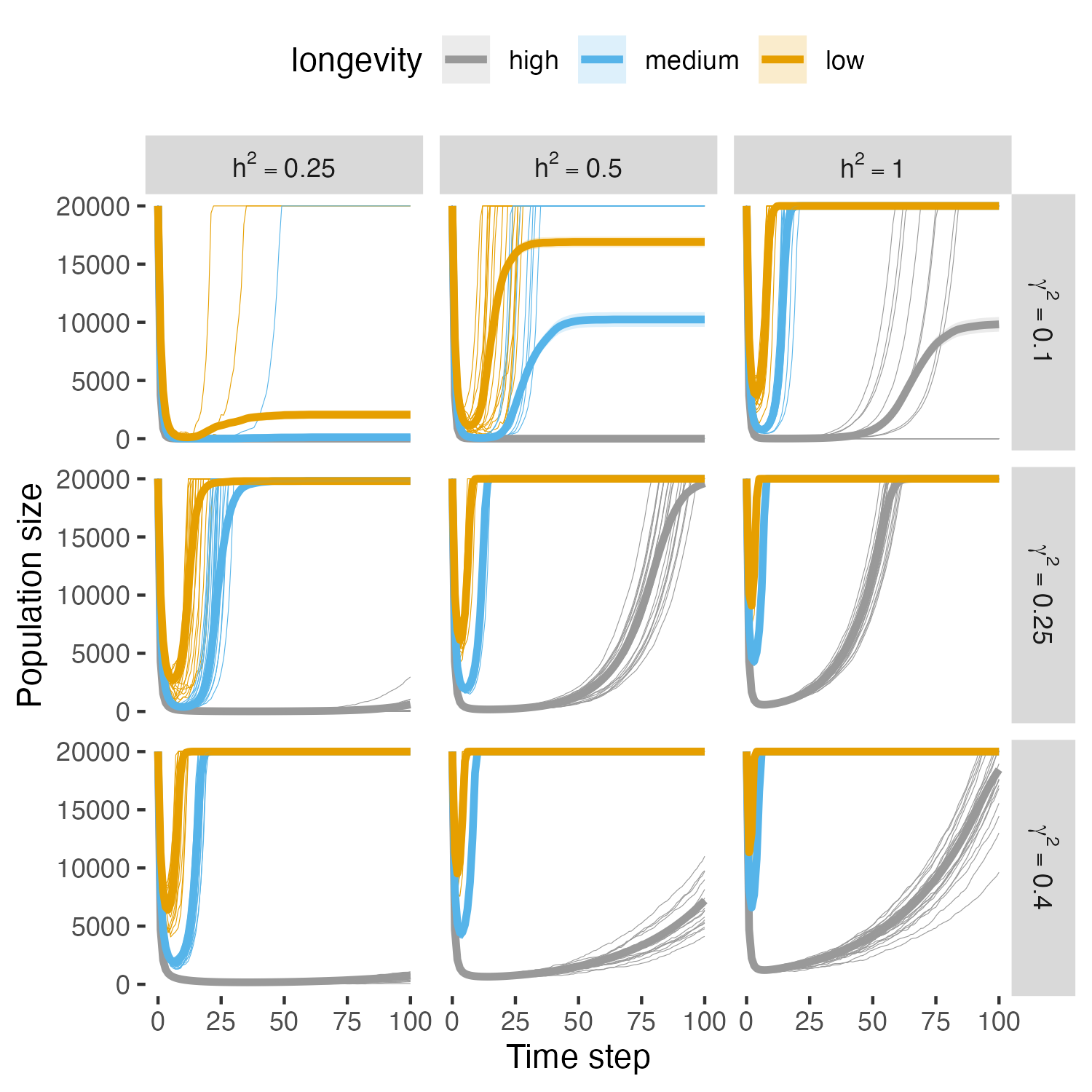

### fig3_extinctions.png

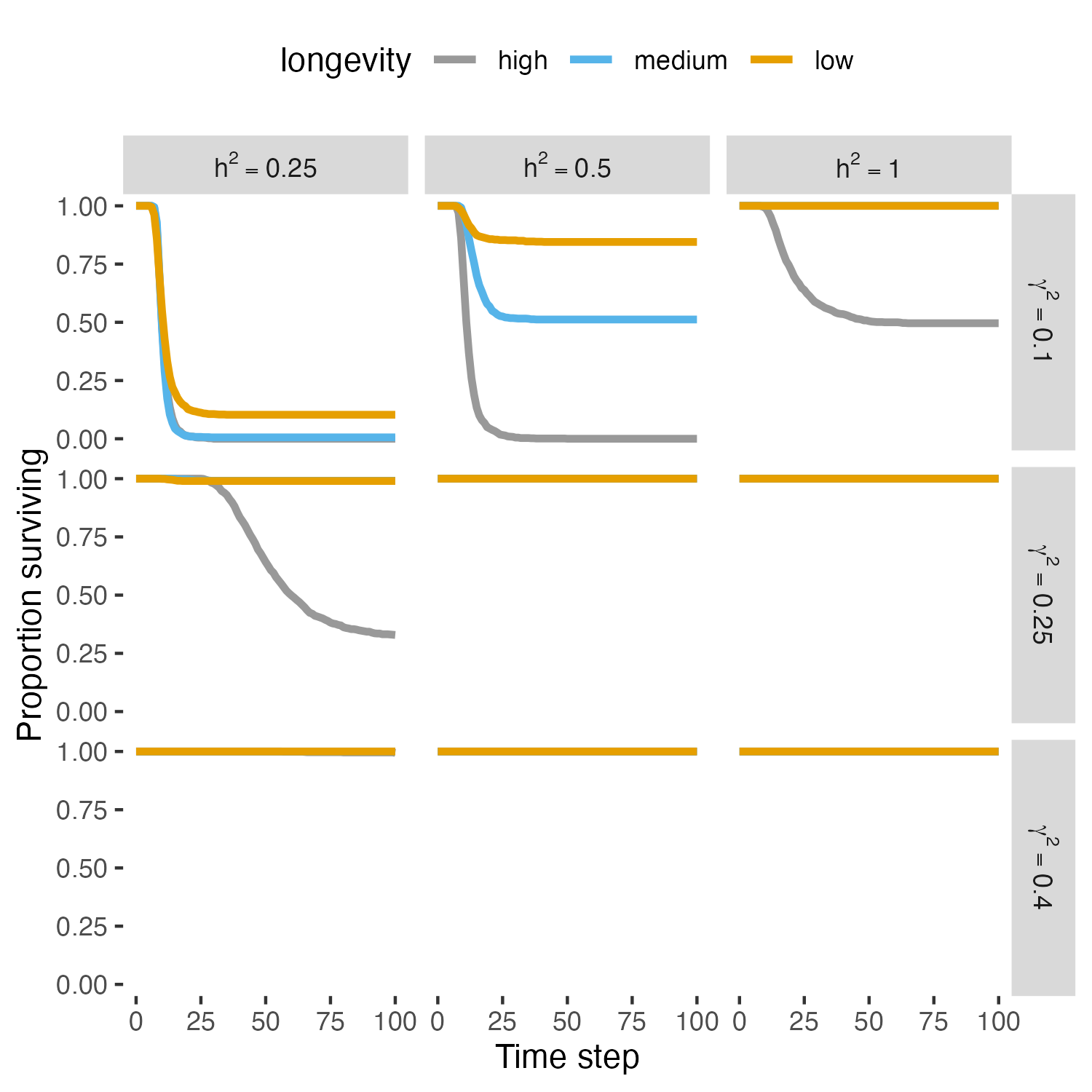

### fig4_pheno_comp_mean.png

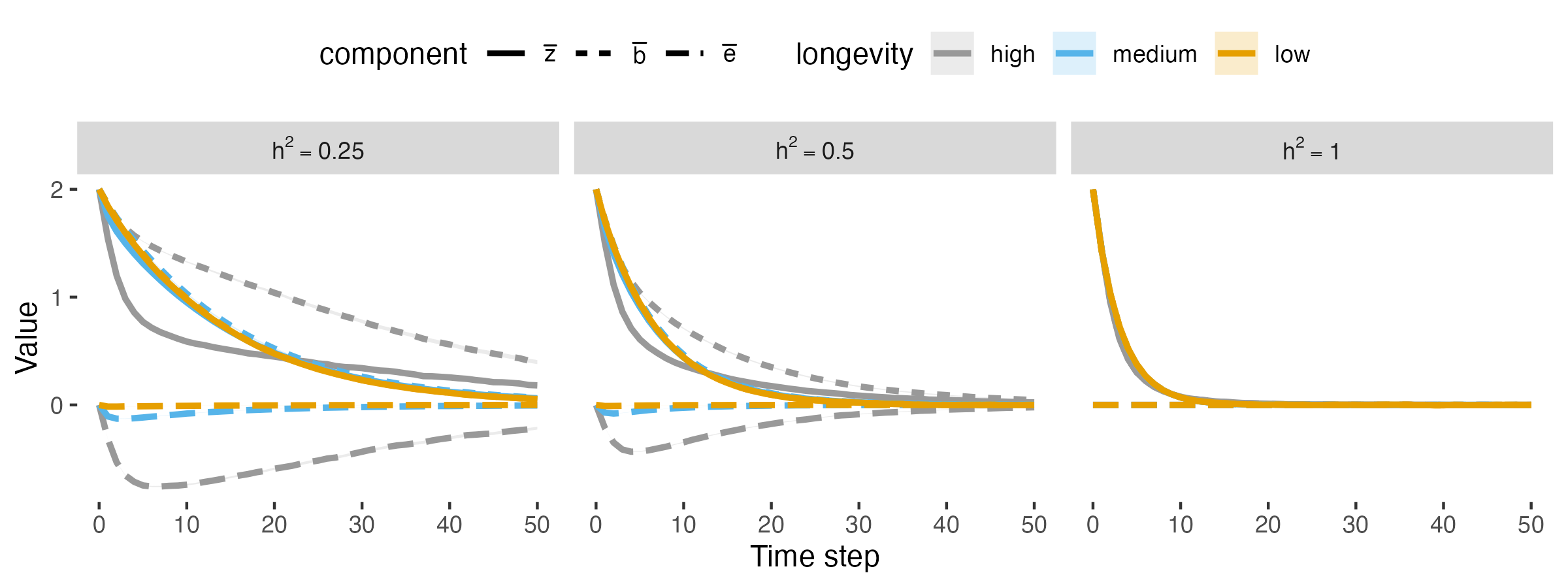

### fig5_agedist_highlong.png

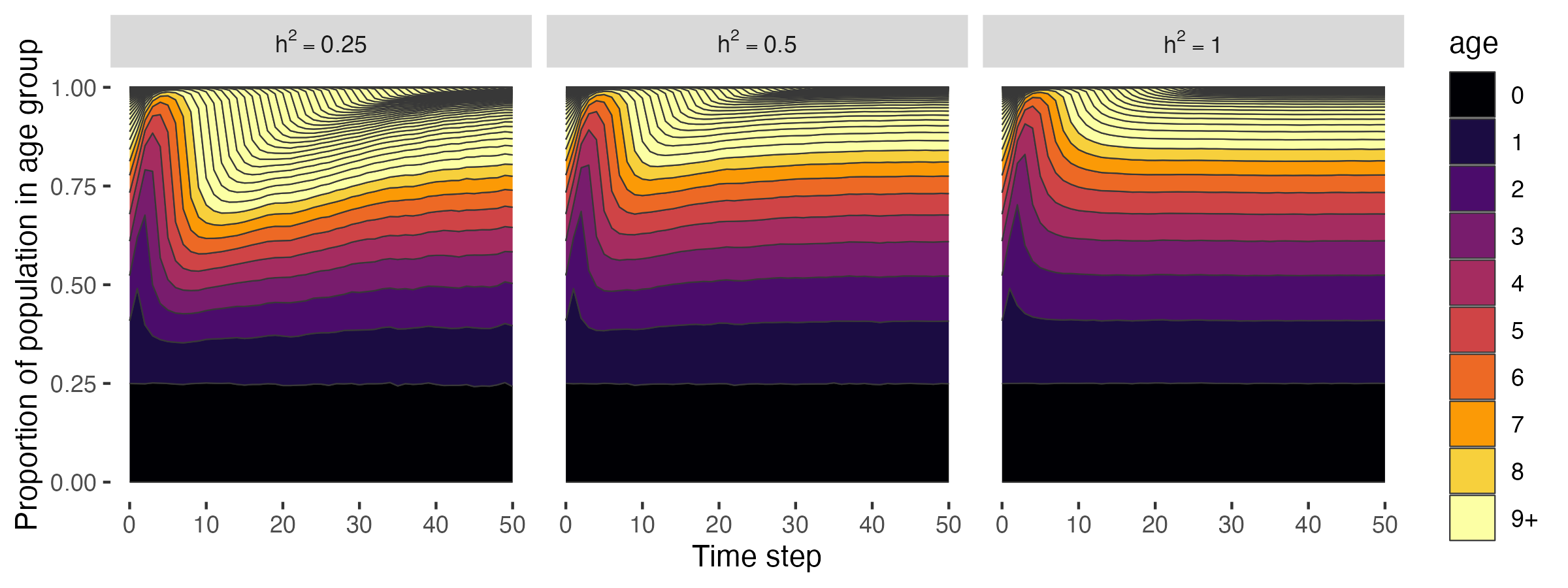

### fig6_pheno_comp_vars.png

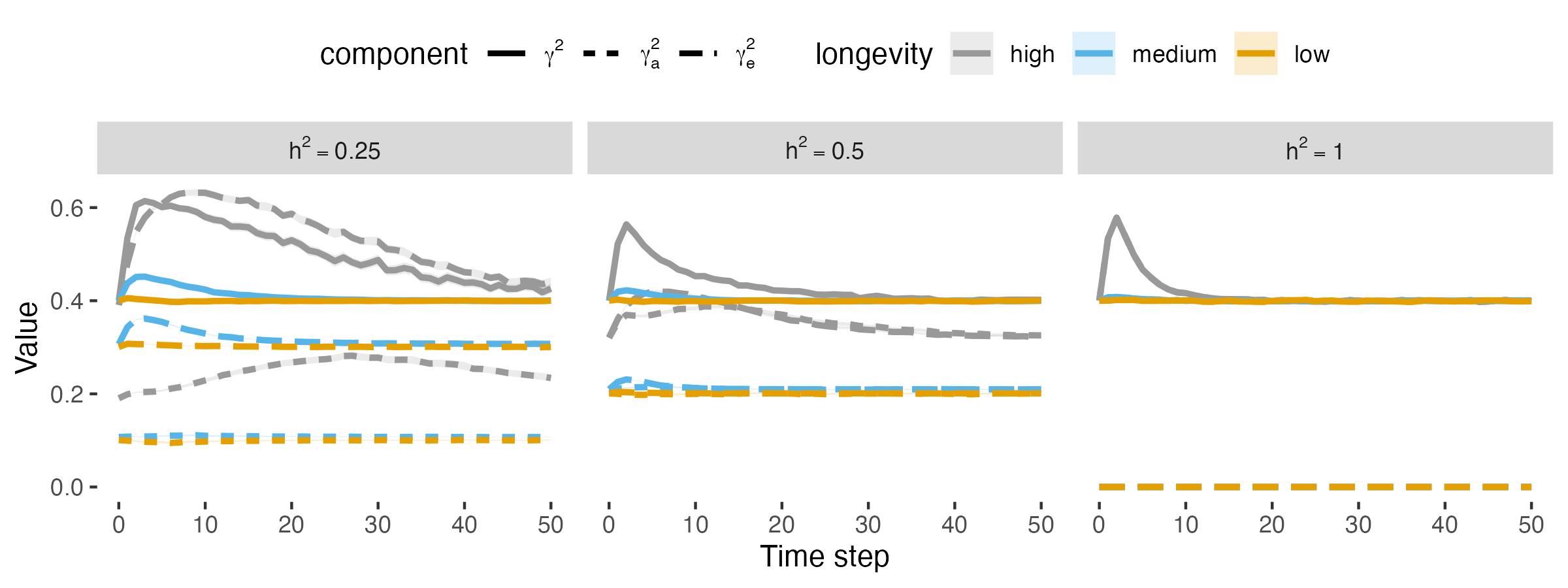

### fig_v2_age_full.png

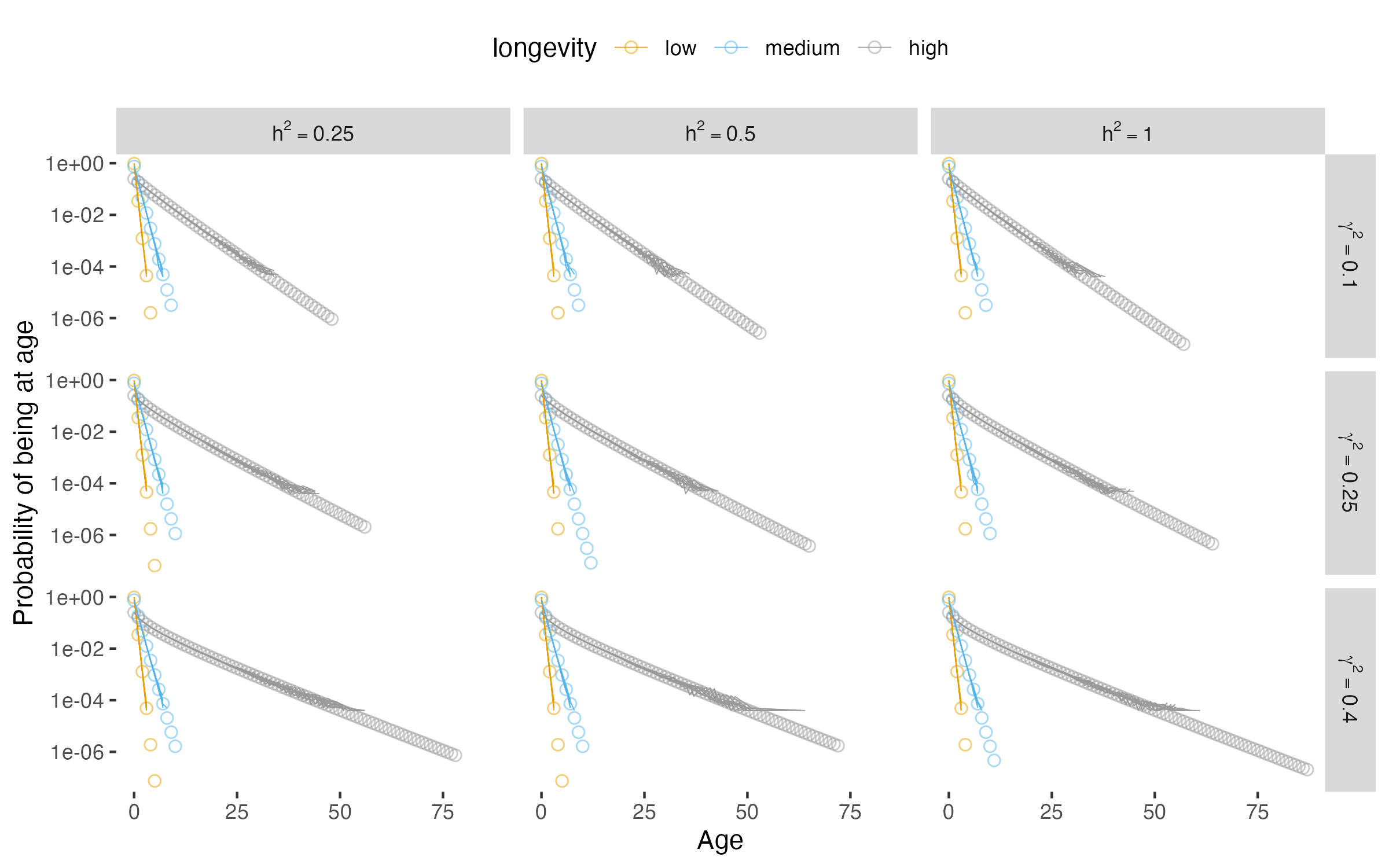

### fig_v3_age_0-9.png

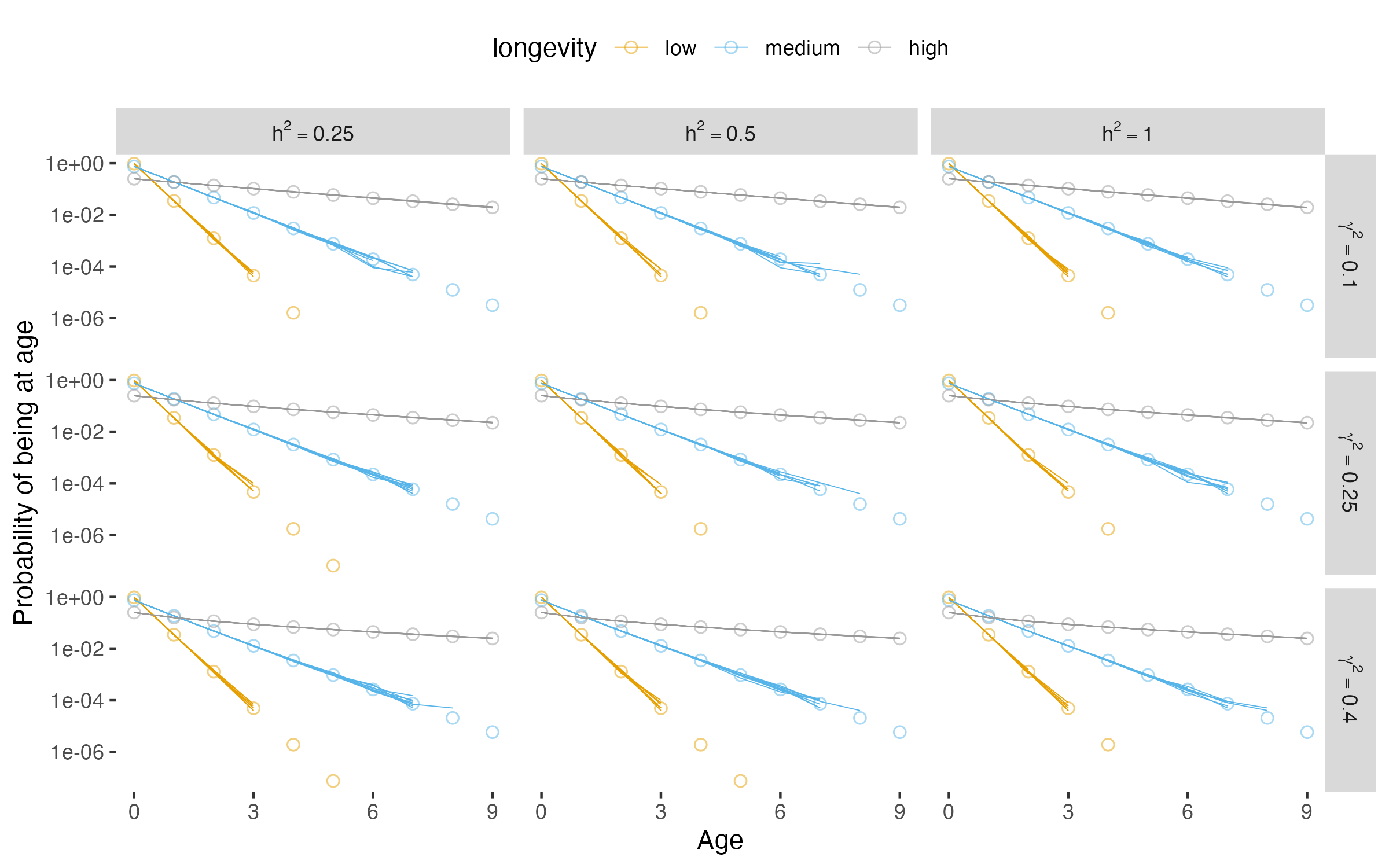

### fig_v4_bbar_steady.png

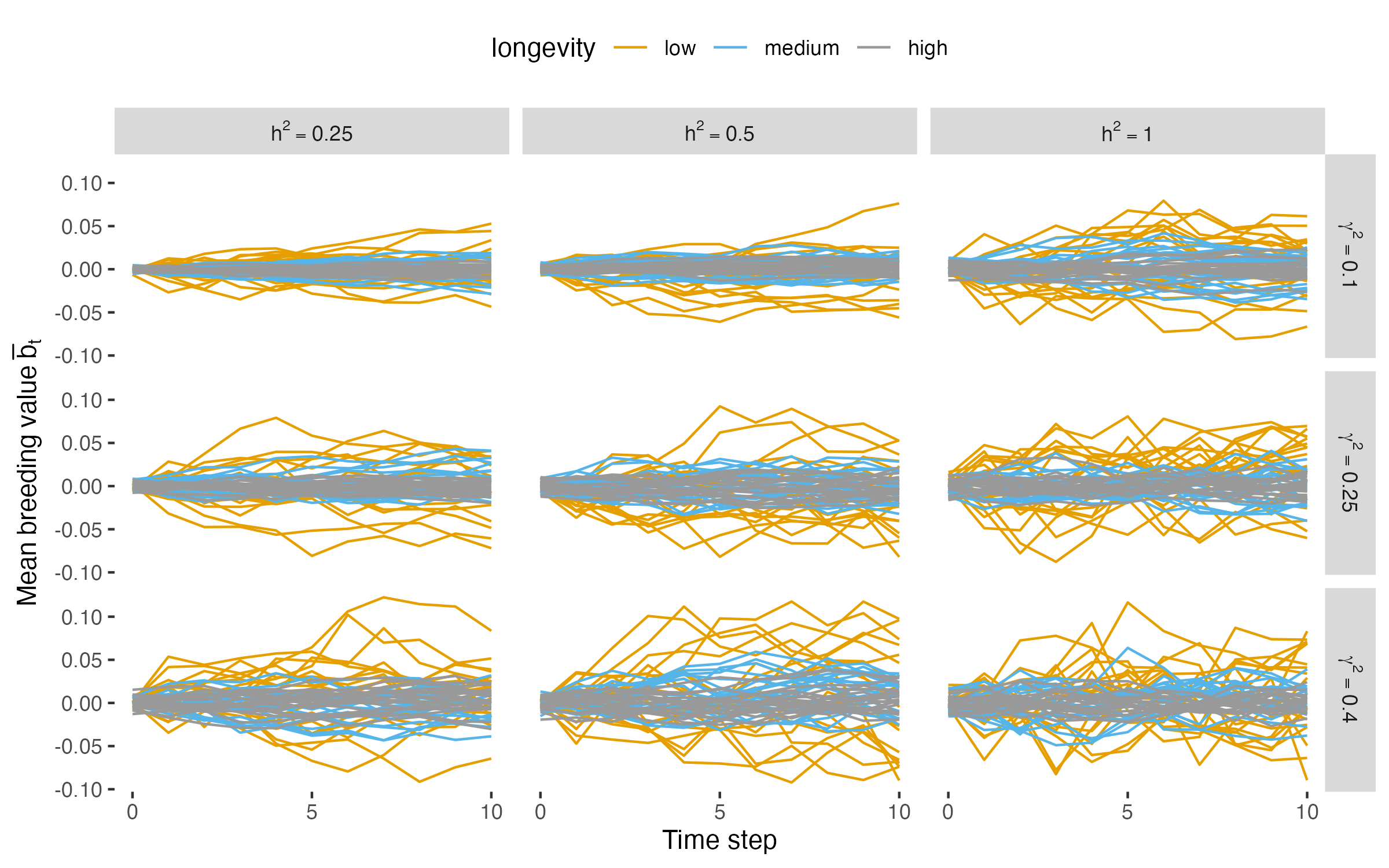

### fig_v5_ebar_steady.png

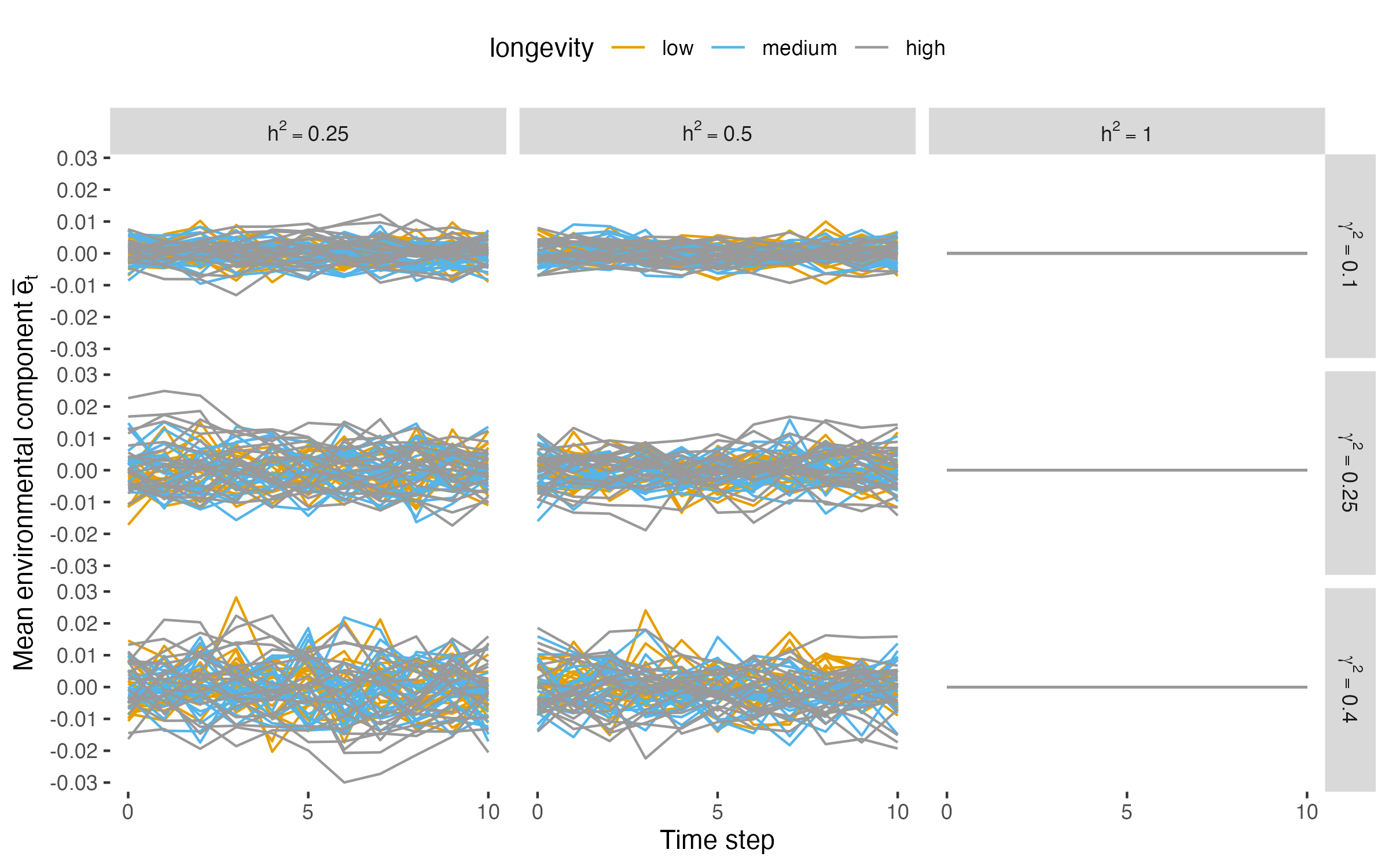

### fig_v6_zbar_steady.png

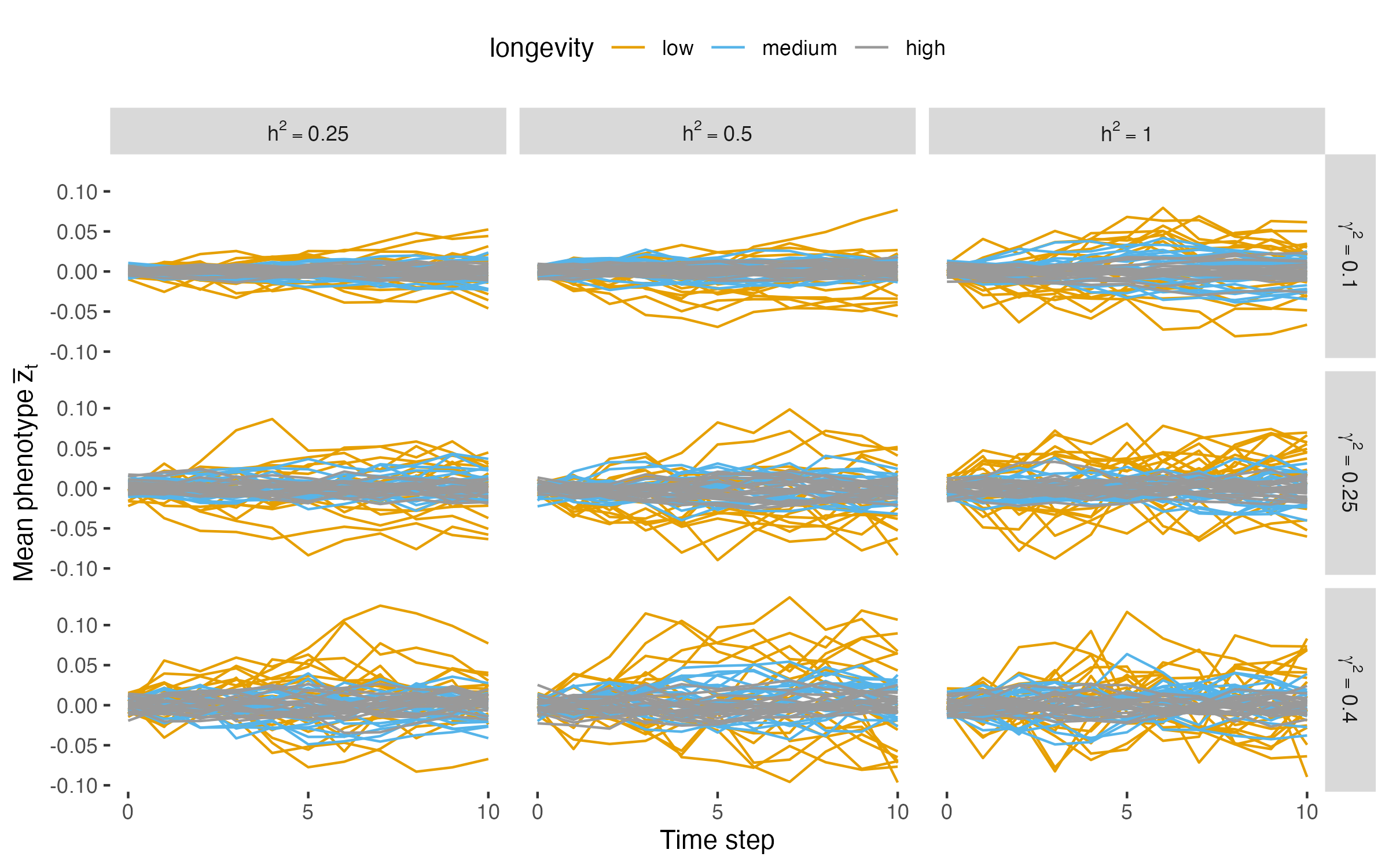

### fig_v7_bvar_age_steady.png

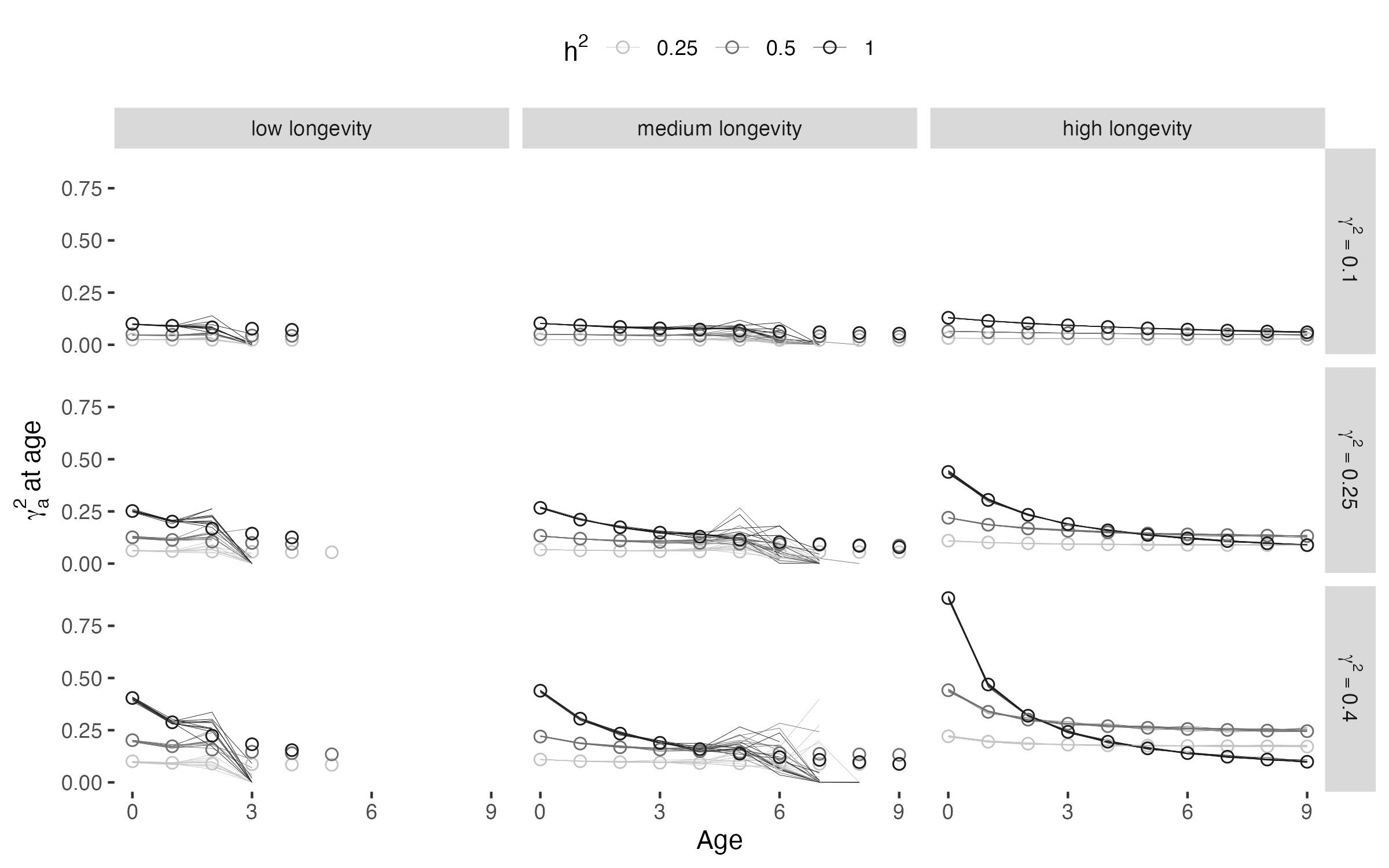

### fig_v8_evar_age_steady.png

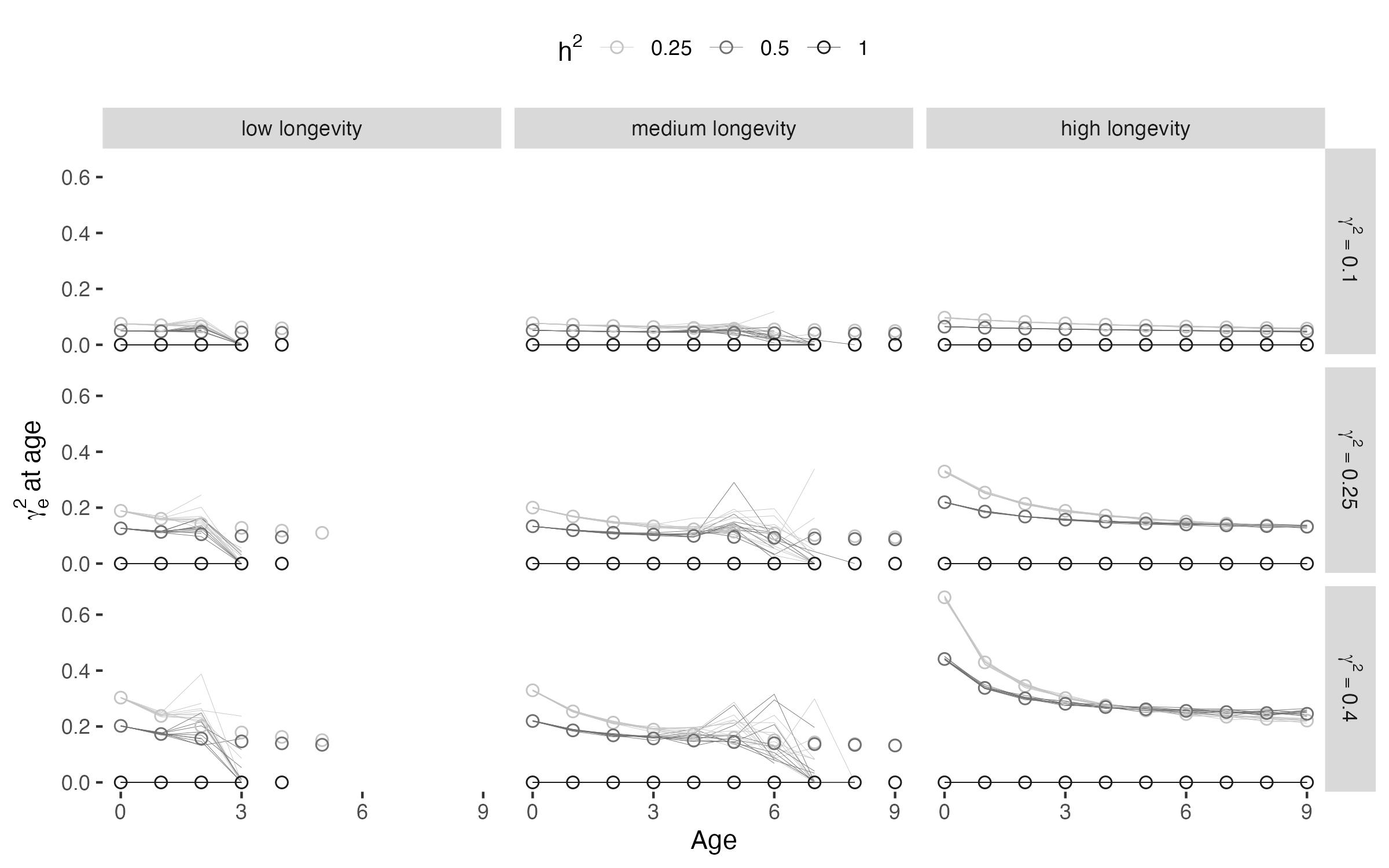

### fig_v9_zvar_age_steady.png

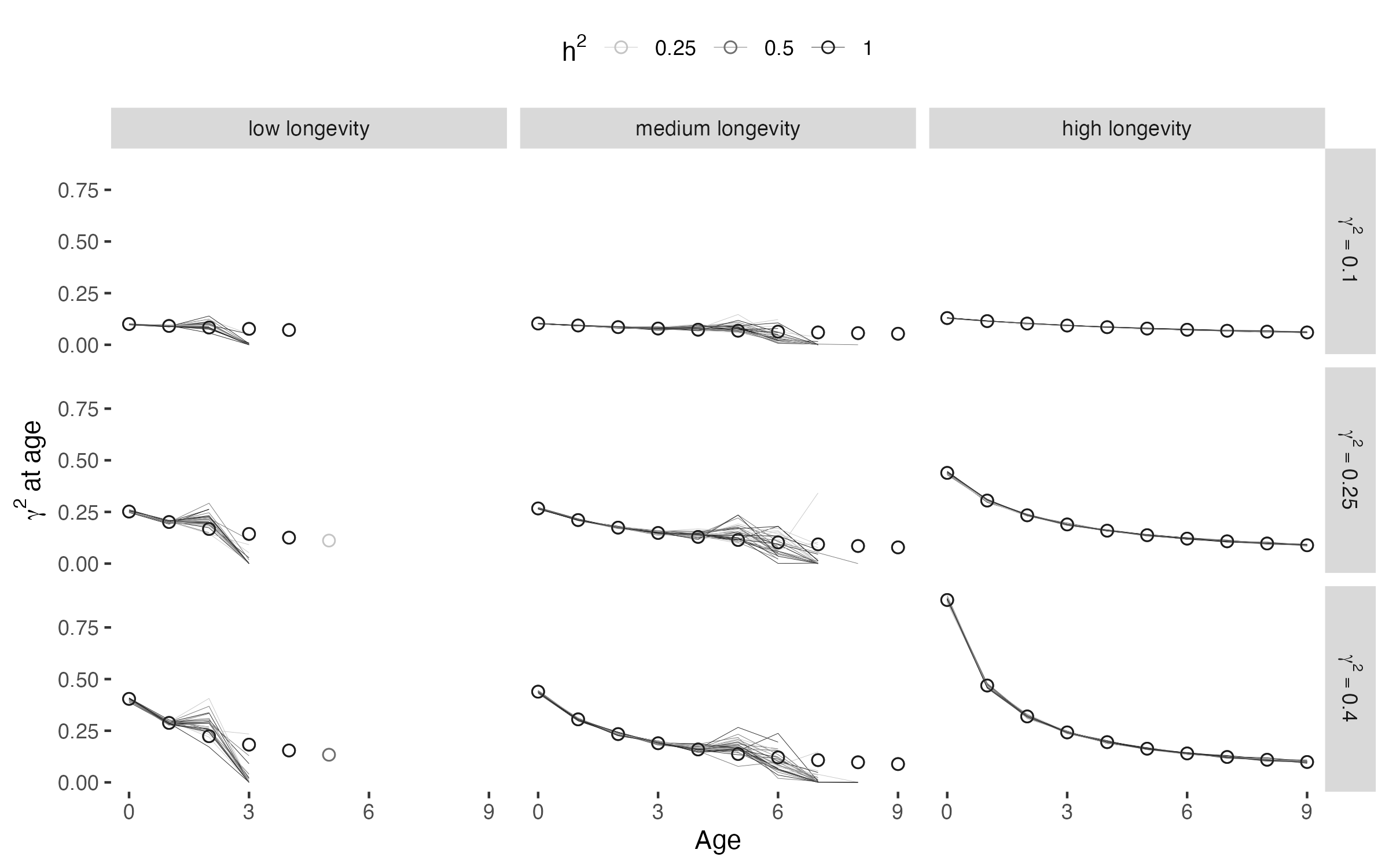

### fig_v10_rhok_age_steady.png

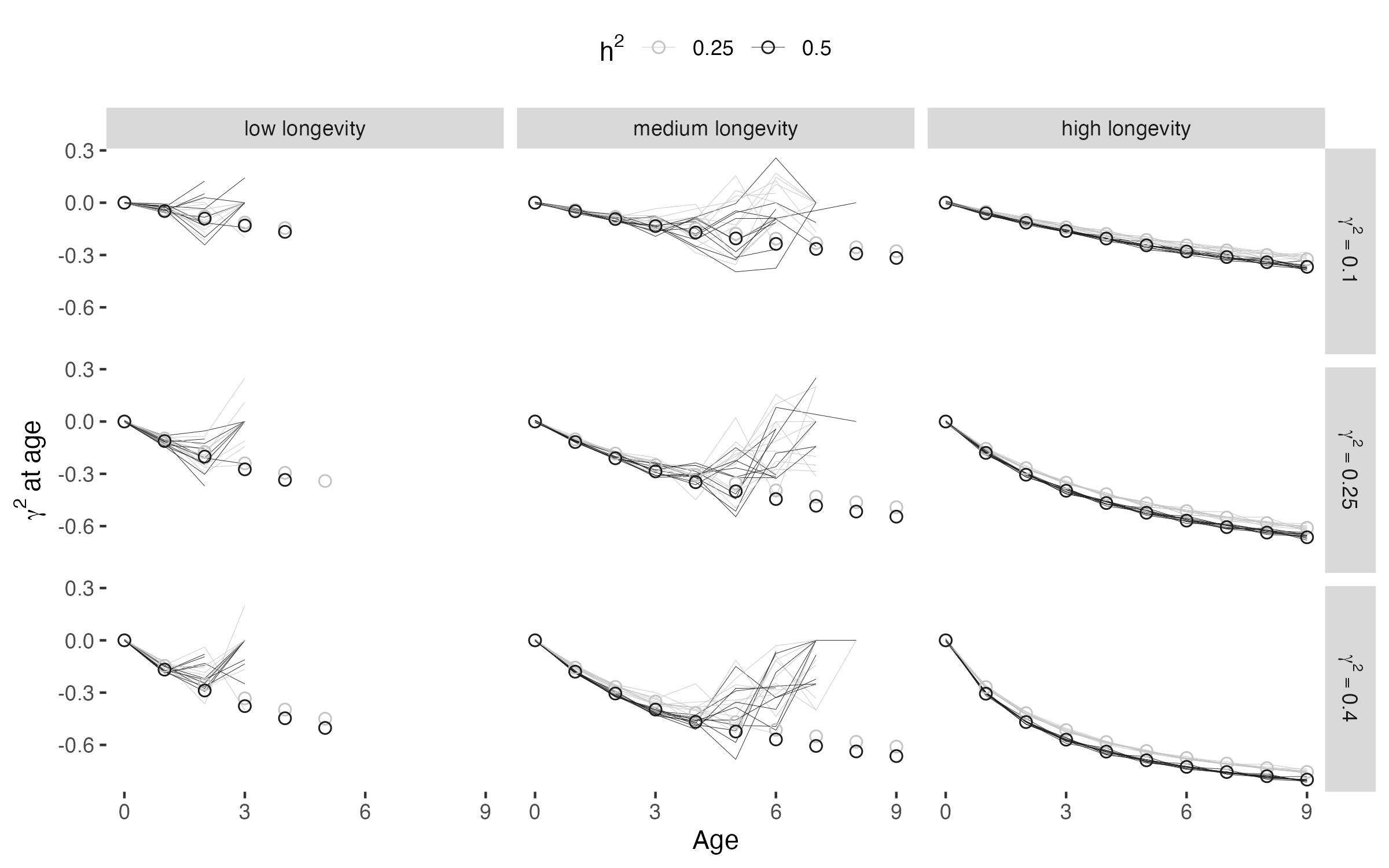

### fig_v11_bbar_change.png

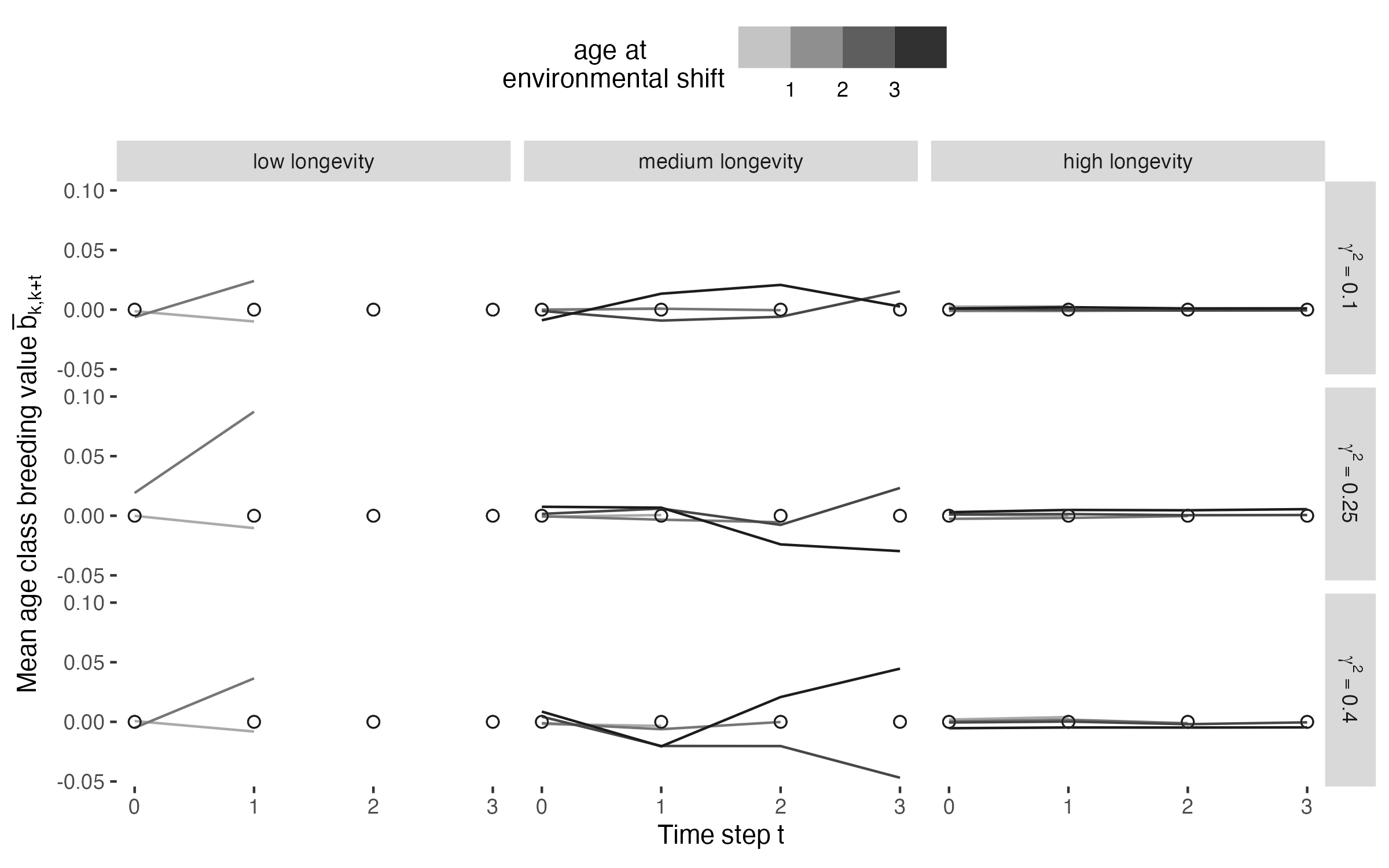

### fig_v12_ebar_change.png

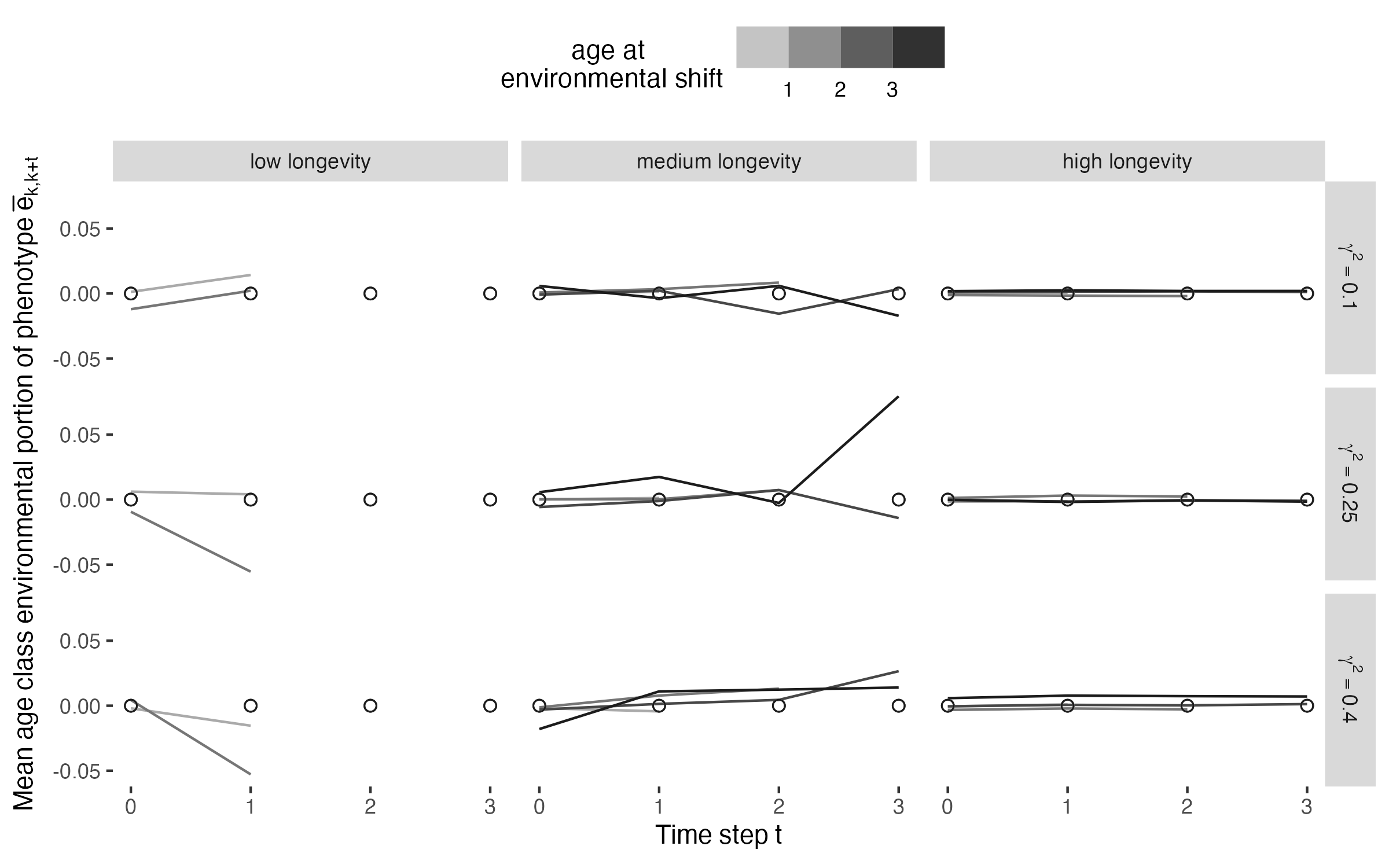

### fig_v13_zbar_change.png

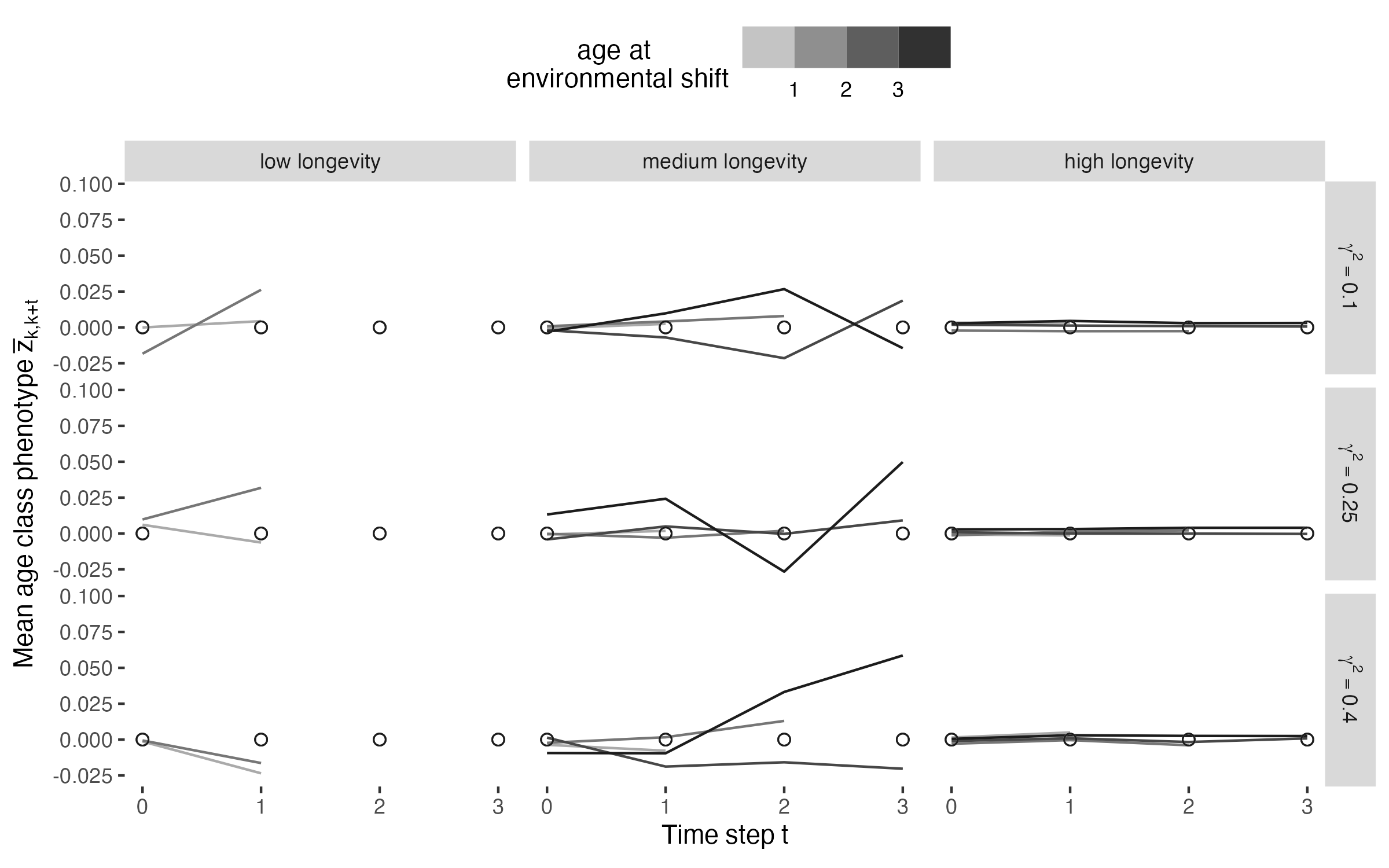

### fig_v14_bvar_change.png

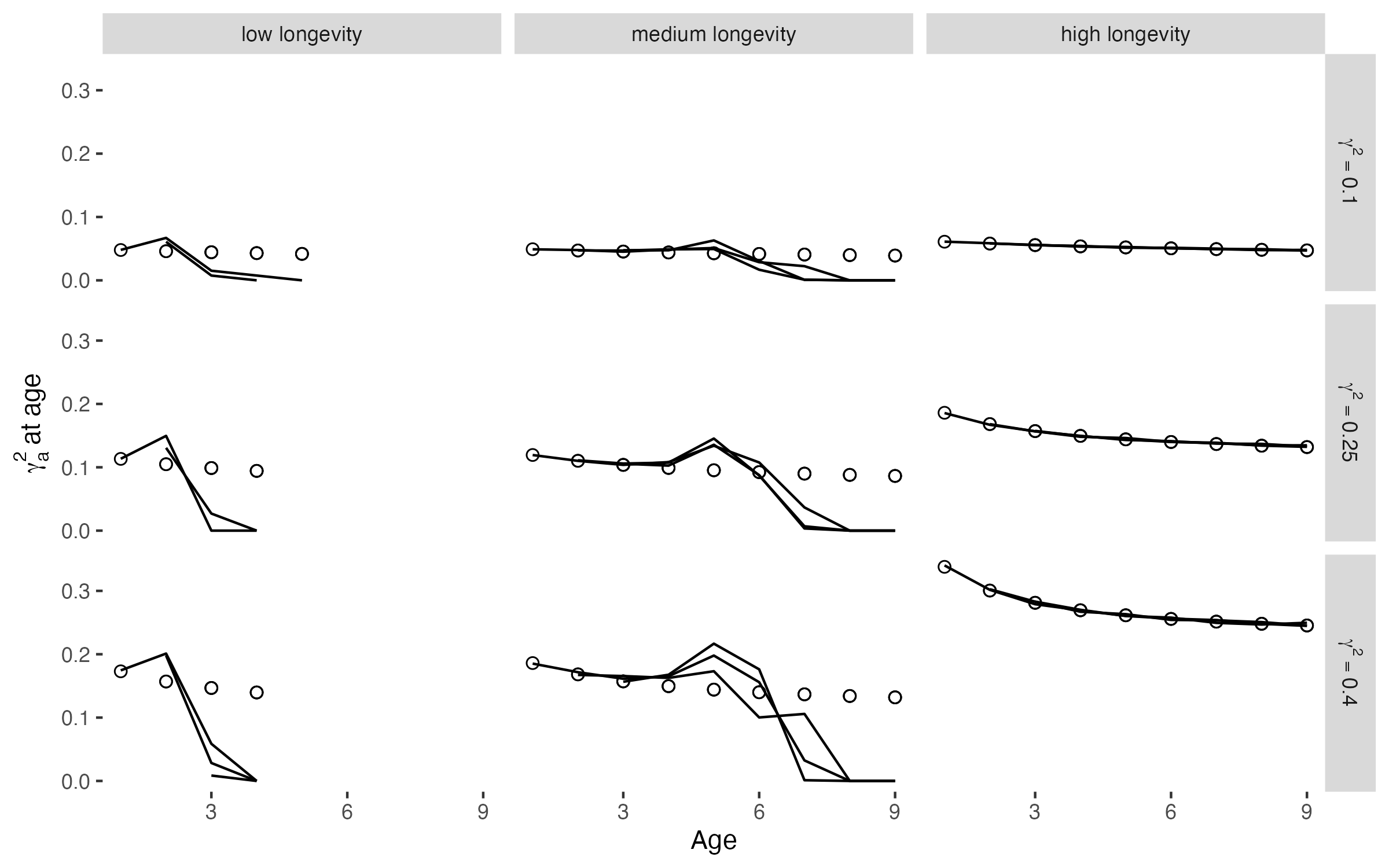

### fig_v15_evar_change.png

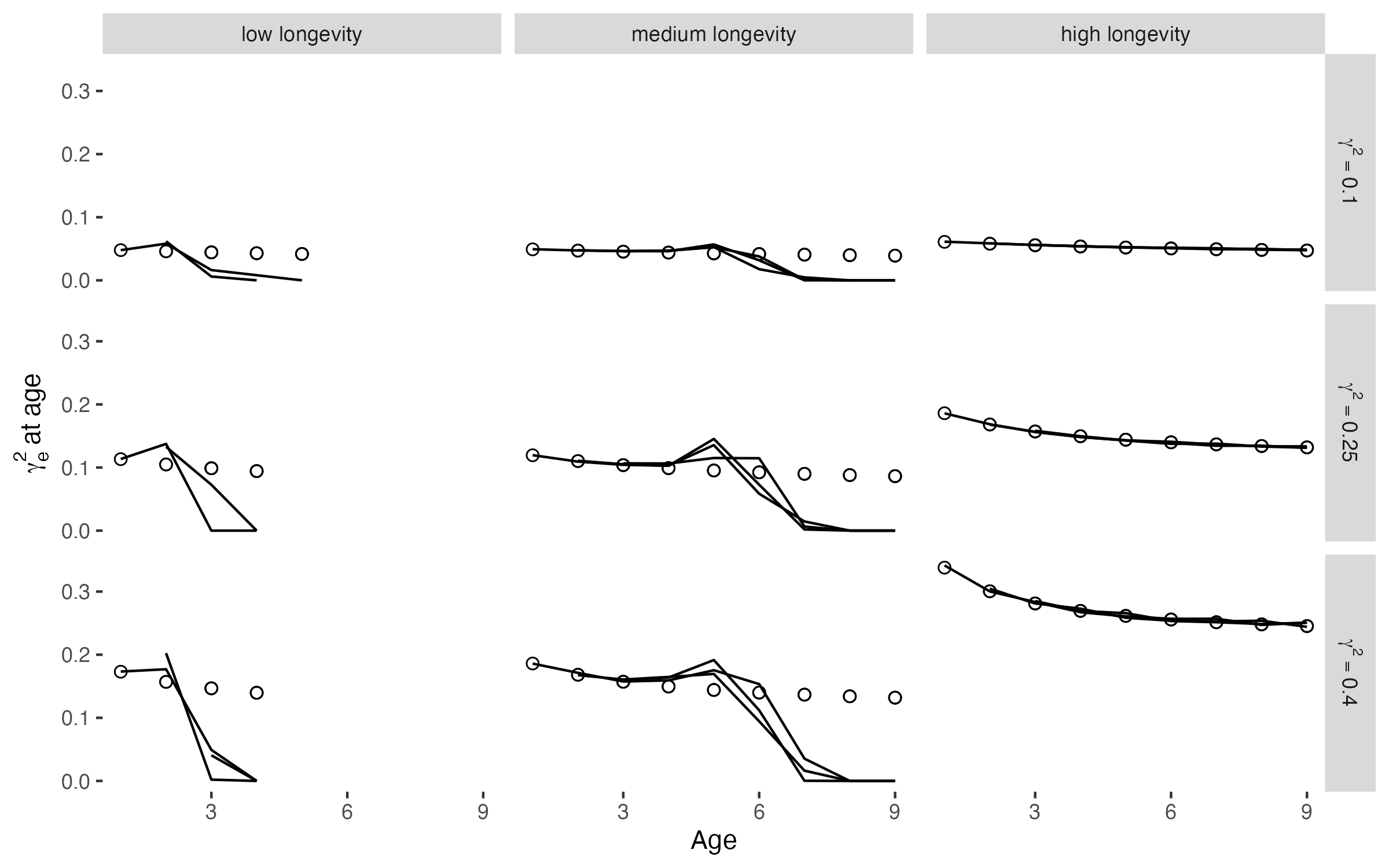

### fig_v16_zvar_change.png

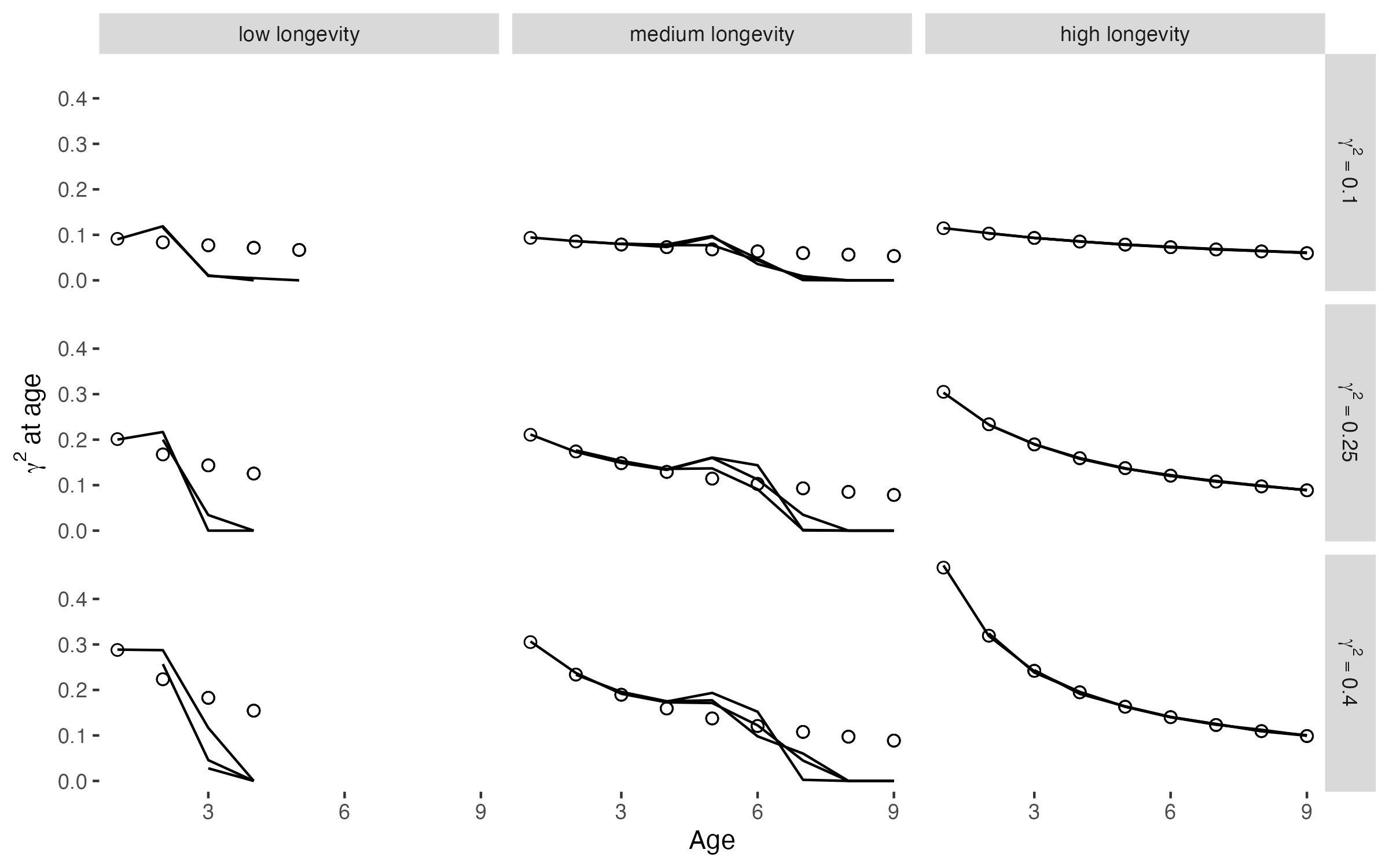

### fig_v17_rhok_change.png

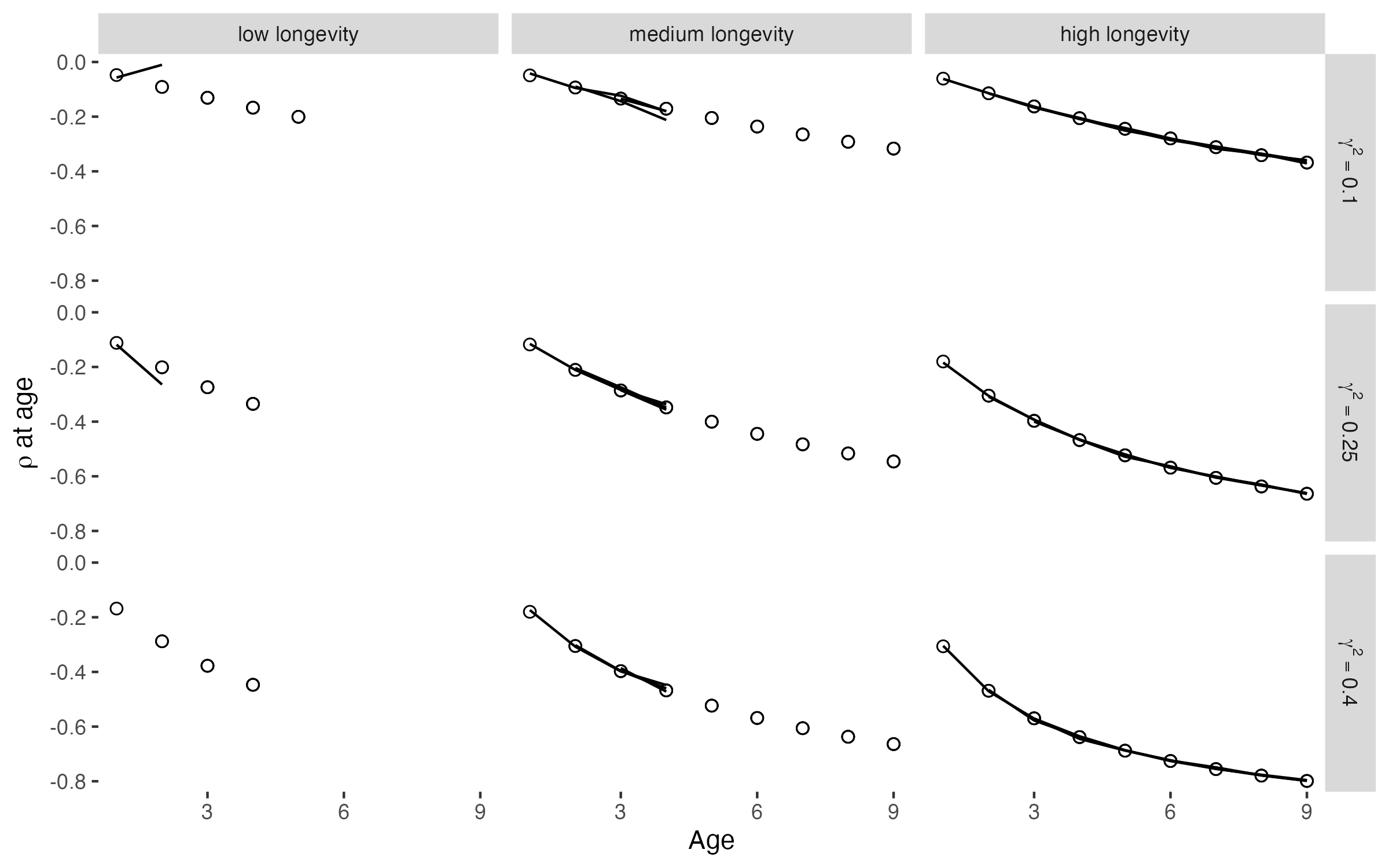

### figE1_popsize.png

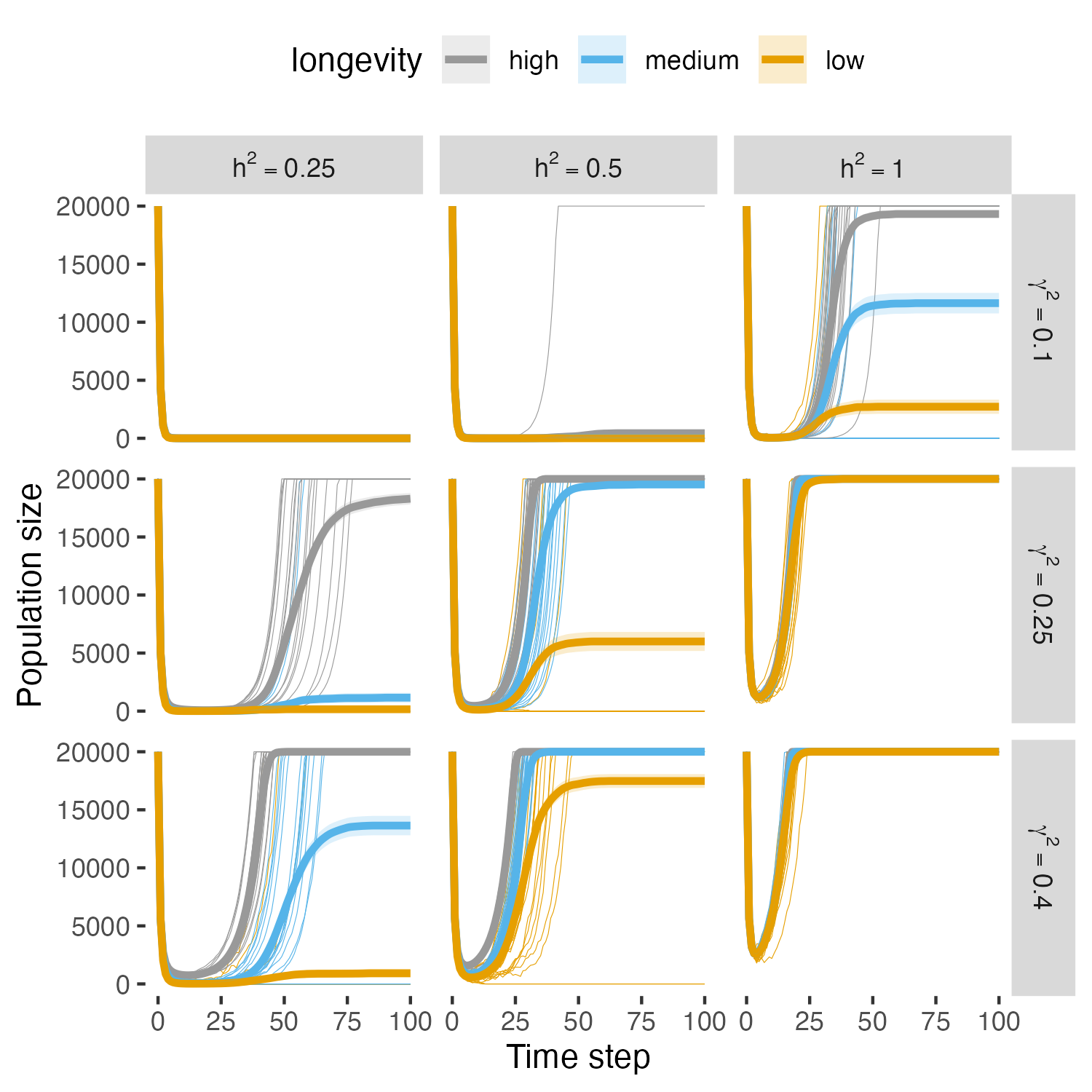

### figE2_extinctions.png

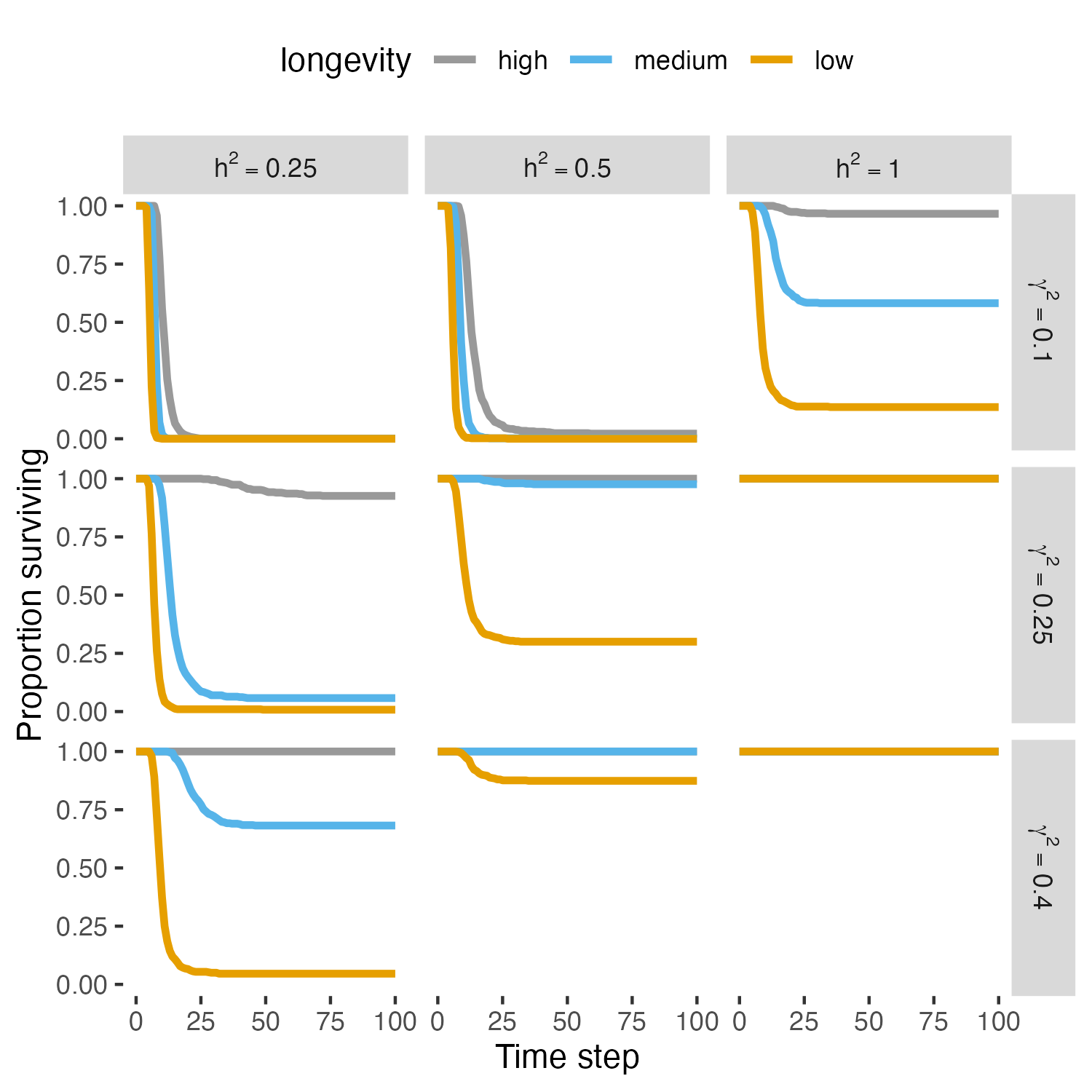

### figE3_pheno_comp_mean.png

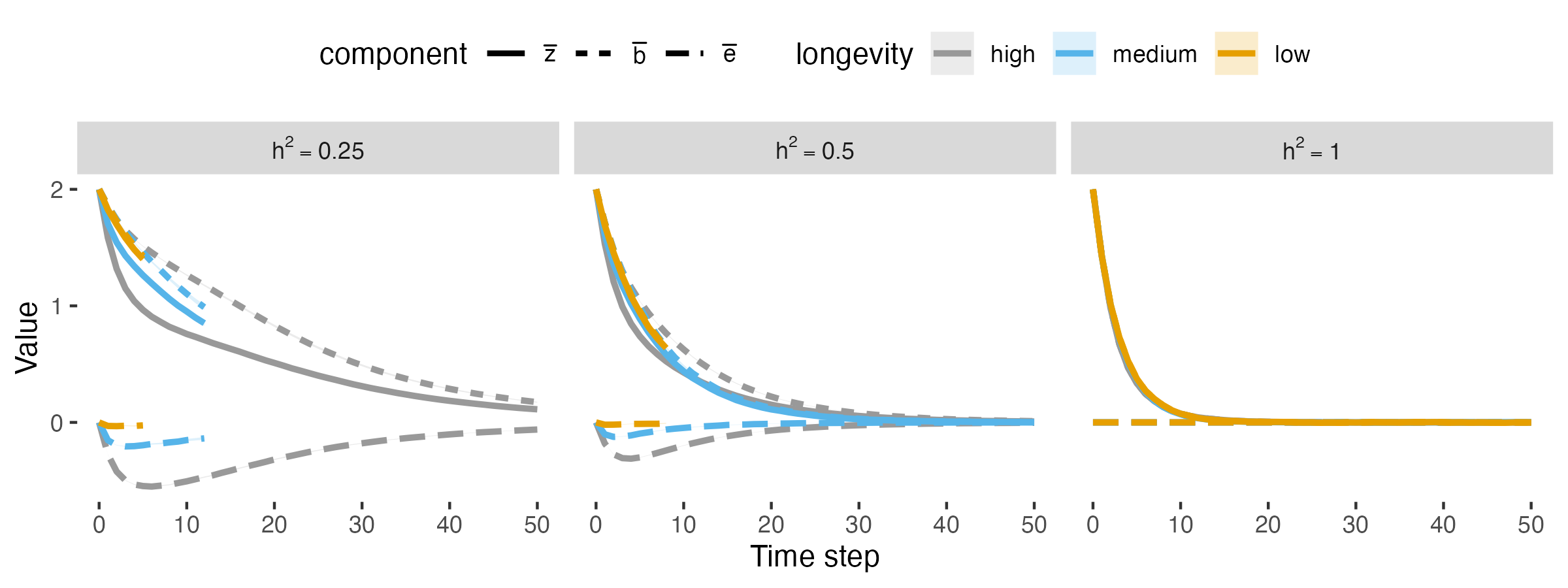

### figE4_pheno_comp_vars.png

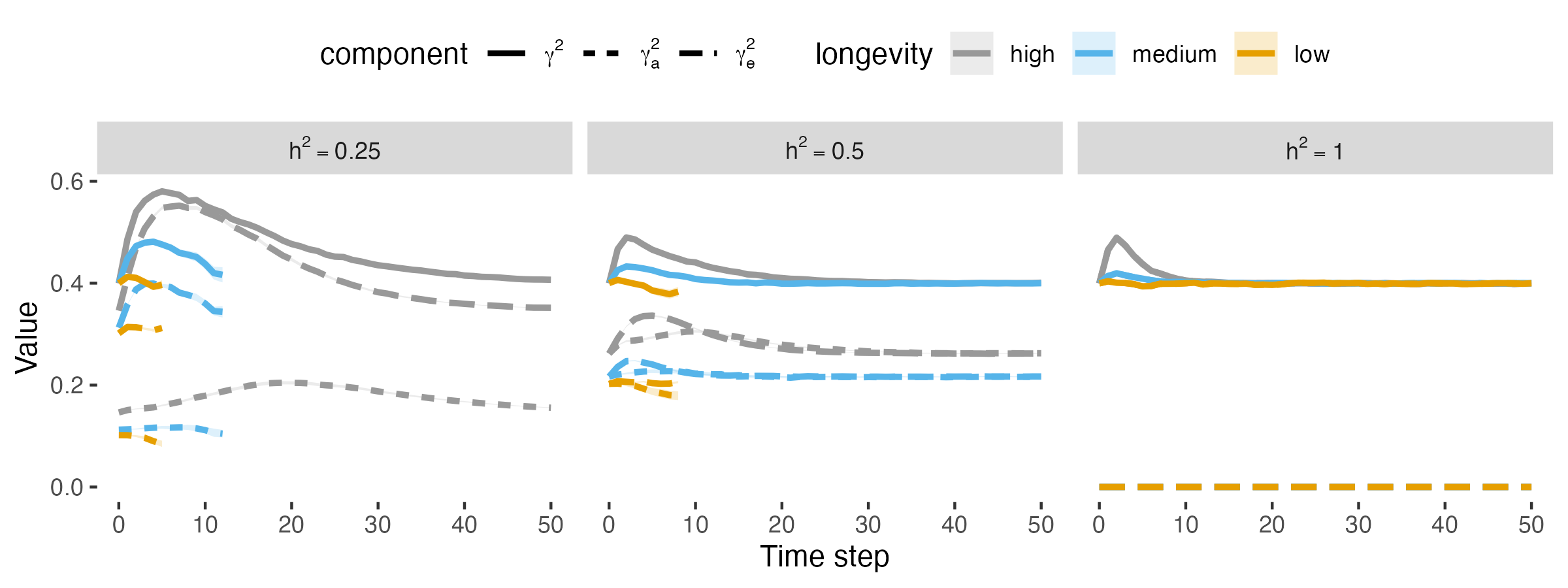

### figS1_age_distn.png

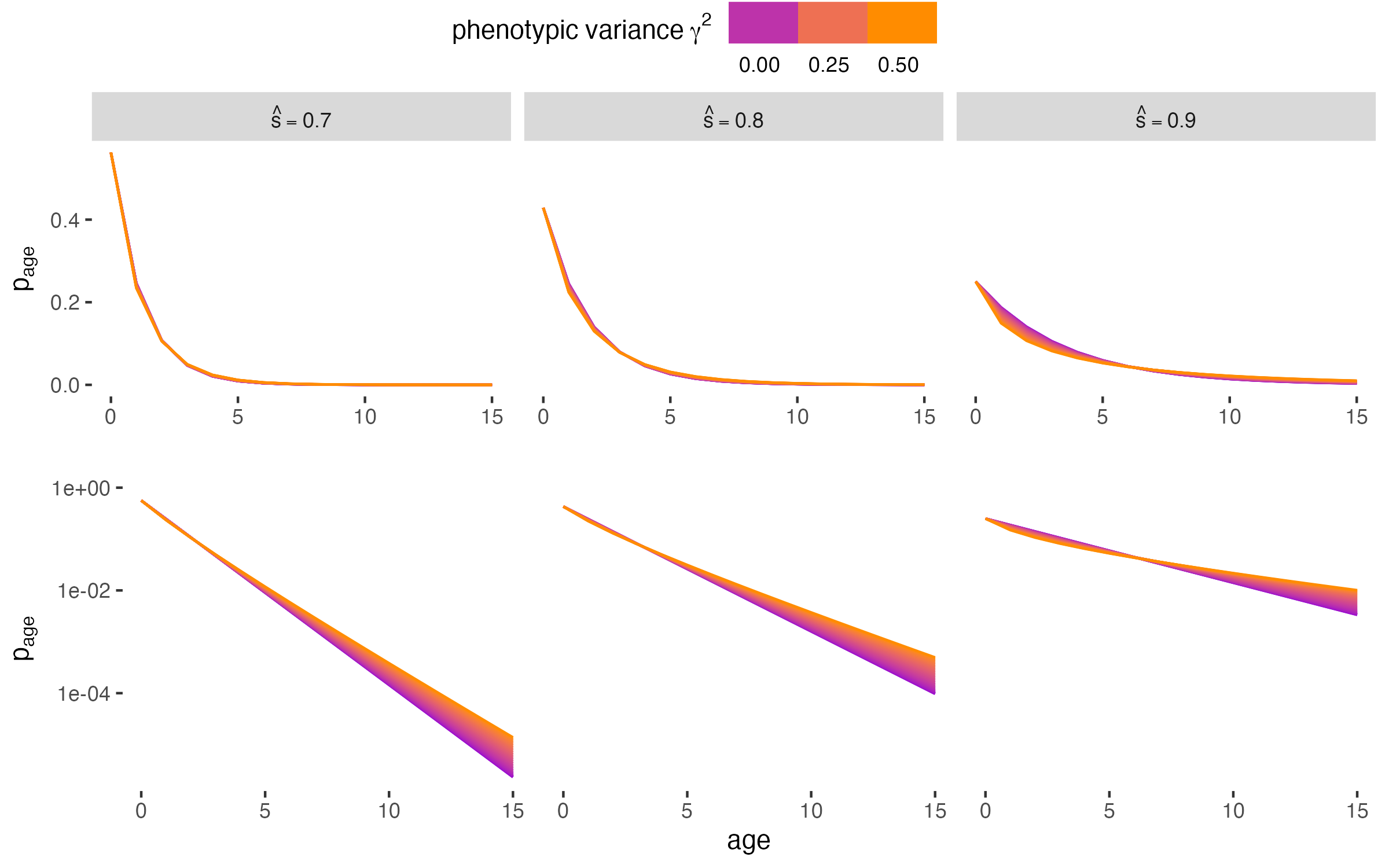

### figS3_pheno_comp_vars_postsel.png

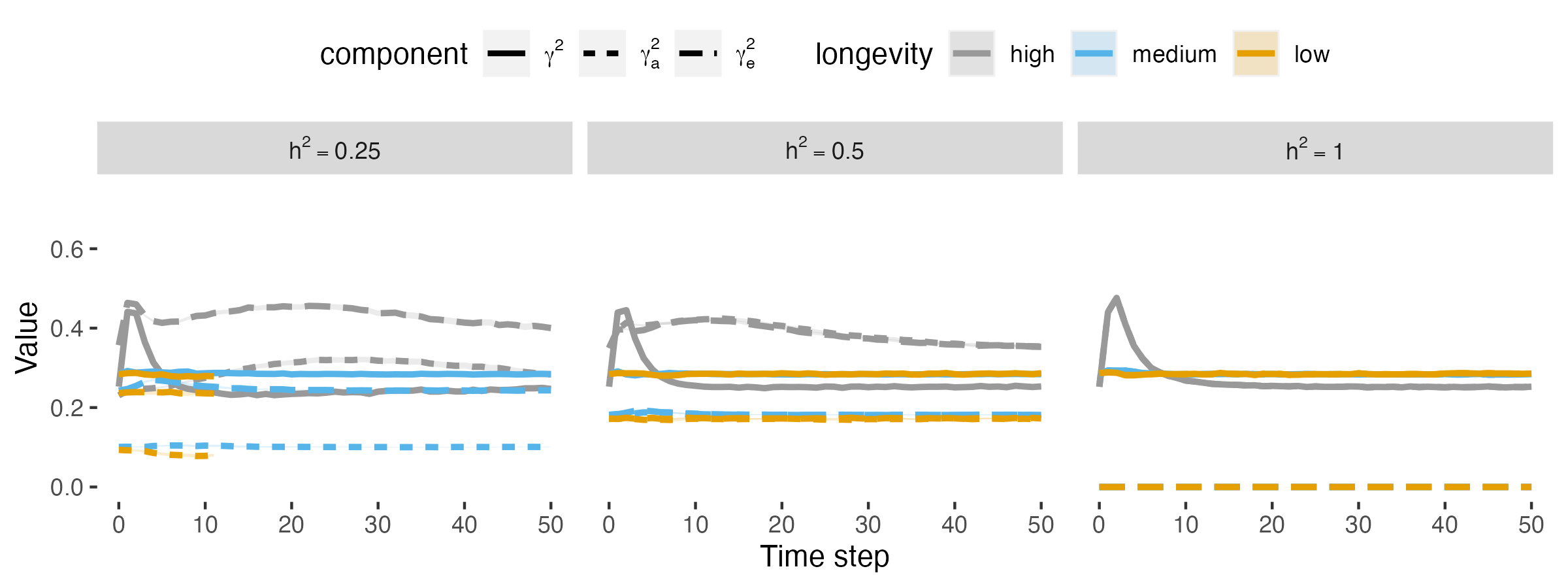

### figS19_generation_time_scaling.png

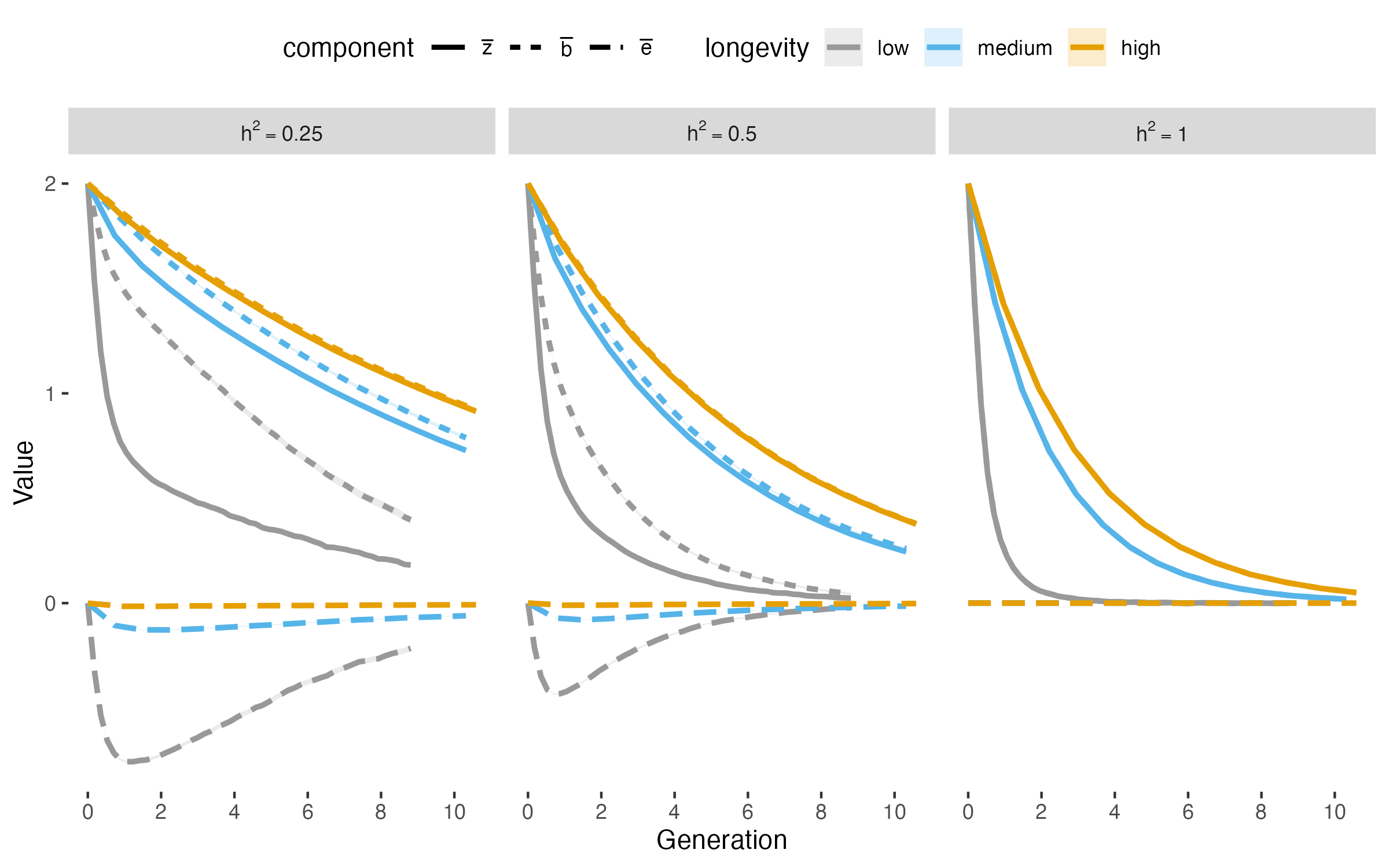

### figS20_agedist_medlowlong.png

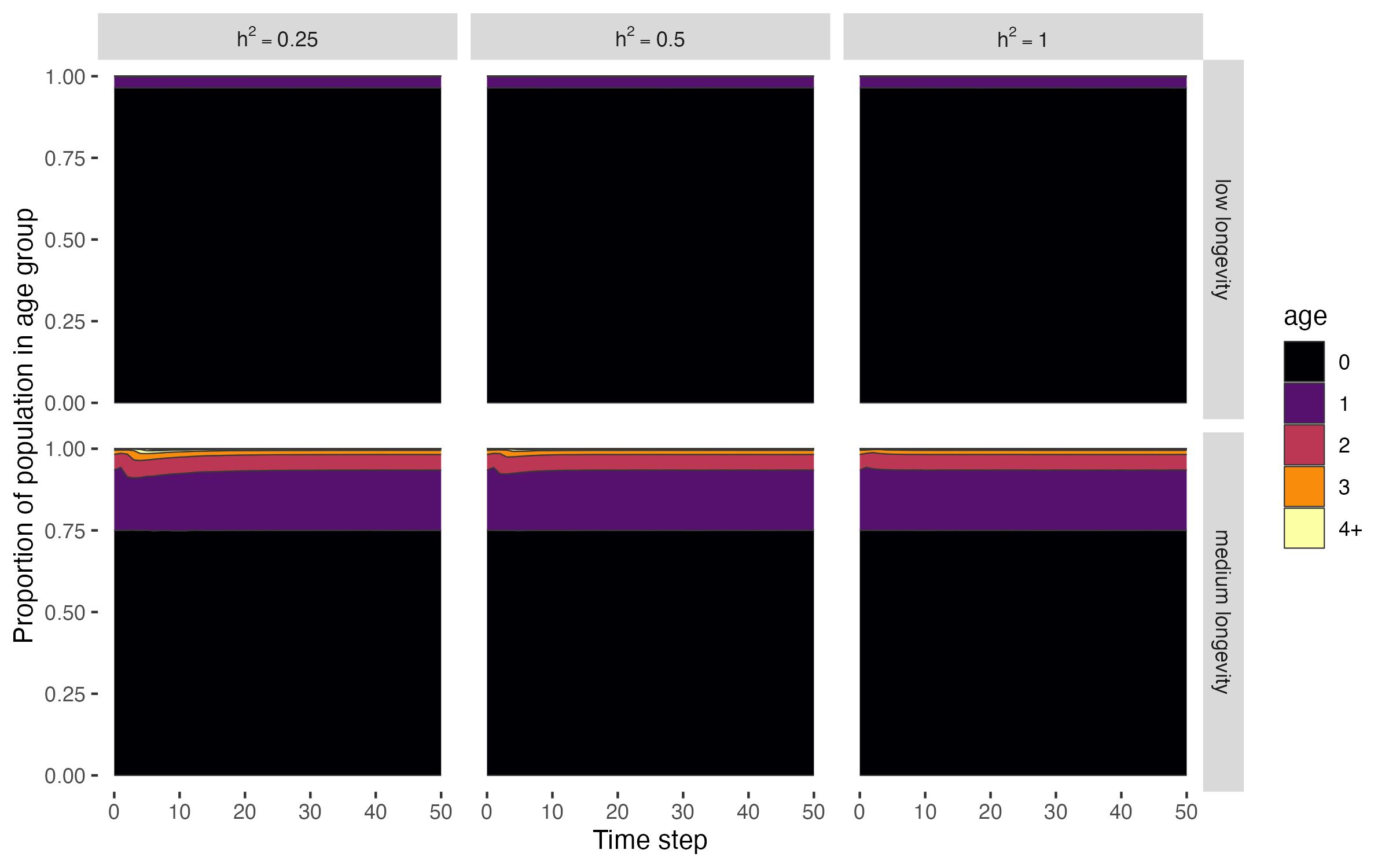

### figS21_generation_time_scaling.png

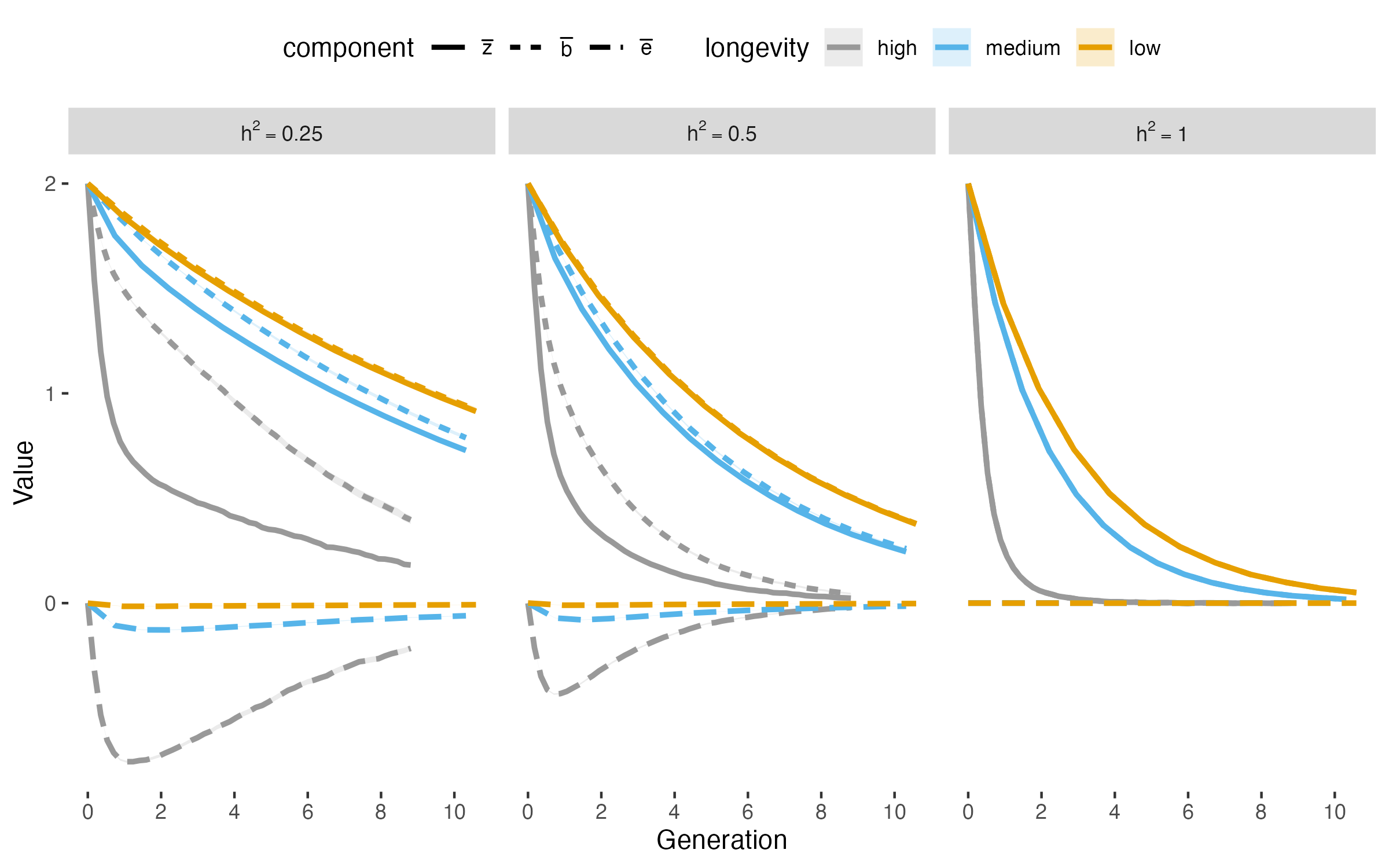

### figS22_lifetime_fitness.png

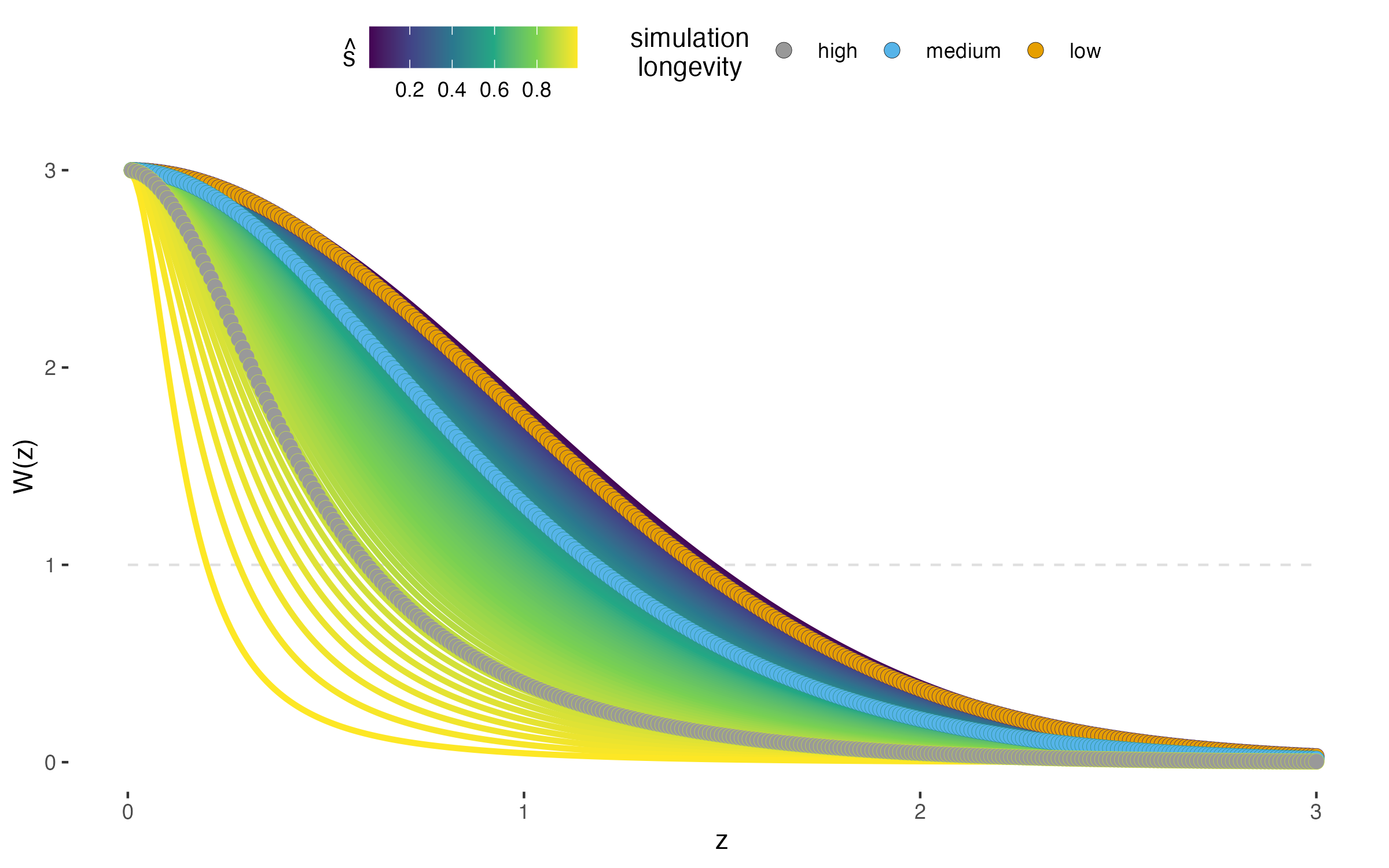
